# The class XIV myosin of *Toxoplasma gondii*, TgMyoA, is druggable in an animal model of infection

**DOI:** 10.1101/2022.09.09.507210

**Authors:** Anne Kelsen, Robyn S Kent, Anne K Snyder, Eddie Wehri, Stephen J Bishop, Rachel V Stadler, Cameron Powell, Bruno Martorelli di Genova, Pramod K Rompikuntal, Martin J Boulanger, David M Warshaw, Nicholas J Westwood, Julia Schaletzky, Gary E Ward

## Abstract

*Toxoplasma gondii* is a widespread apicomplexan parasite that can cause severe disease in its human hosts. The ability of *T. gondii* and other apicomplexan parasites to invade into, egress from, and move between cells of the hosts they infect is critical to parasite virulence and disease progression. An unusual and highly conserved parasite myosin motor (TgMyoA) plays a central role in *T. gondii* motility. The goal of this work was to test whether pharmacological inhibition of TgMyoA can alter disease progression in an animal model of infection. To this end, we sought to identify small molecule inhibitors of TgMyoA by screening a collection of 50,000 structurally diverse small molecules for inhibitors of the recombinant motors actin-activated ATPase activity. The top hit to emerge from the screen, KNX-002, inhibited TgMyoA with little to no effect on any of the vertebrate myosins tested. KNX-002 was also active against parasites, inhibiting parasite motility and growth in culture in a dose-dependent manner. We used chemical mutagenesis, selection in KNX-002, and targeted sequencing to identify a mutation in TgMyoA (T130A) that renders the recombinant motor less sensitive to compound. Compared to wild-type parasites, parasites expressing the T130A mutation showed reduced sensitivity to KNX-002 in motility and growth assays, confirming TgMyoA as a biologically relevant target of KNX-002. Finally, KNX-002 was shown to slow disease progression in mice infected with wild-type parasites, but not parasites expressing the resistance-conferring TgMyoA T130A mutation. These data demonstrate the specificity of KNX-002 for TgMyoA, both *in vitro* and *in vivo*, and validate TgMyoA as a druggable target for toxoplasmosis. Since TgMyoA is essential for virulence, conserved in apicomplexan parasites, and distinctly different from the myosins found in humans, pharmacological inhibition of MyoA offers a promising new approach to treating the devastating diseases caused by *T. gondii* and other apicomplexan parasites.

## Introduction

Nearly one third of the world’s population are or have been infected with the apicomplexan parasite, *Toxoplasma gondii*. Although most infections are subclinical, acute toxoplasmosis can have severe consequences in neonates and immunocompromised individuals. Congenital infection can lead to spontaneous abortion or stillbirth, and even children born with subclinical infection frequently experience sequelae later in life including neurological damage and vision impairment [1, 2]. Among the immunocompromised, toxoplasmic encephalitis is a particularly significant risk in AIDS patients who are unaware of their HIV status [3-5] and in the ∼50% of HIV-infected individuals worldwide who do not have access to antiretroviral therapy [6, 7]. In individuals with AIDS, adverse effects of the currently available drugs to suppress toxoplasmosis [8] cause the discontinuation of treatment in up to 40% of patients [4, 9-11], highlighting the need for new, better-tolerated drugs to reduce toxoplasmosis-related morbidity and mortality.

Within the brain and other tissues of a patient suffering from active toxoplasmosis, the tachyzoite stage of the parasite uses a unique form of substrate-dependent “gliding” motility to invade into and egress from the cells of its host, to spread throughout the patient’s tissues, and to migrate across biological barriers [12-15]. *T. gondii* tachyzoites move in the extracellular matrix along flattened corkscrew-like trajectories, with regularly repeating periods of acceleration and deceleration [16]. The mechanisms underlying gliding motility are thought to be conserved among apicomplexan parasites, including *Plasmodium spp*. (the causative agents of malaria) and *Cryptosporidium spp*. (which cause severe diarrheal disease). Motility is driven, at least in part, by an unusual class XIVa myosin motor protein, myosinA (MyoA), found only in apicomplexan parasites and a few ciliates. In *T. gondii*, this motor consists of the TgMyoA heavy chain and two associated light chains, TgMLC1 and either TgELC1 or TgELC2 [17-19]. TgMyoA is a single-headed motor with a very low duty ratio (*i*.*e*., it remains strongly bound to actin for only ∼1% of its catalytic cycle) yet moves actin filaments at the same speed as skeletal muscle myosin, a double-headed motor with a 5-fold higher duty ratio [20, 21]. TgMyoA lacks a conventional tail, as well as several residues conserved in other myosins that normally function to regulate actomyosin activity [22, 23], and employs atypical strategies for chemomechanical coupling and force transduction [24, 25].

Depletion of *TgMyoA* results in severely reduced parasite motility, host cell invasion, and host cell egress [26-29]. As a result, parasites lacking TgMyoA are avirulent in an animal model of infection [28]. Despite its importance in the *T. gondii* life cycle, we know little about how TgMyoA generates the complex pattern of parasite motility seen in 3D or how TgMyoA function contributes to disease pathogenesis in an infected host. Because the parasite can compensate for the loss or reduced expression of proteins important to its life cycle [30-32], small-molecule inhibitors of TgMyoA would serve as valuable complementary tools for determining how different aspects of motor function contribute to parasite motility and the role played by TgMyoA in parasite dissemination and virulence. Specific inhibitors are also necessary to establish TgMyoA as a “druggable” target, *i*.*e*., a protein whose activity is amenable to inhibition by small molecules, and to test whether pharmacological inhibition of TgMyoA activity *in vivo* alters the course of an infection.

We describe here the identification and characterization of KNX-002, the first specific inhibitor of an apicomplexan Class XIVa myosin. KNX-002 inhibits TgMyoA motor activity, parasite motility, and parasite growth in culture in a dose-dependent manner, with little or no effect on a variety of other vertebrate myosins. On-target activity in parasites was confirmed by identifying a mutation in TgMyoA that rescues parasite motility and growth in the presence of the compound. The availability of a specific inhibitor of TgMyoA, together with isogenic parasite strains that show differential sensitivity to the compound, enabled us to undertake the first direct test of whether pharmacological inhibition of MyoA can be used to alter the progression of disease caused by an apicomplexan parasite.

## Results

### High-throughput screening identifies a novel inhibitor of TgMyoA

To identify inhibitors of TgMyoA motor activity, we used our previously described method for producing large amounts of functional motor, in which the TgMyoA heavy chain is co-expressed in insect cells with its two light chains, TgMLC1 and TgELC1, and a myosin co-chaperone protein [33]. We developed a miniaturized coupled enzyme assay to measure actin-dependent ATPase activity of the purified recombinant motor and screened 50,000 compounds from the compound library at Cytokinetics, Inc., for inhibitors of this ATPase activity. Hit follow-up and characterization included dose-response analysis using resupplied compound that was determined to be >95% pure by LC/MS analysis and control assays demonstrating the compound was inactive against an unrelated ATPase, hexokinase. The most potent hit was 1-(4-methoxyphenyl)-N-((3-(thiophen-2-yl)-1H-pyrazol-4-yl)methyl)cyclopropan-1-amine (Figure 1A, inset), subsequently named KNX-002. KNX-002 inhibits TgMyoA ATPase activity in a dose-dependent manner with an IC50 of 2.8 μM (95% C.I. 2.4-3.2 μM; Figure 1A). In contrast, KNX-002 has little to no detectable effect on the actin-activated ATPase activity of bovine cardiac myosin, chicken gizzard smooth muscle myosin, rabbit fast skeletal muscle myosin, or bovine slow skeletal muscle myosin, up to the highest concentrations tested (40 μM; Figure 1B and Suppl. Figure 1). These data identify KNX-002 as a potentially specific inhibitor of *T. gondii* class XIVa myosin activity.

**Figure 1:**
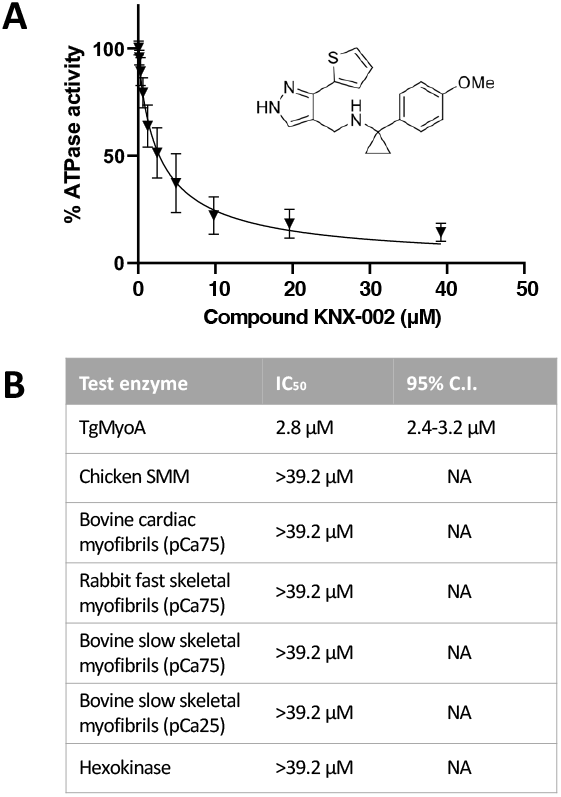
KNX-002 inhibits the actin-activated ATPase activity of TgMyoA but not vertebrate myosins. **(A)** The effect of various concentrations of KNX-002 on TgMyoA ATPase activity, normalized to the activity with an equivalent amount of DMSO (vehicle) only. Inset shows the structure of KNX-002. Each data point represents mean ATPase activity of three different batches of KNX-002 ± SEM. **(B)** The inhibitory effect of KNX-002 on TgMyoA and various vertebrate myosins (IC_50_ and 95% confidence interval [C.I.]). The dose-response curves for the individual vertebrate myosins are shown in Suppl. Figure 1.

### KNX-002 inhibits the growth of *T. gondii* in cell culture, with no detectable toxicity to human cells

The three lipid bilayers that comprise the pellicle of *T. gondii, i*.*e*., the plasma membrane and subjacent double bilayer of the inner membrane complex [34, 35], represent a potential barrier to the penetration of externally applied drugs. To determine if KNX-002 can access and inhibit the motor in live parasites, we determined the compound’s effect on parasite expansion in culture in a modified plate-based growth assay (see Methods and ref. [36]). KNX-002 does indeed inhibit parasite growth, in a dose-dependent manner (Figure 2A), with an IC50 of 16.2 μM (95% confidence interval [C.I.] 13.0 - 20.5 μM; Figure 2B).

**Figure 2:**
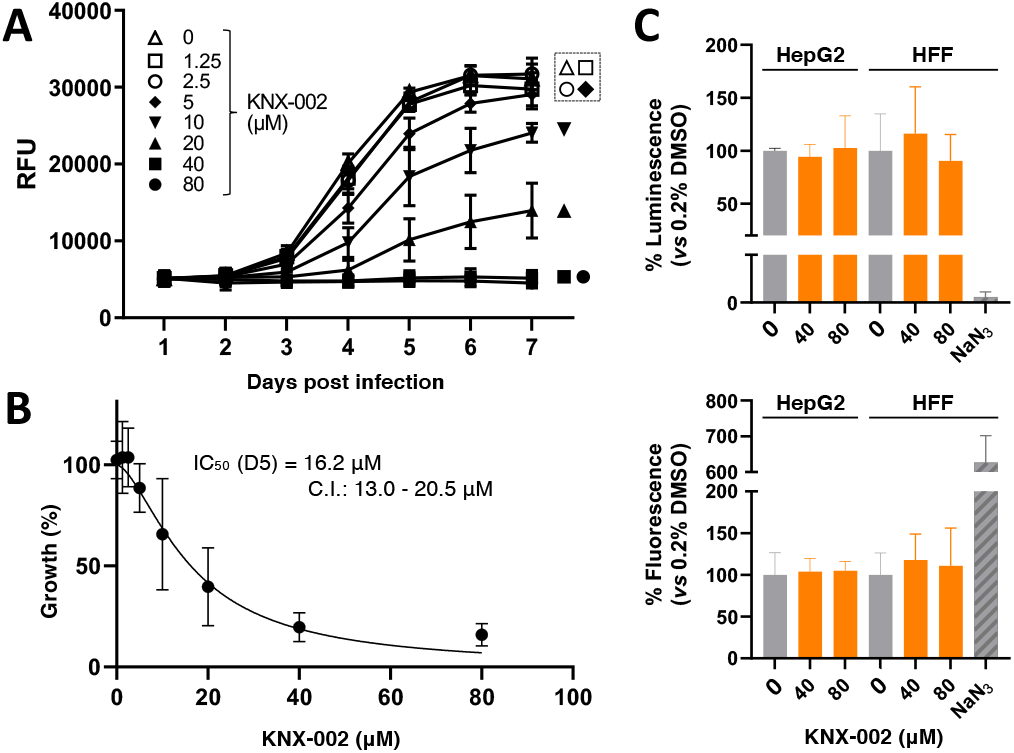
KNX-002 inhibits the growth of *T. gondii* in culture, with no detectable host cell toxicity. **(A)** tdTomato-expressing parasites were preincubated with various concentrations of KNX-002 for 5 minutes (0 = DMSO vehicle only) and then added to HFF cells on a 384-well plate. Fluorescence was measured daily to quantify parasite growth, in the continued presence of compound, over the next 7 days. RFU = relative fluorescence units. The data shown are the mean ± SEM from two independent biological replicates, each consisting of 3 technical replicates at all time points. **(B)** The data from day 5 (D5) of the growth assay shown in panel (A) were used to calculate the IC_50_ of KNX-002 for parasite growth (16.2 μM, 95% C.I. = 13.0 to 20.5). See Figure 5A for a repeat of these growth assays and IC_50_ calculations. **(C)** Non-confluent HepG2 and HFF cells were cultured in the presence of 0 (DMSO vehicle only), 40, or 80μM KNX-002 for 72 hours. Cell viability/growth was measured using Promega CellTiter-Glo® (top panel) and toxicity was measured using Promega CellTox™ green (bottom panel). 0.1% w/v NaN_3_ served as a positive control for cytotoxicity / growth inhibition. Results are plotted relative to the DMSO controls. Bars show the mean of three independent experiments ± SEM. KNX-002 shows no evidence of cytotoxicity or growth inhibition in either cell type at either 40 or 80μM.

Parasite growth in culture involves repeated lytic cycles comprised of host cell invasion, intracellular replication, lysis of and egress from the infected cell, and migration to new host cells. While invasion, egress and migration are all TgMyoA-dependent processes, intracellular replication does not require TgMyoA [26, 28]. To test whether any portion of the observed growth inhibition in culture is due to off-target effect(s) of KNX-002 on parasite replication, we scored the number of parasites per parasitophorous vacuole over time during the lytic cycle. The compound had no detectable effect on parasite replication in this assay (Suppl. Figure 2).

To exclude the possibility that the effect of KNX-002 on parasite growth was due to a deleterious effect on the host cells, we also tested the effect of the compound on human foreskin fibroblasts (the host cells used in the growth assays) and human HepG2 liver cells (sensitive indicators of cytotoxicity and genotoxicity [37]) in 72-hour multiplexed cytotoxicity and cellular viability assays. KNX-002 showed no detectable deleterious effect on either cell line in either assay at the highest concentration tested (80 μM; Figure 2C).

### Structure-activity relationship (SAR) analysis

Next, we used the ATPase and parasite growth assays to carry out a preliminary SAR study, with the goal of determining which parts of KNX-002 are important for its biological activity. We began by modifying the *p*-OMe substituted aryl ring in KNX-002 (colored red in KNX-002 structure, Figure 3). Replacement of this structural unit in full by a hydrogen atom (in **VEST1**) confirmed the importance of this part of the molecule for biological activity (Figure 3 and Suppl. Figures 3 and 4). In terms of the preferred substituent on this aryl ring, inhibition of both *in vitro* ATPase activity and parasite growth were retained when the *p*-OMe substituent in KNX-002 was replaced by *p*-Cl (**VEST2**), and partially retained in analogs containing *p*-Me (**VEST3**) or *m,p*-dichloro (**VEST4**) substituents. **VEST5**, an analog in which the *p*-OMe in KNX-002 was replaced by a hydrogen, was significantly less active than KNX-002 in the purified protein assay but a strong inhibitor of parasite growth. **VEST5** was not toxic to host cells, extracellular parasites, or intracellular parasites (Suppl. Figures 5 and 6); further studies will be required to understand the activity of this analog in the parasite growth assay. The mild to moderate toxicity levels shown by **VEST2** (*p*-Cl) against HepG2 and HFF cells, compared to the lack of toxicity in either cell line demonstrated by KNX-002, led to the preliminary conclusion that the *p*-OMe aryl substituent present in the starting KNX-002 analogue was preferred.

**Figure 3:**
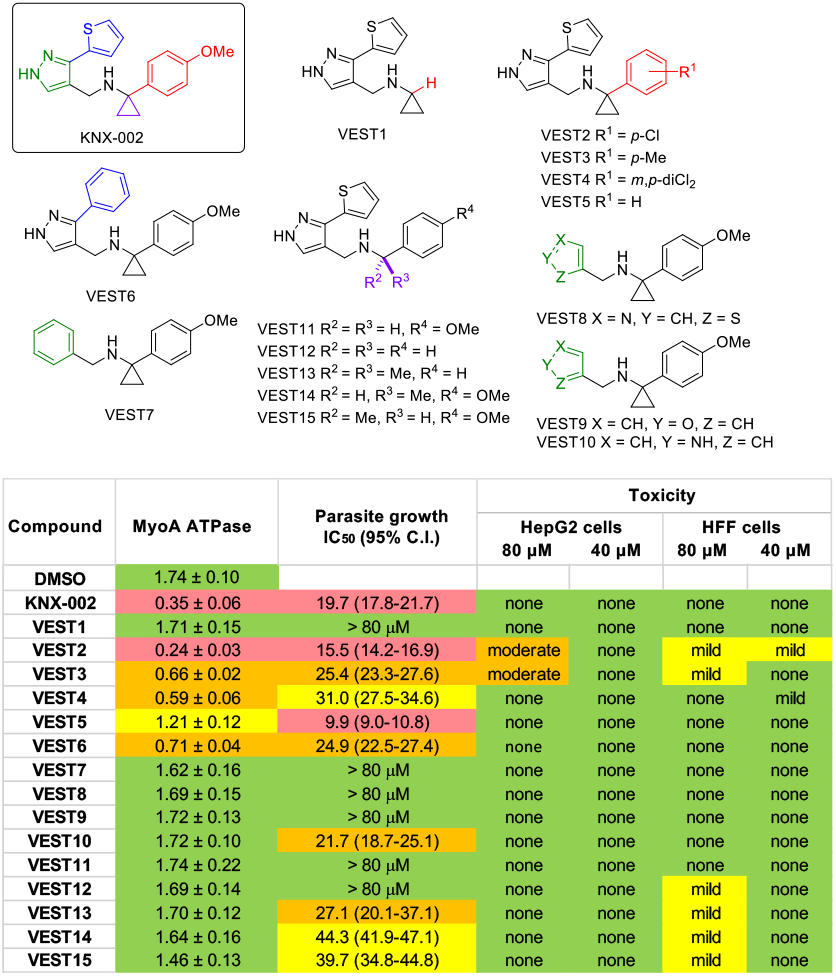
Preliminary structure-activity relationship (SAR) analysis. The structures of KNX-002 and 15 analogs (VEST1-15) are shown at the top. The table summarizes the activity of each of the compounds in assays measuring recombinant TgMyoA ATPase activity, parasite growth and mammalian cell toxicity; the data on which the table is based are shown in Suppl. Figures 3-5. <u>ATPase assay</u> data correspond to relative luminescence units × 10^−6^ (mean from 3 independent replicates +/-SEM) in the presence of 20 μM compound (Suppl. Figure 3). Green = less than 20% inhibition, yellow = 20-40% inhibition; orange = 45-70% inhibition; red = greater than 70% inhibition compared to vehicle (DMSO) controls. <u>Growth assay</u> data correspond to calculated IC_50_ and 95% confidence interval from a 5-6 day growth assay (Suppl. Figure 4). Red = IC_50_ less than 20 μM; orange = IC_50_ of 20-35 μM; yellow = IC_50_ of 35-50 μM; green = no detectable growth inhibition relative to vehicle (DMSO) controls. <u>Toxicity assay</u> entries summarize the results from 72-hour CellTox™ Green toxicity and CellTiter-Glo_®_ viability assays on both human foreskin fibroblasts and HepG2 cells. Thresholds for mild and moderate toxicity and loss of viability compared to vehicle (DMSO) controls are indicated by the colored bars on the right hand side of Suppl. Figure 5.

**Figure 4:**
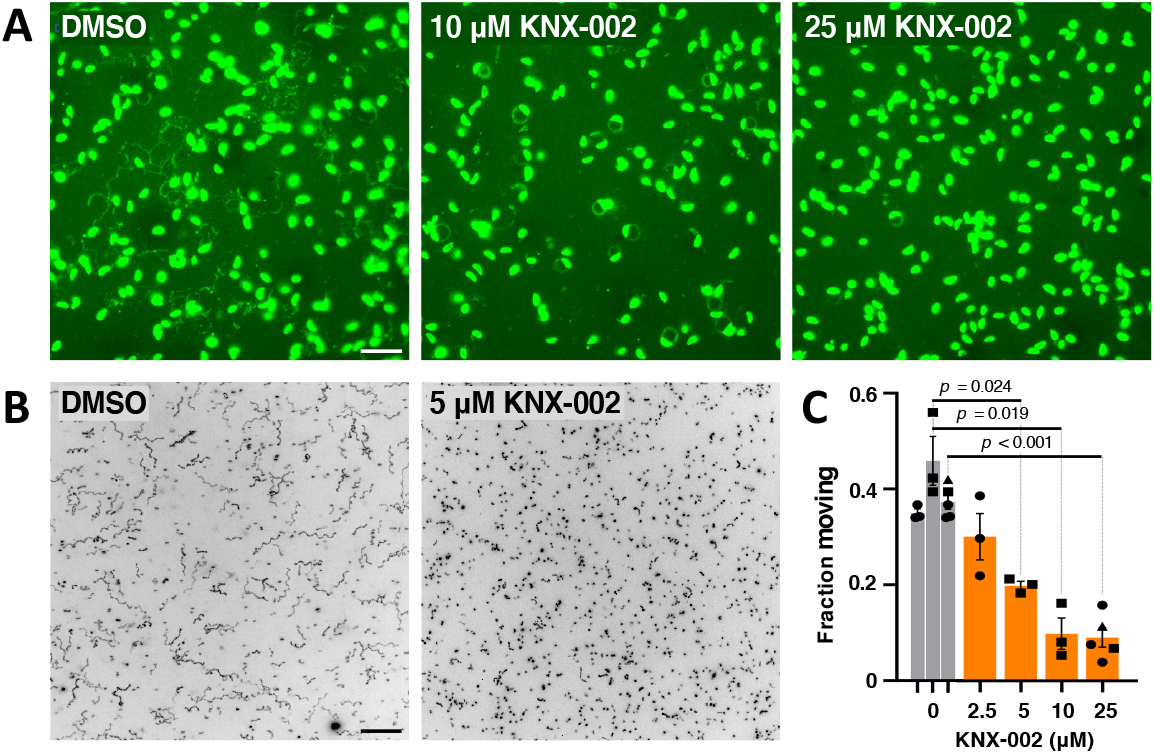
KNX-002 inhibits the motility of *T. gondii* tachyzoites. **(A)** Representative images from a 2D trail assay of parasites treated with DMSO (vehicle) only, 10 μM KNX-002, or 25 μM KNX-002. The parasites and trails were visualized by indirect immunofluorescence using an antibody against TgSAG1 after 15 min of gliding on the coverslip. Scale bar = 20 μm. **(B)** Representative maximum intensity projections showing parasite trajectories during 60 sec of motility in Matrigel in the absence (DMSO) or presence of 5 μM KNX-002. Scale bar = 40 μm. The grayscale images were inverted to provide clearer visualization of the trajectories. **(C)** Fraction of the parasite population moving during a 60 sec assay in the presence of the indicated concentrations of KNX-002 (IC_50_ = 5.3 μM, 95% C.I. = 3.1 – 7.8 μM). Each data point represents a single biological replicate consisting of three technical replicates. Sets of data captured on the same days, indicated by the similar symbol shapes, were compared by Student’s one-tailed paired t-tests.

**Figure 5:**
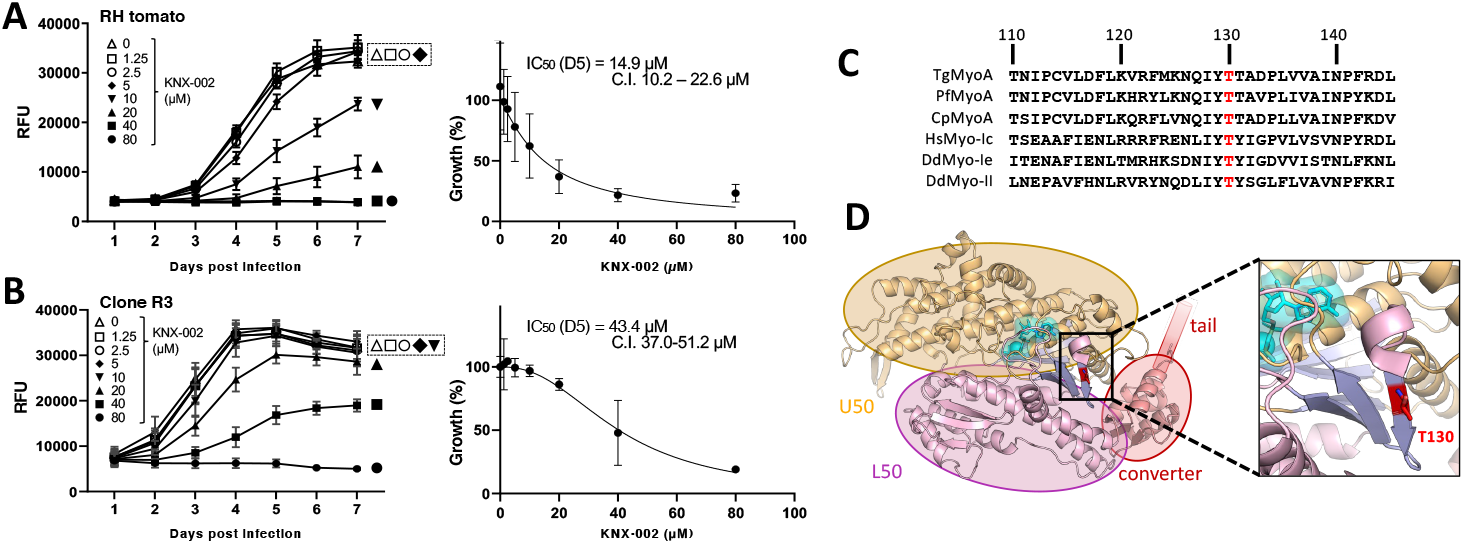
Identification of a mutant parasite showing reduced sensitivity to KNX-002 in growth assays. **(A, B)** Left panels: tdTomato-expressing wild-type parasites (A) or clone 3R (B), which was generated by mutagenesis and selection in KNX-002, were preincubated with various concentrations of KNX-002 for 5 minutes and then added to HFF cells on a 384-well plate. Fluorescence was measured daily over the next 7 days to quantify parasite growth in the continuing presence of compound. RFU = relative fluorescence units. The data shown are from three independent biological replicates, each consisting of 2-3 technical replicates at all time points. Right panels: the data from day 5 (D5) of the growth assays were used to calculate the IC_50_ and 95% confidence intervals (C.I.) of KNX-002 for parasite growth. Vertical bars indicate SEM (left panels) or 95% C.I. (right panels). **(C)** Multiple sequence alignment showing the high degree of conservation of T130 in apicomplexan MyoAs (Tg = *T. gondii*, Pf = *Plasmodium falciparum*, Cp = *Cryptosporidium parvum*) and MyoIc, Ie and II from *Homo sapiens* (Hs) and *Dictyostelium discoideum* (Dd). **(D)** Structure of the TgMyoA (PDB ID [6DUE]) motor domain, showing the position of T130 (red) relative to the converter, U50 and L50 subdomains.

**Figure 6:**
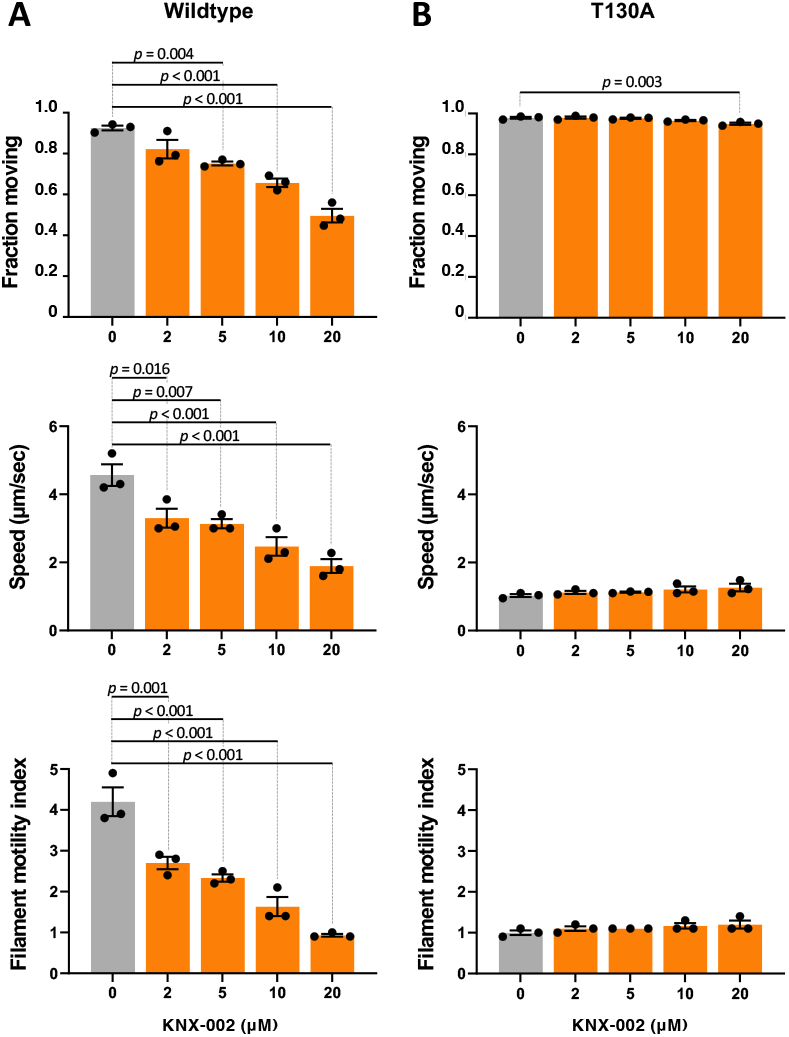
KNX-002 inhibits *in vitro* motility driven by wildtype but not T130A TgMyoA. *In vitro* motility assays, showing the fraction of actin filaments moving, their sliding speed, and the corresponding filament motility Index (fraction moving x mean speed) of TgMyoA treated with various concentrations of KNX-002, as indicated (0 = DMSO vehicle only). **(A)** Wildtype TgMyoA, **(B)** T130A TgMyoA. Bars show the mean of three independent experiments ± SEM. For each motility parameter, compound treatments were compared to DMSO by one-way ANOVA with Dunnett’s test for multiple comparisons. Only the statistically significant differences (*p* < 0.05) are indicated.

A series of additional analogs containing the *p*-OMe aryl substituent were also assessed. Substitution of the thiophene ring in KNX-002 for a phenyl ring (in **VEST6**, highlighted in blue, Figure 3A) led to a drop in activity in both the *in vitro* ATPase and growth inhibition assays. However, the retention of some biological activity by **VEST6** may provide a way to assess rapidly the importance of additional substitution in this region of the molecule in future studies through the incorporation of substituents on the new phenyl ring. No tolerance for replacement of the pyrazole (highlighted in green) and thiophene (blue) rings in KNX-002 was observed, with analogs containing only a phenyl, thiazole or furan substituent, instead of the thiophene-linked pyrazole, all being inactive in both assays (**VEST7, VEST8** and **VEST9**). Substitution of a pyrrole into KNX-002 to give **VEST10** led to loss of activity in the *in vitro* assay but retention of some activity in the growth inhibition assay. Finally, the importance of the cyclopropyl group in KNX-002 (highlighted in purple, Figure 3A) was confirmed by the lack of biological activity in either assay of the *des*-cyclopropyl analog **VEST11**. The growth inhibition activity associated with **VEST5** was also shown to require the cyclopropyl group as analogs **VEST12** (CH2,) and **VEST13** (CMe2) were both less active in the growth assay. Attempts to incorporate a stereogenic center through the replacement of the cyclopropyl group with a single methyl substituent led to analogs with decreased biological activity and mild cytotoxicity against HFF cells (**VEST14** and **VEST15**).

These preliminary studies have identified regions of KNX-002 important for its biological activity, which will inform subsequent efforts at compound optimization (see Discussion). Since no analogs with higher potency than KNX-002 were identified in this first round of SAR analysis, all experiments reported below were carried out using KNX-002.

### KNX-002 inhibits parasite motility

To test whether KNX-002 inhibits parasite motility, we first used a standard immunofluorescence-based assay that visualizes the protein “trails” left behind the parasites as they glide on a glass coverslip (*e*.*g*., [38, 39]). Untreated parasites deposit a variety of helical, meandering, and large diameter circular trails in this qualitative two-dimensional assay (Figure 4A). Treatment with 10 μM KNX-002 results in a markedly different motility pattern, consisting almost entirely of small diameter circles, and 25 μM KNX-002 dramatically inhibits the number of parasites undergoing any movement at all (Figure 4A).

To more quantitatively evaluate differences in parasite motility in the presence of KNX-002, we used a 3D motility assay that automatically analyzes thousands of trajectories in a single experiment, providing robust statistical comparisons between populations of parasites [16, 21]. KNX-002 treatment decreases the fraction of the parasite population moving in the 3D assay in a dose-dependent manner (Figures 4B and 4C; IC50 = 5.3 μM, 95% C.I. 3.1-7.8 μM). Of those parasites that move, KNX-002 treatment also reduces their average displacement and speed (Suppl. Figure 7).

**Figure 7:**
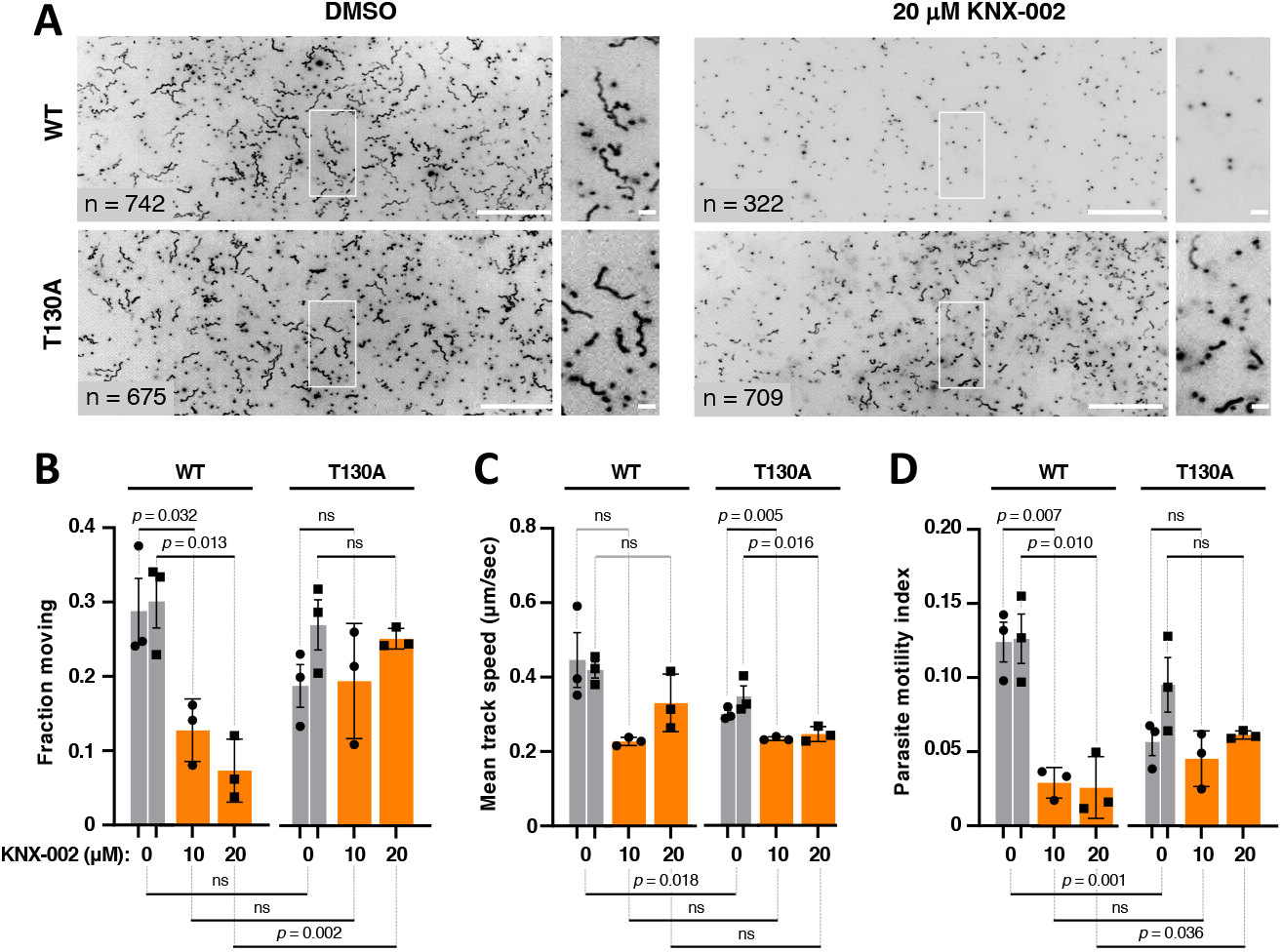
The TgMyoA T130A mutation reduces parasite sensitivity to KNX-002 in 3D motility assays. **(A)** Large panels: representative maximum intensity projections showing the trajectories of wildtype (WT) and T130A mutant parasites during 80 sec of motility in Matrigel, in the presence of DMSO (vehicle control) or 20 μM KNX-002. The grayscale images were inverted to provide clearer visualization of the trajectories; scale bar = 100 μm. The boxed area in each large panel is magnified and shown on the right; scale bar = 10 μm. **(B)** Percent of wild-type (WT) and T130A mutant parasites moving in the presence of DMSO (vehicle), 10 μM KNX-002, or 20 μM KNX-002. **(C)** The mean speeds of the parasites analyzed in panel B. **(D)** The corresponding parasite motility index (fraction of parasites moving x mean speed). Each data point in panels B-D represents a single biological replicate composed of three technical replicates. Three of the biological replicates were collected on the same three days (circles), and three of the biological replicates were collected on a different three days (squares). Bars show the mean of the biological replicates ± SEM. Sets of biological replicates using the same parasite line and collected on the same days were compared by Student’s one-tailed paired t-tests (significance indicated above the graphs). Sets of biological replicates comparing different parasite lines were analyzed by Student’s two-tailed unpaired t-tests (significance indicated below the graphs). ns = not significant *(p* > 0.05).

### Identification of a mutation in TgMyoA that confers partial resistance to KNX-002

The results presented thus far show that KNX-002 is an inhibitor of TgMyoA activity, parasite growth in culture, and parasite motility. The data do not, however, directly link the compound’s effects on parasite motility and growth to inhibition of TgMyoA function; it remains possible that the inhibition of parasite motility and growth are due to off-target effect(s). As a first step towards validating the specificity of the compound, we chemically mutagenized parasites with N-ethyl-N-nitrosourea [40] and selected for parasites resistant to KNX-002 by growth in 40 μM KNX-002. Parasites that grew up after 10-14 days under selection were cloned and their sensitivity to KNX-002 determined using the plate-based growth assay. We isolated 26 such clones from two independent mutagenesis/selection experiments, five of which showed at least a 2.5-fold increase in IC50 for KNX-002 (Suppl. Figure 8). cDNA was generated from the five clones, and each of the motor constituents (TgMyoA, TgMLC1, TgELC1 and TgELC2) was sequenced. Four of the five resistant clones contained no amino acid-altering mutations in the motor genes; the fifth (clone R3) contained a single A to G point mutation in the coding sequence of TgMyoA that changes threonine130 to alanine (T130A). This clone showed a 2.9-fold increase in IC50 for KNX-002 in the growth assay (Figure 5A,B), from 14.9 μM (wildtype; 95% C.I. = 10.2 to 22.6 μM) to 43.4 μM (T130A mutant; 95% C.I. = 37.0 to 51.2 μM). T130 is highly conserved in eukaryotic myosins (Figure 5C) and located on the first of seven β-strands comprising the transducer subdomain (Figure 5D). The residues immediately flanking T130 are also well conserved between the MyoA homologs of other apicomplexan parasites, but diverge in the vertebrate myosins examined (Figure 5C).

**Figure 8:**
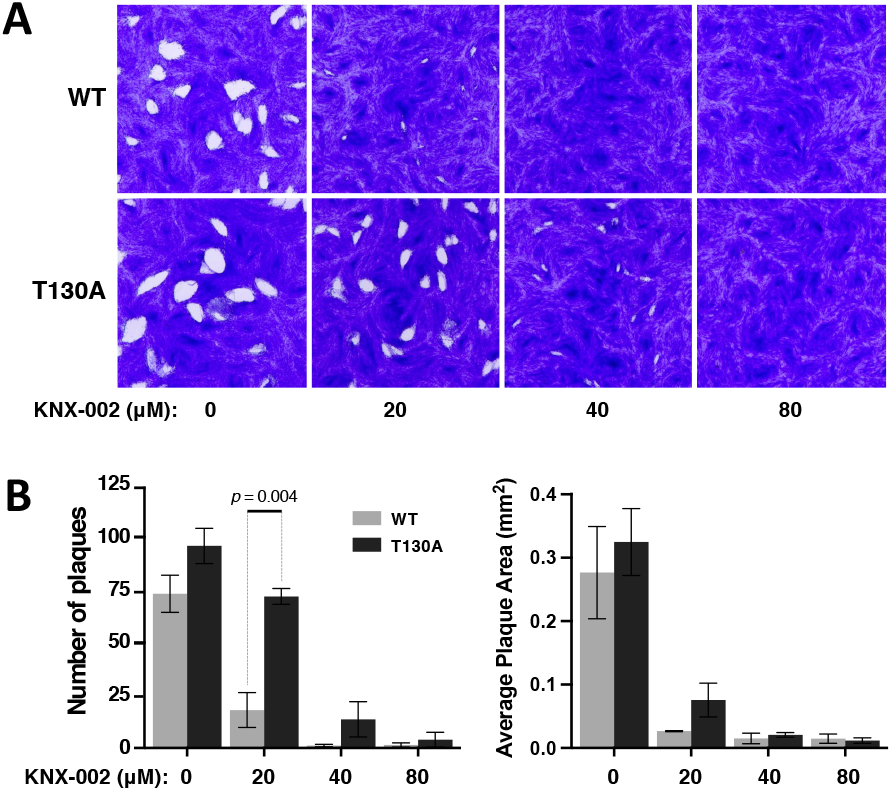
The TgMyoA T130A mutation reduces parasite sensitivity to KNX-002 in growth assays. **(A)** Representative images of plaque assays using wild-type (WT; top row) and T130A mutant (bottom row) parasites in the presence of 0 (DMSO vehicle only), 20, 40, and 80 μM KNX-002, 7 days after inoculating confluent HFF monolayers with 200 parasites/well. An 11.3 × 11.3 mm area from the middle of the well is shown; scale bar = 1 mm. **(B)** Total number of plaques per well and average plaque area of HFF monolayers inoculated with 200 parasites/well (WT or T130A) and grown for 7 days in the presence of 0 (DMSO), 20, 40, or 80 μM KNX-002. Bars show the mean of three biological replicates ± SEM. Treatments were compared by Student’s two-tailed unpaired t-test; only the statistically significant differences (*p* < 0.05) are indicated.

To independently establish that the T130A mutation alters the sensitivity of TgMyoA to KNX-002, we expressed recombinant TgMyoA containing the T130A mutation and compared its motion-generating capacity to that of wild-type TgMyoA in an *in vitro* motility assay that measures the speed with which fluorescently labeled actin filaments are moved by myosin adsorbed to a glass coverslip [20, 41]. In the absence of added compound, the wild-type motor moved 92.5± 1.2% of the filaments on the coverslip, with an average mean speed of 4.6 ± 0.3 μm/sec (Figure 6A, top and middle panels, grey bars). The T130A mutation had no effect on the fraction of filaments moving in the assay, but decreased the basal speed of filament sliding by 77 ± 3% compared to wild-type (Figure 6B, top and middle panels, grey bars). Accordingly, we tested whether the mutant was misfolded, prone to aggregation, or otherwise unstable by comparing its size-exclusion chromatography (SEC) elution profile and thermal stability, as measured by differential scanning fluorimetry (DSF), to that of the wild-type motor (Suppl. Figure 9; both proteins assayed in the near-rigor [SEC] and pre-powerstroke [DSF] states). No mutation-associated differences were seen in either assay, suggesting that the effects of the T130A mutation on motor function are not due to large structural changes in the protein caused by the amino acid substitution.

**Figure 9:**
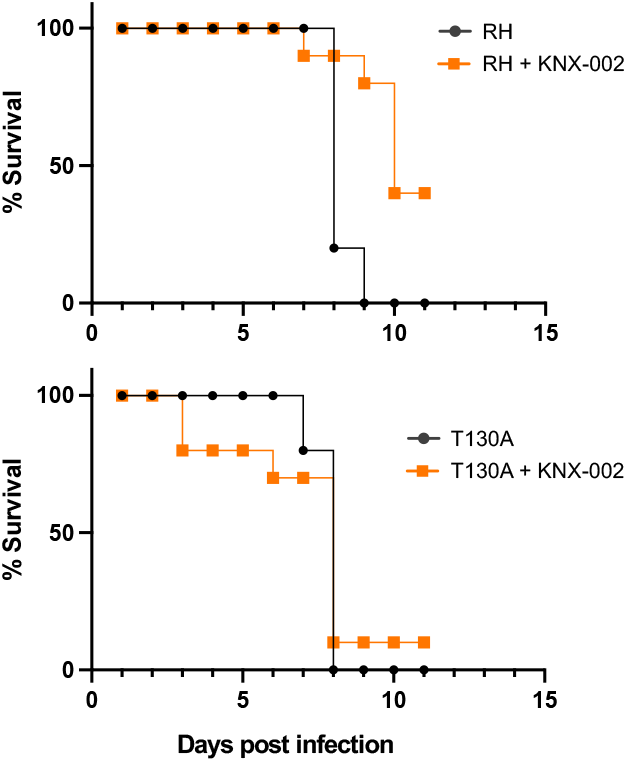
KNX-002 slows disease progression in mice infected with wild-type but not T130A mutant parasites. Four groups of mice were injected intraperitoneally on day 0 with either KNX-002 (20mg/kg) or the equivalent volume of DMSO (vehicle). Thirty minutes later, the mice were infected with 500 wild-type parasites (RH; top panel) or 500 parasites expressing T130A mutant myosin (bottom panel). On day 2, the mice were treated again either with compound or DMSO. Differences between the survival curves were analyzed using a log-rank Mantel-Cox test (*p* = 0.004 for RH *vs* RH + KNX-002 [top panel]; *p* = 0.922 for T130A *vs* T130A + KNX-002 [bottom panel]); and *p*=0.008 for RH + KNX-002 *vs* T130A + KNX-002 (orange curves).

Including KNX-002 in the *in vitro* motility assay with wild-type motor inhibits both the fraction of actin filaments moving and the speed of those filaments that do move, each in a dose-dependent manner (Figure 6A, top and middle panels). Combining these two effects into a single “filament motility index” (*i*.*e*., the fraction of filaments moving multiplied by the mean speed of the filaments that do move) provides a better measure of the overall effect of the compound in the assay. KNX-002 induces a large decrease in the filament motility index (Figure 6A, bottom panel).

In marked contrast to the wild-type motor, KNX-002 had little to no effect on the fraction of actin filaments moved by the motor containing the T130A mutation (Figure 6B, top panel). Furthermore, although the basal filament sliding speed of the mutant motor was lower than that of wildtype, this reduced sliding speed was also insensitive to compound (Figure 6B, middle panel), as was the mutant motor’s overall filament motility index (Figure 6B, bottom panel).

Taken together, these data confirm that the T130A mutation renders TgMyoA less sensitive to the inhibitory effects of KNX-002.

### Validation of the specificity of KNX-002 in parasites

The resistant parasite line generated by mutagenesis and selection (clone R3, Figure 5B) is expected to contain as many as 70 different mutations throughout its genome [40] in addition to the T130A mutation. To confirm that resistance to KNX-002 in this parasite line was due to the identified mutation in TgMyoA, we recreated the single T130A mutation in the wild-type background using CRISPR/Cas9 (Suppl. Figure 10). To quantitatively evaluate effects of the mutation on the parasite’s lytic cycle, we compared wild-type and T130A parasites in a plaque assay, which encompasses the parasite’s entire lytic cycle. The T130A mutation had no significant effect on the number of plaques formed or plaque area (Suppl. Figure 11A, B), demonstrating little to no effect of the mutation on the parasite’s lytic cycle or growth in culture. Consistent with the plaque assay results, the T130A mutation also had little to no effect on the fraction of the parasite population moving in the 3D motility assay (compare the leftmost pair of panels in Figure 7A and two sets of grey bars in Figure 7B) and only a minor effect on their mean trajectory speeds (compare the two sets of grey bars, Figure 7C). Finally, the T130A mutation had no detectable effect on the virulence of the parasite in a mouse model of infection (Suppl. Figure 11C).

We next asked whether parasites expressing the T130A mutation were less sensitive to KNX-002 in the 3D motility assay). As shown above (Figure 4C), KNX-002 strongly inhibits the fraction of wild-type parasites that move in Matrigel (Figure 7B, left three sets of bars). In contrast, the fraction of T130A parasites that move during the assay was insensitive to KNX-002, up to concentrations of 20 μM (Figure 7B, right three sets of bars). The parasite speed data are more variable (Figure 7C and Suppl. Figure 7B) but show that the mutant parasites retain some sensitivity to the compound (Figure 7C, right three sets of bars). To capture the combined impact of changes to these two motility parameters and to facilitate comparison to the *in vitro* motility results with recombinant motor, we calculated the “parasite motility index” (the fraction of parasites moving multiplied by the average speed of those that do move) under each condition. The wild-type parasite motility index is more severely impacted by KNX-002 treatment than the motility index of T130A parasites (Figure 7D).

The displacement of both wild-type and mutant parasites is also inhibited by KNX-002 (Suppl. Figure 7A and Suppl. Figure 12), but the mutants are again less sensitive to the compound, *i*.*e*., despite moving shorter distances in the absence of compound (compare the two sets of grey bars in Suppl. Figure 12), the T130A parasites moved farther than wildtype in the presence of 20 μM KNX-002 (compare the third and sixth bars in Suppl. Figure 12). The decreased sensitivity of the T130A parasites to the effects of KNX-002 is apparent in the maximum intensity projections from the 3D motility assay, which show more T130A parasites than wildtype moving in the presence of 20 μM KNX-002, with longer average displacements (compare right two sets of panels in Figure 7A).

Plaque assays were used to compare the growth of wildtype and mutant parasites in culture in the presence of KNX-002. Parasites expressing the mutant motor formed a significantly larger number of plaques in the presence of 20 μM compound than wild-type parasites, and even at 40 μM KNX-002 the mutant parasites were still able to generate small plaques (Figure 8).

These data demonstrate that the T130A mutation confers partial resistance to the compound’s effects on motor function, parasite motility and parasite growth, confirming that TgMyoA is a biologically relevant target for KNX-002 in *T. gondii*.

### KNX-002 treatment slows disease progression in mice infected with a lethal dose of *T. gondii*, through a specific effect on TgMyoA

Finally, we tested whether KNX-002 (20mg/kg, administered intraperitoneally on the day of infection and two days later) can alter the progression of disease in a mouse model of infection. Only 20% of the mice that were infected with wild-type parasites and treated with vehicle were still alive 8 days post-infection, and none survived to day 9 (Figure 9, top panel, black circles). In marked contrast, 90% of the mice treated with KNX-002 survived to day 8, 80% to day 9, and 40% were still alive on day 11 when the experiment was terminated (Figure 9, top panel, orange squares).

When the same experiment was done with parasites expressing the T130A mutant myosin, the vehicle-treated mice did not survive beyond day 8 (Figure 9, bottom panel, black circles), similar to what we observed with wild-type parasites. Critically, and in marked contrast to infection with wild-type parasites, KNX-002 treatment did not significantly improve the survival of mice infected with the T130A parasites (Figure 9, bottom panel, orange squares).

Together, these data show that KNX-002 can slow disease progression in mice infected with *T. gondii*, and prove that the compound exerts its effects through inhibition of its intended target, TgMyoA.

## Discussion

The intense interest in motility among those who study apicomplexan parasites reflects both the unique nature of the process and its importance in parasite biology and virulence. These parasites move without the cilia, flagella or protruding leading edge that drives the motility of most other eukaryotic cells. The class XIV myosins clearly play a central role in motility, but they are unusual myosins in many respects [17, 23], and precisely how these myosins and their interacting proteins function to drive motility remains controversial [27, 42-47]. The mechanisms underlying motility are therefore of fundamental cell biological interest. Since MyoA is essential for virulence, highly conserved in apicomplexan parasites, and different in several respects from the myosins found in humans, the MyoA motor complex also represents a potentially attractive target for drug development. We describe here the identification and characterization of a novel small molecule, KNX-002, which will be a useful chemical probe for future studies of the function of TgMyoA. Importantly, we also identified a mutation in TgMyoA that reduces the sensitivity of the motor to KNX-002. We used isogenic parasites lines expressing either the wild-type or mutant myosin to unequivocally demonstrate that TgMyoA is a biologically relevant target of the compound in parasites and to show that pharmacological inhibition of TgMyoA can slow the progression of disease in mice infected with a lethal dose of *T. gondii*.

### KNX-002 as a chemical probe for studying class XIV motor function

Most previous studies of TgMyoA motor function involved phenotypic characterization of parasites in which the genes encoding TgMyoA or its associated proteins were either disrupted or their expression downregulated (*e*.*g*., [26, 27, 46, 48]). Such reverse genetic approaches, particularly gene disruptions, are powerful but blunt tools for studying the mechanisms underlying processes important to parasite survival, such as motility. Pharmacological inhibitors that interact specifically with a target of interest provide a useful complementary approach for studying motility and other facets of parasite biology [49, 50], since different aspects of the target’s function can be perturbed at specific times and in a controlled manner. Because the parasites are exposed to the compound for only short periods of time, compensatory adaptations (*e*.*g*., through mutation or changes in gene expression) that can confound data interpretation in experiments involving gene disruption [31, 32] are unlikely to occur. Even experiments involving controlled protein depletion or transcriptional downregulation of a gene of interest, which can be achieved in hours to days, run the risk of inducing compensatory changes in transcription, translation, protein stability, post-translational modifications, or the localization of proteins with overlapping function (*e*.*g*., [30-32]). Determining the effect of a pharmacological inhibitor of motility can, in contrast, be accomplished within minutes (*e*.*g*., Figures 4, 7).

There have been only two previously reported attempts to pharmacologically manipulate the function of TgMyoA. In the first, the general myosin inhibitor 2,3-butanedione monoxime (BDM) was shown to inhibit parasite motility [38]; however, BDM is now recognized as an inhibitor with so many off-target effects that little can be concluded from its use in cells [51, 52]. The other study used computational methods and a homology model of TgMLC1 bound to TgMyoA to identify two predicted inhibitors of TgMLC1-TgMyoA interaction [53]. These two compounds (C3-20 and C3-21) inhibit *T. gondii* growth with sub-micromolar IC50s [53] but whether this effect on parasite growth is due to inhibition of TgMyoA has not been established.

The compound we have identified and characterized here, KNX-002, inhibits the ability of recombinant TgMyoA to propel actin filaments in the *in vitro* motility assay, and decreases the speed of the filaments that move (Figure 6A). Similar effects are seen in the 3D motility assay: compound treatment reduces both the number of parasites moving and, to a lesser extent, the speeds with which these parasites move (Figure 7B, C and Suppl. Figure 7B). To provide a better picture of the combined impact of changes to these two motility parameters, we multiplied the fraction of actin filaments or parasites moving by the mean actin filament/parasite speed to generate the filament/parasite motility index. In the *in vitro* motility assay with wild-type TgMyoA, fraction moving and speed contributed approximately equally to the filament motility index in both the presence and absence of the compound. In wild-type parasites, the compound-induced changes to fraction moving are larger than to speed and therefore the predominant contributor to the changes in parasite motility index upon compound treatment. Precisely how the effects of the compound on motor function in the *in vitro* motility assay relate to the observed changes in parasite motility will require further study. Nevertheless, the data presented demonstrate the value of KNX-002 as a new chemical probe for studying the mechanism(s) underlying the unusual form of eukaryotic cell motility used by *T. gondii*.

Excitingly, KNX-002 and a recently discovered analog named KNX-115 were shown to inhibit PfMyoA, the TgMyoA homolog of the malaria parasite, *P. falciparum* [54, 55], and to block the growth of blood stage *P. falciparum* in culture and the invasion of hepatocytes by liver stage parasites [54, 55]. Structure-function studies revealed that the compound sequesters PfMyoA in a state of low affinity for actin [54], consistent with a decrease in duty ratio we observe in wild-type TgMyoA treated with KNX-002 (Suppl. Figure 13).

### Effects of the T130A mutation

The T130A mutation that confers partial resistance to KNX-002 was identified by chemical mutagenesis and selection in KNX-002. While the mutagenesis/selection approach is commonly used to generate resistant parasites in *T. gondii* and other apicomplexan parasites (*e*.*g*., [56-58]), the next step after isolating resistant clones is typically to do whole genome sequencing on the most promising of the resistant clones to identify potentially relevant mutations. We took a more direct and scalable approach in this study, by sequencing only the motor protein loci (TgMyoA, TgMLC1, TgELC1, TgELC2) in the resistant parasites, since (a) the goal of this study was to identify a direct inhibitor of motor function, and (b) we had already shown that KNX-002 can inhibit the activity of recombinant motor consisting only of TgMyoA, TgMLC1, and TgELC1 (note: since TgELC1 and TgELC2 are functionally redundant [19], a mutation in the gene encoding either could confer compound resistance). Of the five resistant clones whose IC50 for KNX-002 shifted 2.5-fold or more relative to wildtype in the growth assay, only one had a mutation in the coding sequence of one of the motor protein genes, T130A in TgMyoA. The source of resistance in the other four clones is unknown.

T130, which lies in the first *β*-strand at the extreme N-terminus of the transducer domain [24], is not predicted to be directly involved in the binding of KNX-002 [54], even though other aspects of the transducer may be involved [54]. Thus, the mechanism by which the T130A mutation confers partial resistance to the compound is likely to be indirect. For example, the T130A mutation could induce long range structural changes that alter the KNX-002 binding pocket, thereby mitigating the effect of the compound. However, myosin motors depend on a complex and highly interconnected network of allosteric interactions to perform their function [59], making it difficult to predict precisely how the T130A mutation reduces the sensitivity of TgMyoA to KNX-002.

While the T130A mutation reduces the sensitivity of the motor to KNX-002, the mutation is not without its effects on motor performance, causing a 75-80% reduction in basal filament sliding speed in the *in vitro* motility assay (Figure 6B). However, when expressed in parasites the mutation had a much less pronounced effect on parasite motility (compare the first and fourth sets of grey bars in Figures 7B-D). We explored several possible explanations for this differential effect of the T130A mutation on the activity of the recombinant motor *vs*. motor-driven motility of the parasites. First, the mutation might alter the folding or stability – and therefore the activity – of the recombinant TgMyoA used in the *in vitro* assays. This seems unlikely, since the mutant motor was indistinguishable from the wild-type motor by SEC and DSF (Suppl. Figure 9). Second, the partially functional motor might be overexpressed in the mutant parasites. This possibility was ruled out by quantitative western blotting (Suppl. Figure 14A). Third, the parasite might compensate for the loss of TgMyoA activity in the T130A mutants through changes in the transcription of other myosins, myosin light chains, glideosome components, actin or actin regulatory proteins. No such changes to known motility-associated proteins were observed by RNAseq analysis (Suppl. Figure 14B, right panel). Furthermore, analysis of all differentially regulated transcripts revealed that 57% of the genes up/down regulated more than log2-fold were annotated as hypothetical proteins, and the remainder were largely associated with trafficking and metabolism (Suppl. Figure 14B, left panel) so cannot readily explain a compensation mechanism in parasites with impaired motor function, although further studies would be required to definitively rule out this possibility. The compound might have less of an effect on motor function in the parasite due to some protein or factor present in the parasite (but not the insect cell system used to produce recombinant motor) that can counteract the deleterious effects of the mutation. Alternatively, the density of TgMyoA motors might be higher in the parasite pellicle than on the glass coverslips used in the *in vitro* motility assays, and therefore require higher concentrations of compound to achieve equivalent levels of inhibition. Regardless of the reason for this differential impact of the mutation on *in vitro* and parasite motility, it is notable that the fraction of actin filaments moving *in vitro* and the fraction of parasites moving within Matrigel were better correlated than the speeds of actin filament and parasite movement. The fraction of filaments moving in the *in vitro* motility assay may therefore be more relevant to motor-driven parasite motility than filament sliding speed.

The mutagenesis protocol used to generate the T130A mutation kills 70% of the initial parasite population and results in an average of 63 mutations per parasite across the genome [40]. It is unclear how these harsh experimental mutagenesis conditions relate to what occurs during a natural infection and therefore how readily parasites showing reduced sensitivity to KNX-002 (or, more likely, an optimized compound based on the KNX-002 scaffold) would arise during treatment of a human infection. From the drug development perspective, it would be useful to know what other resistance mechanisms are available to the parasite for this class of compounds. Further characterization of the four partially resistant mutants identified here that have no mutations in the TgMyoA motor (Suppl. Figure 8) would be a good place to start. TgMyoA knockout parasites are still capable of a small degree of 3D motility (13% of wild-type levels [46] under the same assay conditions used here). Although controversial [46], it has been proposed that TgMyoC can to some extent functionally compensate for the loss of TgMyoA [30, 31], so mutations in TgMyoC would be of particular interest. It should be noted that because humans are a dead-end host for *Toxoplasma*, even if resistance were to emerge within a treated individual through mutations in TgMyoA or some other locus in the genome, the chances of this resistance spreading human-to-human or into the zoonotic reservoir are exceedingly small.

### MyoA as a druggable target

A few small molecule inhibitors of mammalian myosins have shown potential as therapeutics (reviewed in [51]). The most well characterized myosin inhibitor is blebbistatin, a noncompetitive inhibitor of myosin II [60]. Blebbistatin has been a useful chemical probe for studying myosin II function [51], and derivatives are being considered for cancer chemotherapy [61, 62] and drug addiction [63]. A more recently discovered inhibitor, MYK-461 (mavacamten), decreases the duty ratio of cardiac muscle myosin through an effect on Pi release, reduces the development of ventricular hypertrophy in a mouse model of hypertrophic cardiomyopathy (HCM; [64]), and has been approved by the FDA for use in the treatment of hypertrophic cardiomyopathy patients.

The MyoA motor complex of apicomplexan parasites has long been considered an attractive target for drug development [25, 33, 53, 65-67]. However, no previous studies have tested whether small molecule inhibitors of TgMyoA or other apicomplexan myosins can alter the course of infection *in vivo*. The argument against MyoA as a drug target has been that the parasite may have alternative and/or compensatory mechanisms to enable motility in the absence of MyoA [26, 27, 43, 45, 46], meaning that simultaneous inhibition of more than one motility pathway may be necessary to provide sufficient protection against infection. However, *T. gondii* engineered to express low levels of TgMyoA, which may be more analogous to drug-treated parasites than a knockout, are completely avirulent [28], arguing that sufficiently strong inhibition of TgMyoA is likely, on its own, to be therapeutically useful.

Given this rationale, a major motivation behind the work reported here was to develop the tools necessary to rigorously test whether MyoA can be targeted by small molecules to alter the course of an infection. We demonstrated that it is possible to identify small molecules that inhibit parasite TgMyoA, have no detectable effect on vertebrate myosins, and show no toxicity to mammalian cells. Furthermore, one of these compounds, KNX-002, was shown to be capable of inhibiting myosin function in the parasite, thereby disrupting parasite motility and the lytic cycle. The specificity of KNX-002 for TgMyoA in these experiments was demonstrated using parasites expressing the T130A mutation, and the observation that treatment with KNX-002 decreases the susceptibility of mice to a lethal infection with wild-type, but not T130A parasites, proves that TgMyoA is indeed a druggable target, *in vivo*. Whether KNX-002 can also inhibit the activity of any of the parasite’s other ten myosins [68] and disrupt their biological function(s) will be the subject of future study.

While KNX-002 provided the means to rigorously test the druggability of TgMyoA, it caused weight loss and histological evidence of liver damage in the treated infected mice. Before further animal work, it will therefore be necessary to develop more potent and less toxic analogs that retain specificity for parasite myosin. Our preliminary SAR data (summarized in Figure 3) are consistent with a key role for the pyrazole, cyclopropyl, and *p*-OMe aryl units within KNX-002, with a generally good correlation between the results of the recombinant myosin ATPase and parasite growth inhibition assays.

Subsequent studies, inspired both by our SAR data and the recently reported structure of the PfMyoA-KNX-002 complex [54], could explore the effect on biological activity of: (i) alternative replacements for the thiophene group and (ii) a switch in the positions of the NH and CH2 groups, amongst other options. Encouragingly, Trivedi *et al* recently identified an analog of KNX-002 that shows >20-fold improved potency against recombinant PfMyoA [55]. Although neither the structure of this analog nor the SAR strategy that led to its discovery were disclosed, this result suggests there is considerable opportunity to optimize the biological properties of the KNX-002 scaffold through a systematic SAR campaign.

KNX-002 and any improved analogs developed through further rounds of iterative SAR will be useful new experimental tools for studying the role of MyoA in parasite growth, dissemination and virulence, and provide an exciting opportunity to develop an entirely new class of drugs that can be used to prevent or treat the severe diseases caused by *T. gondii* and other apicomplexan parasites.

## Methods

### Cell and parasite culture

Parasites were propagated by serial passage in human foreskin fibroblasts (HFFs). HFFs were grown to confluence in Dulbecco’s Modified Eagle’s Medium (DMEM) (Life Technologies, Carlsbad, CA) containing 10% v/v heat-inactivated fetal bovine serum (FBS) (Life Technologies, Carlsbad, CA), 10 mM HEPES pH 7, and 100 units/ml penicillin and 100 μg/ml streptomycin, as previously described [69]. Prior to infection with *T. gondii*, the medium was changed to DMEM supplemented with 10mM HEPES pH 7, 100 units/ml penicillin and 100 μg/ml streptomycin, and 1% v/v FBS.

### Compounds

KNX-002 was either obtained commercially from Asinex (Winston-Salem NC; Figures 1, 2, 4-8 and S1, S2, S6-S8, S11-13) or synthesized, purified and characterized as described below (Figures 3, 9 and S3-5, S10). The commercially sourced and in-house synthesized compounds had similar IC50s in parasite growth assays: see IC50 values and overlapping 95% confidence intervals in Figures 2 and 5 (commercial) and Figure 3 (synthesized). Compounds were dissolved in dimethyl sulfoxide (DMSO) and stored at -20ºC, in the dark, until use. In all experiments involving treatment with compound, an equivalent amount of DMSO was added to all samples in that experiment, including the DMSO only vehicle controls. Final concentrations of DMSO were 0.5% v/v for experiments with recombinant TgMyoA, 0.1% v/v for experiments with parasites and cells, and 34 μl DMSO per injection per 20g mouse for the *in vivo* experiments.

### High-throughput screen

Compound CK2140597 was originally identified through high-throughput screening of a library of 50000 compounds (40uM final concentration, 2% v/v DMSO), performed by JS and EW at Cytokinetics, Inc., to identify inhibitors of the actin-activated ATPase of TgMyoA, provided by AK and GEW. The same compounds were screened, in parallel, against *Plasmodium falciparum* MyoA (PfMyoA; protein provided by Kathy Trybus, University of Vermont). The assay used was a coupled enzymatic assay [70]. Actin was first polymerized using 1 mM ATP, 2 mM MgCl2, 20 mM KCl and left on ice for 30 minutes. Buffer A was made with PM12 (12 mM Pipes-KOH, pH 7.0, 2 mM MgCl2), 1mM DTT, 0.1 mg/ml BSA, 0.4 mM PK/LDH, 1 μg/ml TgMyoA, 0.009% v/v Anitfoam (Millipore Sigma, St. Louis MO). Buffer B was made with PM12, 1 mM DTT, 0.1 mg/ml BSA, 1 mM ATP, 1 mM NADH, 1.5 mM PEP, 0.6 mg/ml previously polymerized actin and 0.009% v/v Antifoam. 12.5 μl of each buffer were added to a plate with a Packard MiniTrak liquid handler and then mixed at 2400 for 45 seconds with an BioShake 3000 ELM orbital plate mixer (Qinstruments). The decrease in optical density at 340 nm as a function of time was measured on an Envision 2104 spectrophotometer (Perkin Elmer) with a 600 s data monitoring window sampled every 60 s. Hit progression included confirmation of activity in an independent experiment, inactivity against an unrelated ATPase (hexokinase from Millipore Sigma), and dose-responsive inhibition. Hexokinase reactions were performed using glucose (1 mM) and sufficient hexokinase to generate ADP from ATP at a rate similar to the target myosins. Based on pre-established thresholds, CK2140597 was identified as a hit in the TgMyoA but not the PfMyoA screen; however, resupplied compound was subsequently shown to also be active against PfMyoA by Trybus and colleagues. CK2140597 was licensed by Kainomyx (www.kainomyx.com), renamed KNX-002, and released for academic studies in the laboratory of GEW (for *Toxoplasma*) and Kathy Trybus (for *Plasmodium*).

### Vertebrate myosin ATPase assays

Myofibril preparations from striated muscles are a well-established model for assaying myosin ATPase activity in a semi-native context [71]. Bovine cardiac, rabbit fast skeletal, and bovine masseter slow skeletal myofibrils were prepared and assayed as previously described [72], except they were tested at 75% calcium activation (pCa75) since this high level of basal ATPase activity enables the sensitive detection of inhibitors. For smooth muscle myosin (SMM), native chicken gizzard SMM S1 was prepared, crosslinked to bovine cardiac actin using EDC/NHS, and assayed as previously described [73].

### Expression and purification of recombinant TgMyoA

TgMyoA was co-expressed with TgMLC1, TgELC1, and TgUNC in Sf9 cells and purified as previously described [33]. TgMyoA containing the T130A mutation was generated using the QuikChange II-Site-Directed Mutagenesis method (Agilent Scientific, Santa Clara CA). Briefly, mutant strand synthesis was performed by denaturing baculovirus plasmid pSmTgMyoAcBiocfFLAG, annealing primers P21 and P22 (Table 1), which contain the desired mutation and using PfuUltra high fidelity taq polymerase in a thermocycling reaction to incorporate and extend the primer product. The product was then digested by DpnI overnight at 37 °C and the resulting plasmid was purified using a Qiagen PCR purification kit. Subsequently, the purified PCR product was used to transform electro-competent DH5-alpha cells. Positive clones were screened, plasmid prepped and the presence of the mutation verified by sequencing.

**Table 1:**
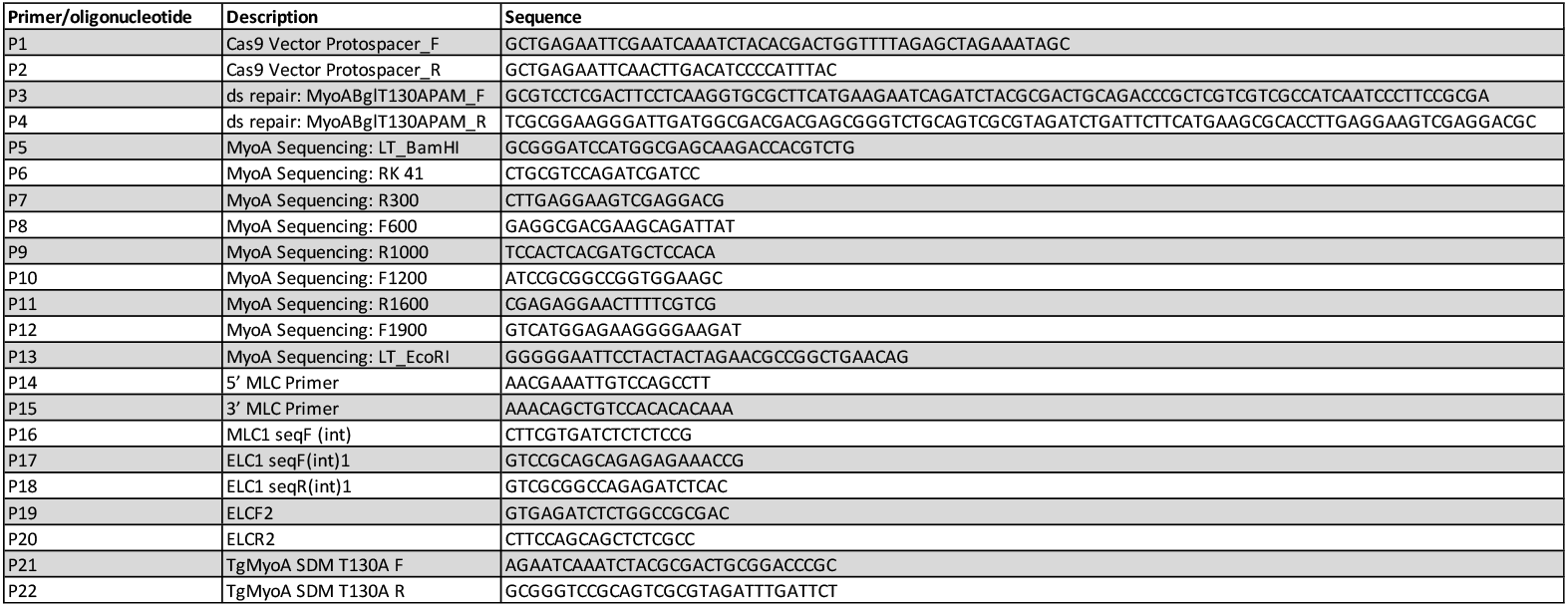
Primers and oligonucleotides used in this study.

### *In vitro* motility

*in vitro* motility assays with chicken skeletal muscle actin were performed as previously described [33] with the following modifications: (a) Neutravidin (40 ug/ml, Molecular Probes) in buffer B was added to the flow cells first, followed by BSA (in buffer B) and 3 washes with buffer B; (b) approximately 0.16 μg of FLAG purified TgMyoA was used per flow cell; (c) Buffer C was in all cases supplemented with oxygen scavengers (3 mg/ml glucose (Sigma-Aldrich), 0.125 mg/ml glucose oxidase (Sigma-Aldrich), and 0.05 mg/ml catalase (Sigma-Aldrich); (d) All washes with Buffer C were done 3 times; and (e) bacterially expressed TgEMLC1 and TgELC1 were each added to the flow cells at a concentration of 25 μg/ml. Data were collected on an Eclipse Ti-U inverted microscope (Nikon) equipped with a 100× Plan Apo objective lens (1.49 NA) and a XR/Turbo-Z camera (Stanford Photonics) running Piper Control software (v2.3.39). Data were collected at 10 frames per second. Actin filament tracking and analysis was done in Imaris ×64 v. 7.6.1 (Bitplane AG, Zurich, Switzerland).

### Duty ratio measurements

To compare the duty cycle of WT recombinant TgMyoA with and without KNX-002 treatment, the number of myosin heads capable of interacting with an actin filament was estimated as described previously [20, 74]. Briefly, the *in vitro* motility assay was carried out as described above using various concentrations of myosin treated with either 10 μM KNX-002 or the equivalent amount (0.5% v/v) of DMSO. At each concentrations of myosin on the coverslip, actin filament length and velocity was measured in Imaris ×64 v. 7.6.1 (Bitplane AG, Zurich, Switzerland) and an estimate of the number of heads available to interact with an actin filament was determined. Duty cycle (f) were calculated by plotting filament velocity (V) against number of heads (N). These data were then fit to the equation V= (*a* ×Vmax)×[1−(1−f)^n^] by non-linear regression analysis (GraphPad Prism (Version 9.4.0 (453)) as previously described [74, 75].

### Parasite growth assays

<u>Fluorescence-based growth assay</u>. To quantify growth in the presence of compound(s), 1×10^4^ tachyzoites were preincubated with compound for 5 minutes prior to inoculation onto 3 wells of a black sided, clear bottom 384-well plate containing a confluent monolayer of HFF cells. For compound assays, parasites expressing tdTomato were incubated with vehicle or serial dilutions of compound, maintaining a constant DMSO concentration (0.2% v/v). The fluorescent signal was quantified daily for 7 days (ex 530/25, em 590/35) on a Biotek synergy 2 plate reader. The IC50 of growth inhibitory compounds was quantified using GraphPad Prism. <u>Plaque assay</u>. 10, 100, 200, or 500 parasites were used to inoculate a confluent monolayer of HFF cells in a 12-well plate and incubated for 7 days at 37ºC, 5% CO2. Infected monolayers were then washed with phosphate-buffered saline, fixed with 100% EtOH for five minutes, stained with 5X crystal violet solution (12.5 g crystal violet, 125 ml 100% EtOH, 500 ml 1% w/v ammonium oxalate) for 5 minutes, washed with PBS, and allowed to dry. Wells were imaged using BioTek ImmunoSpot S6 MACRO Analyzer, and plaque number and area were measured using the Analyze Particles tools in Fiji v2.3.0.

### Parasite motility assays

<u>2D motility</u> assays were performed as previously described [76], using monoclonal antibody DG52 (generously provided by Dr. David Sibley) against TgSAG1 for immunofluorescent staining of the trails.

<u>3D motility</u> assays were performed as previously described [16]. Briefly, parasites were syringed released and filtered using a 3-μm Whatman Nuclepore filter (Millipore Sigma) before spinning and resuspending in motility buffer containing 0.5 mg/ml Hoechst 33342. A mixture of 1:3:3 of parasites, motility buffer with compound, and Matrigel was created and pipetted into a flow cell. Flow cells were incubated for 7 minutes at 27ºC and then 3 minutes at 35ºC. Images were captured using a Nikon Eclipse TE300 epifluorescence microscope (Nikon Instruments, Melville, NY) using a 20× PlanApo λ objective and a NanoScanZ piezo Z stage insert (Prior Scientific, Rockland, MA). For the data shown in Figures 4B,C and Suppl. Figure 7, an iXON 885 EMCCD camera (Andor Technology, Belfast, Ireland) driven by NIS Elements software v. 3.20 (Nikon Instruments, Melville, NY) was used to collect time lapse image (502 pixel × 500 pixel) stacks of the fluorescently label parasite nuclei, with Mercury lamp excitation. Images were captured for 60 sec. The camera was set to trigger mode, 2×2 binning, readout speed of 35 MHz, conversion gain of 3.8x, and EM gain setting of 3. For the data shown in Figure 7 and Suppl. Figure 12, 1024 pixel × 384 pixel images were captured using an iXON Life 888 EMCCD camera (Andor Technology) driven by NIS Elements software v.5.11, including the Illumination Sequence acquisition module (Nikon Instruments), and excited using a pE-4000 LED illumination system (CoolLED, Andover England). Images were captured for 80 sec. The camera was set to trigger mode, no binning, readout speed of 30 Mhz, conversion gain of 3.8x, and an EM gain of 300.

All motility assays captured image stacks consisting of 41 z-slices, captured 1 um apart with a 16ms exposure time. The data were analyzed using Imaris ×64 v. 9.2.0 (Bitplane AG, Zurich, Switzerland). The ImarisTrack module tracked the parasite nuclei using an estimated spot volume of 3.0 × 3.0 × 6.0 μm. A maximum distance of 6.0 μm and maximum gap size of 2 were applied to the tracking algorithm. A track duration filter of 10 seconds was used to avoid tracking artifacts and a 2 μm displacement filter was used to eliminate parasites moving by Brownian motion [16].

### Mutagenesis and selection for resistant parasites

The dose of ENU required to kill 70% of parasites [40] during a 4 hour treatment was determined by plaque assay (see above). For each mutagenesis, tdTomato-expressing parasites were liberated from large vacuoles by scraping the monolayer and syringe release through a 26G needle, then treated with ENU at the 70% kill concentration for 4 hours. The parasites were washed 3x in DMEM containing 1% v/v fetal bovine serum to remove residual ENU and inoculated onto a T25 of confluent HFF cells. 24 - 48h post infection, surviving parasites that had lysed their host cells were transferred to fresh HFF cells and grown for until plaques became visible (10-14 days) in the presence of 40 μM KNX-002, which is close to the compound’s estimated EC90 value in the growth assay (37.6μM; 95% C.I. 18.9-136.4 μM). Clonal lines were isolated by serial dilution in five 96-well plates per mutagenesis. Parasite sensitivity to KNX-002 was then determined by fluorescence-based growth assay (see above).

RNA was isolated from resistant clones with the GeneJET RNA purification kit and used to generate cDNA with the Agilent cDNA synthase kit for oligoDT synthesis. cDNA encoding TgMyoA, TgMLC1, TgELC1, TgELC2 was amplified and sequenced with primers covering the entire transcript (Table 1, P6-P20). Sequence alignments were analyzed in SnapGene and sequences containing non-synonymous mutations were compared to other strains to identify naturally occurring polymorphisms unlikely to be responsible for resistance.

### Generation of TgMyoA T130A mutant using CRISPR/Cas9

All oligonucleotides were synthesized by Sigma-Aldrich (The Woodlands, TX), and restriction enzymes were purchased from New England Biolabs (Ipswich, MA). To generate the Cas9eGFPTgMyoAguide plasmid (Suppl. Figure 10), plasmid pSAG1::Cas9-U6::sgUPRT [77] was PCR amplified with primers P1 and P2 (Table 1). This linear pSAG1::Cas9-U6::sgUPRT PCR product contained all features of the original vector, the TgMyoA T130 protospacer sequence (5’-GAATCAGATCTACGCGACTG-3’), and introduced an EcoRI site. The T130A double stranded repair template was produced by annealing oligonucleotides P3 and P4 (Table 1). This repair template contains the T130A mutation and two additional mutations that do not alter the amino acid sequence: a mutation introducing a diagnostic BglII site and a mutation ablating the PAM sequence (Suppl. Figure 10A). RHΔku80Δhxgprt parasites were transfected with the ds repair template and Cas9eGFPTgMyoAguide plasmid (Suppl. Figure 10B). Wild-type RHΔku80Δhxgprt and transfected tachyzoites were syringe released and filtered 2 days post-transfection. eGFP+ parasites were sorted 3-per well onto a 96-well plate in the UVM Flow Cytometry and Cell Sorting Facility using a FACSAria cell sorter (Becton Dickinson Biosciences, Franklin Lakes NJ). Positive clonal populations were validated for GFP expression 7 days post-sorting.

To identify T130A positive mutant clones, gDNA was purified from WT and transfected parasites directly from 96-well plates using previously methods [78]. PCR fragments of approximately the first 3kb of MyoA were generated using the Phire tissue direct PCR Master kit (Thermo Fisher Scientific, Waltham MA) with primers P5 and P11 (Table 1). Diagnostic digests with BglII provided an initial screen for T130A mutant parasites as the ds repair template introduced a third BglII site into the amplified sequence (Suppl. Figure 10C). Positive T130A mutant clones were further verified by harvesting tachyzoite cDNA and sequencing using primers P5-P13 (Table 1).

### Western Blotting

Protein samples were resolved by SDS–polyacrylamide gel electrophoresis and transferred to Immobilon®-FL polyvinylidene fluoride membrane (Millipore Sigma) as previously described [76]. Blots were probed with rabbit anti-TgMyoA antiserum UVT109 (diluted 1:1000) and rabbit anti-TgACT1 (1:10,000; generous gift from Dr. David Sibley). Goat anti-rabbit IRDye800CW (1:20,000; LI-COR Biosciences, Lincoln, NE) was used as a secondary antibody. The blots were scanned using an Odyssey CLx infrared imager (LI-COR) and analyzed using Image Studio software v.2.1.10 (LI-COR).

### Mammalian cell toxicity assays

HFF and HepG2 cells were grown to ∼80% confluence in 384-well plates before media was replaced with DMEM containing 10% v/v fetal bovine serum and serial dilutions of compound or vehicle while maintaining a consistent DMSO concentration (0.2% v/v). Cells were incubated with compound at 37°C for 72h before multiplexed quantitation of toxicity using CellTox green™ and cell viability using Cell-TiterGlo® (both assay kits from Promega Corp, Madison WI). As a positive control for toxicity 24h prior to cessation of the assay 0.5 mM NaN3 was added to half of the vehicle wells to induce cell death. <u>CellTox green</u>. To quantify dead, dying and damaged cells in each well, an equal volume (25μl) of CellTox green reagent (1:500 CellTox green dye in CellTox green assay buffer) was added to each well and mixed on an orbital shaker (700 - 900rpm) for 1 minute. The plate was equilibrated at room temperature for 15 minutes before quantifying the fluorescence signal (ex 485/20, em 528/20) using a Biotek Synergy 2 plate reader. <u>CellTiter-Glo.</u> Following quantification of toxicity, the number of viable cells/well was quantified by adding 25μl CellTiter-Glo reagent to each well. To maximize cell lysis plates were mixed on an orbital shaker (700 - 900rpm) for 5 minutes before equilibration at room temperature for an additional 10 minutes. Quantitation was achieved by measuring luminescence per well using a Biotek Synergy 2 plate reader.

### Parasite replication assay

Confluent HFF monolayers grown in clear bottomed black-sided 96-well plates were challenged with 1×10^4^ tachyzoites and incubated for 1 hour at 37°C with 5% CO2. Monolayers were then washed once with DMEM containing 1% v/v fetal bovine serum before incubating with 0 (DMSO vehicle only) or compound. Monolayers were fixed with 2% v/v PFA 12h, 24h and 36h after compound addition. After blocking and permeabilization in 0.2% v/v Triton X-100 and 1% w/v bovine serum albumin, samples were incubated with Mab 45.36 against TgIMC1 for 2 hours (3.7 μg/ml) at 23ºC. Monolayers were washed three times with blocking buffer before incubation in the dark with a 1:1000 dilution of Alexa546-conjugated anti-rabbit IgG (ThermoFisher) for 1 hour at 23ºC. The number in parasites in each of 50 vacuoles per sample were counted blind.

### Parasite toxicity assays

<u>Extracellular parasite toxicity</u>. 1×10^7^ parasites in DMEM containing 1% v/v fetal bovine serum were mixed in a 96-well plate with 0 (DMSO vehicle only), 40, or 80μM KNX-002, in a final volume of 50μl. The plate was incubated for 1 or 3 hours at 37°C with 5% CO2. 50μl CellTiter-Glo reagent was added to each well and mixed on an orbital shaker for 5 minutes (500 – 700rpm), followed by a 15-minute incubation at 23ºC. The relative number of viable parasites was quantified by luminescence, read on a Biotek Synergy 2 plate reader. <u>Intracellular parasite toxicity</u> in the presence of various concentrations of compound was determined using the replication assay described above.

### ATPase assays

ATPase assays for the SAR studies were performed per manufacturer’s recommendations using the ADP-Glo Kinase Assay (Promega Corp, Madison WI). Briefly, 8 μl kinase reactions were performed by combining 5 μl of myosin (0.128 mg/ml), 1 μl of actin (4.8 mg/ml), 1 μl of ATP (8 mM) and 1 μl of 1X Kinase Reaction Buffer (40 mM Tris pH 7.6, 20 mM MgCl2, and 0.1 mg/ml BSA). After a 90 minute incubation at room temperature, 5 ul of ADP-Glo Reagent was added to the kinase reaction followed by a 40 minute incubation at room temperature. Finally, 10 ul of Kinase Detection Reagent was added and after a 60 minute incubation, ATP production was detected by measuring luminescence with a Synergy H1 Hybrid plate reader. For compound inhibitor experiments, the control reactions contained 0.5% v/v DMSO and compounds were diluted in 1X Kinase Reaction Buffer. All assays were performed under conditions where ADP generation (measured in relative luminescence units, RLU) was linear with respect to both TgMyoA concentration and incubation time. Reactions were performed in duplicate on each plate and all experiments were replicated three times. Data were analyzed using GraphPad Prism.

### Size-exclusion Chromatography

Twin-strep-II (TST)-tagged TgMyoA (1-778) (ToxoDB ID TGGT1_070410/GenBank™ accession number EPR60343.1) wild-type and T130A mutant were expressed in *Trichoplusia ni* (Tni) insect cells. Protein was purified by streptactin affinity, and finally SEC, using a final buffer containing 20 mM HEPES pH 7.5, 150 mM NaCl, and 1 mM DTT. Prior to DSF experiments, purified, concentrated TgMyoA (1-778) was diluted with buffer containing MgCl2, ADP, NaF, and Al(NO3)3, to final concentrations of 2 mM MgCl2, 2 mM ADP, 6 mM NaF, 1.75 mM Al(NO3)3.

### Differential Scanning Fluorimetry

Purified TgMyoA (1-778) WT and T130A were diluted to 1.0 mg/mL in buffer containing 150 mM NaCl; 20mM HEPES, pH 7.5; 1 mM MgCl2; 1 mM ADP; 6 mM NaF; 1.75 mM Al(NO3)3; 1 mM DTT; and SYPRO® orange dye at a final concentration of 5X. 20 μL reactions were tested in triplicate using the StepOnePlus™ Real-Time PCR System from ThermoFisher, with all replicate samples obtained from a single preparation of MyoA protein.

### Mouse infections

Infections were carried out in 4-5 week old male CBA/J mice purchased from Jackson Laboratories. All mice were socially housed and acclimated for 3 days prior to infections. For virulence studies a total of nine mice (three biological replicates of three mice each) were challenged by intraperitoneal injection of either 50 or 1000 wild-type (RH) or T130A parasites resuspended in a total volume of 200μl sterile PBS. Daily animal health monitoring began two days after infection. Mice were euthanized when predetermined welfare thresholds were exceeded.

For KNX-002 treatment studies, 15 mice were injected intraperitoneally with either 20mg/kg KNX-002 (40mM compound stock in DMSO diluted into PBS containing 30% v/v polyethylene glycol 400) or an equivalent amount of DMSO (vehicle) in the same diluent. 20-30 minutes after compound/vehicle injection, ten compound-treated mice were intraperitoneally infected with either 500 wild-type or 500 T130A mutant parasites, and five vehicle-treated mice were similarly infected with either wild-type or T130A mutant parasites. A second intraperitoneal injection of 20mg/kg KNX-002 or DMSO was given two days later. The rationale for this dosing schedule was two-fold: (a) We expected the compound to work primarily on egressing and extracellular parasites. Since the parasite’s lytic cycle takes approximately 48 hours, this two-day injection schedule was chosen to maximize exposure of the extracellular parasites to freshly injected compound early in establishment of the infection. (b) A preliminary KNX-002 dose tolerance study was conducted prior to the infection experiments. Two doses of 20mg/kg (the highest dose tested) administered two days apart revealed minor hepatoxicity without signs of excessive pain or distress and was therefore considered the maximal tolerated dose.

Daily animal health monitoring began two days after infection, and mice were euthanized when predetermined welfare thresholds were exceeded. All animal experiments were carried out under University of Vermont Institutional Animal Care and Use Committee protocol #PROTO202000126.

### RNA sequencing

For each sample, three confluent flasks of HFFs cells were inoculated with either wild-type or T130A mutant parasites. Two days after infection, when vacuoles were large (≳64 parasites/vacuole), parasites were harvested by scraping the flask and passing the suspension three times through a 26G needle. Released parasites were incubated at 37°C for 1 hr, and RNA was then isolated using the GeneJET RNA Purification kit (Thermo-Fisher) following manufacturer instructions. For each sample, 1μg of total RNA (A260/280 = 1.8-2.2, A260/230 ≥ 1.8) was sent to Novogene for library preparation and sequencing to a read depth of 20 million paired end reads per sample. Quality control and sequence alignment to the *Toxoplasma* genome (toxodb.org) was done in house. The most differentially expressed genes were identified from all genes expressed in all samples, with a FPKM > 0.1 and an adjusted p-value of < 0.05. Expression of known TgMyoA interacting proteins, glideosome components, actin and actin regulatory factors were also separately quantified where detected in all three replicates of both the wildtype and T130A mutant. TgELC2 (TGME49_305050) was not consistently expressed and therefore was excluded from the analysis shown in Suppl. Figure 14.

### Compound synthesis and characterization

Details of compound synthesis and analysis are presented in Supplementary Data. Mass spectroscopic (m/z) data were obtained by Mrs. Caroline Horsburgh at the University of St Andrews mass spectrometry service. Electrospray ionization techniques were applied and a ThermoFisher LTQ Orbitrap XL spectrometer was used. Values are quoted as a ratio of mass to charge (m/z) in Daltons [Da]. NMR spectra were recorded on either a Bruker Ascend 500 (^1^H 600; ^13^C 150 MHz) or a Bruker Advance II 400 (^1^H 400; ^13^C 101 MHz). ^13^C NMR spectra were taken with a DEPTQ pulse sequence. Spectra were recorded using deuterated solvents. For ^1^H NMR analysis, multiplicities of signals are denoted as follows: s (singlet), d (doublet), t (triplet), q (quartet), dd (doublet of doublets), dt (doublet of triplets), m (multiplet). Coupling constants (J) are quoted to the nearest 0.1 Hz. Chemical shifts (δ) are stated in ppm and referenced to residual solvent signals (CHCl3 7.260 ppm (s), 77.160 ppm (t); (CD3)2SO 2.50 ppm (s) or 39.52 ppm (pentet)). IR spectra were obtained using a Shimadzu IRAffinity 1S IR Spectrometer as ATR. The IR data were processed using OriginPro 2022 v9.9.0.225. Optical rotations were determined using a Perkin Elmer Model 341 Polarimeter with a Na/Hal lamp (Na D line, 589 nm). Melting points were measured using capillary tubes on an Electrothermal 9100 melting point apparatus. Values are quoted in ranges and to 1.0 °C.

## Supporting information

Suppl information_Compound synthesis and analysis

## Acknowledgements

This work was supported by U.S. Public Health Service (USPHS) grants AI139201 and AI137767 (GEW) and GM141743 (DMW), and by Canadian Institutes of Health Research Grant 148596 (MJB), as well as fellowships from the USPHS (F31AI145214; RVS) and American Heart Association (20POST35220017; RSK). AS was supported by USPHS training grant T32AI055402. MJB gratefully acknowledges the Canada Research Chair program for salary support. We thank Cytokinetics, Inc., for their willingness to provide staff time and reagents to this project, Samantha Previs for technical support, Dr. Shane Nelson for help with the duty ratio analysis, and Dr. Roxanna del Rio-Guerra for her flow cytometry advice and assistance. Genomic-scale data sets and ancillary information were obtained from the Toxoplasma Genome Database (ToxoDB.org). ToxoDB is a component of the Eukaryotic Pathogen, Vector and Host Informatics Resource [79], supported by the National Institutes of Allergy and Infectious Diseases and the Wellcome Trust; we gratefully acknowledge the staff responsible for developing and maintaining this resource.

The authors declare that they have no conflicts of interest.

## Notes

### Competing Interest Statement

The authors have declared no competing interest.

### Summary of Updates

The manuscript was submitted to Review Commons and has been revised in response to comments from the reviewers. Revisions include: the addition of 2 new figures (Suppl. Figures 2 and 12) and accompanying text; additional statistical analysis of the data in Figure 7; additional rationale for the animal treatment protocol; and other minor changes that provided additional experimental detail, expanded the discussion as suggested by the reviewers, or were made to improve clarity or correct errors in the previous version.

## References

1. Dubey JP, Murata FHA, Cerqueira-Cézar CK, Kwok OCH, Villena I. Congenital toxoplasmosis in humans: an update of worldwide rate of congenital infections. Parasitology. 2021:1-11. Epub 2021/07/14. doi: 10.1017/s0031182021001013. PubMed PMID: 34254575.

2. El Bissati K, Levigne P, Lykins J, Adlaoui EB, Barkat A, Berraho A, et al. Global initiative for congenital toxoplasmosis: an observational and international comparative clinical analysis. Emerg Microbes Infect. 2018;7(1):165. Epub 2018/09/29. doi: 10.1038/s41426-018-0164-4. PubMed PMID: 30262847; PubMed Central PMCID: PMCPMC6160433.

3. Vidal JE, Hernandez AV, de Oliveira AC, Dauar RF, Barbosa SP, Jr., Focaccia R. Cerebral toxoplasmosis in HIV-positive patients in Brazil: clinical features and predictors of treatment response in the HAART era. AIDS Patient Care STDS. 2005;19(10):626-34. Epub 2005/10/20. doi: 10.1089/apc.2005.19.626. PubMed PMID: 16232047.

4. Porter SB, Sande MA. Toxoplasmosis of the central nervous system in the acquired immunodeficiency syndrome. N Engl J Med. 1992;327(23):1643-8. Epub 1992/12/03. doi: 10.1056/NEJM199212033272306. PubMed PMID: 1359410.

5. de Oliveira GB, da Silva MA, Wanderley LB, da Cunha Correia C, Ferreira EC, de Medeiros ZM, et al. Cerebral toxoplasmosis in patients with acquired immune deficiency syndrome in the neurological emergency department of a tertiary hospital. Clin Neurol Neurosurg. 2016;150:23–6. doi: 10.1016/j.clineuro.2016.08.014. PubMed PMID: 27573702.

6. Gandhi RT. Toxoplasmosis in HIV-infected patients. UpToDate. 2015.

7. http://www.unaids.org/en/resources/documents/2017/UNAIDS_FactSheet. Fact Sheet - Latest Global and Regional Statistics on the Status of the AIDS Epidemic Communications and Global Advocacy [Internet]. 2017 July 20, 2017.

8. Ben-Harari RR, Goodwin E, Casoy J. Adverse Event Profile of Pyrimethamine-Based Therapy in Toxoplasmosis: A Systematic Review. Drugs R D. 2017;17(4):523–44. doi: 10.1007/s40268-017-0206-8. PubMed PMID: 28879584; PubMed Central PMCID: PMCPMC5694419.

9. Choquet-Kastylevsky G, Vial T, Descotes J. Allergic adverse reactions to sulfonamides. Curr Allergy Asthma Rep. 2002;2(1):16–25. PubMed PMID: 11895621.

10. McLeod R, Khan AR, Noble GA, Latkany P, Jalbrzikowski J, Boyer K, et al. Severe sulfadiazine hypersensitivity in a child with reactivated congenital toxoplasmic chorioretinitis. Pediatr Infect Dis J. 2006;25(3):270–2. doi: 10.1097/01.inf.0000202070.59190.9a. PubMed PMID: 16511396.

11. McPhillie M, Zhou Y, El Bissati K, Dubey J, Lorenzi H, Capper M, et al. New paradigms for understanding and step changes in treating active and chronic, persistent apicomplexan infections. Scientific reports. 2016;6:29179. doi: 10.1038/srep29179. PubMed PMID: 27412848; PubMed Central PMCID: PMCPMC4944145.

12. Barragan A, Sibley LD. Transepithelial migration of Toxoplasma gondii is linked to parasite motility and virulence. J Exp Med. 2002;195(12):1625–33. PubMed PMID: 12070289.

13. Furtado JM, Bharadwaj AS, Ashander LM, Olivas A, Smith JR. Migration of Toxoplasma gondii-infected dendritic cells across human retinal vascular endothelium. Invest Ophthalmol Vis Sci. 2012;53(11):6856-62. Epub 2012/09/07. doi: 10.1167/iovs.12-10384. PubMed PMID: 22952125; PubMed Central PMCID: PMC3466064.

14. Sibley LD. Intracellular parasite invasion strategies. Science. 2004;304(5668):248–53. PubMed PMID: 15073368.

15. Harker KS, Ueno N, Lodoen MB. Toxoplasma gondii Dissemination: A Parasite’s Journey through the Infected Host. Parasite Immunol. 2015;37:141-9. Epub 2014/11/20. doi: 10.1111/pim.12163. PubMed PMID: 25408224.

16. Leung JM, Rould MA, Konradt C, Hunter CA, Ward GE. Disruption of TgPHIL1 alters specific parameters of Toxoplasma gondii motility measured in a quantitative, three-dimensional live motility assay. PLOS One. 2014;9(1):e85763. Epub 2014/02/04. doi: 10.1371/journal.pone.0085763. PubMed PMID: 24489670; PubMed Central PMCID: PMC3906025.

17. Herm-Gotz A, Weiss S, Stratmann R, Fujita-Becker S, Ruff C, Meyhofer E, et al. Toxoplasma gondii myosin A and its light chain: a fast, single-headed, plus-end-directed motor. EMBO J. 2002;21(9):2149–58.

18. Nebl T, Prieto JH, Kapp E, Smith BJ, Williams MJ, Yates JR, 3rd, et al. Quantitative in vivo analyses reveal calcium-dependent phosphorylation sites and identifies a novel component of the Toxoplasma invasion motor complex. PLOS Pathog. 2011;7(9):e1002222. PubMed PMID: 21980283.

19. Williams MJ, Alonso H, Enciso M, Egarter S, Sheiner L, Meissner M, et al. Two Essential Light Chains Regulate the MyoA Lever Arm To Promote Toxoplasma Gliding Motility. MBio. 2015;6(5). Epub 2015/09/17. doi: 10.1128/mBio.00845-15. PubMed PMID: 26374117.

20. Heaslip AT, Leung JM, Carey KL, Catti F, Warshaw DM, Westwood NJ, et al. A small-molecule inhibitor of T. gondii motility induces the posttranslational modification of myosin light chain-1 and inhibits myosin motor activity. PLOS Pathog. 2010;6(1):e1000720. doi: 10.1371/journal.ppat.1000720. PubMed PMID: 20084115.

21. Leung JM, Tran F, Pathak RB, Poupart S, Heaslip A, Ballif BA, et al. Identification of T. gondii myosin light chain-1 as a direct target of tachypleginA-2, a small-molecule inhibitor of parasite motility and invasion. PLOS One. 2014;9(6):e98056. doi: 10.1371/journal.pone.0098056. PubMed Central PMCID: PMC4043638.

22. Bement WM, Mooseker MS. TEDS rule: a molecular rationale for differential regulation of myosins by phosphorylation of the heavy chain head. Cell Motil Cytoskeleton. 1995;31(2):87-92. Epub 1995/01/01. doi: 10.1002/cm.970310202. PubMed PMID: 7553910.

23. Heintzelman MB, Schwartzman JD. A novel class of unconventional myosins from Toxoplasma gondii. J Mol Biol. 1997;271(1):139–46.

24. Powell CJ, Ramaswamy R, Kelsen A, Hamelin DJ, Warshaw DM, Bosch J, et al. Structural and mechanistic insights into the function of the unconventional class XIV myosin MyoA from Toxoplasma gondii. Proc Natl Acad Sci U S A. 2018;115(45):E10548-e55. Epub 2018/10/24. doi: 10.1073/pnas.1811167115. PubMed PMID: 30348763; PubMed Central PMCID: PMCPMC6233092.

25. Trivedi DV, Nag S, Spudich A, Ruppel KM, Spudich JA. The Myosin Family of Mechanoenzymes: From Mechanisms to Therapeutic Approaches. Annu Rev Biochem. 2020;89:667-93. Epub 2020/03/15. doi: 10.1146/annurev-biochem-011520-105234. PubMed PMID: 32169021.

26. Egarter S, Andenmatten N, Jackson AJ, Whitelaw JA, Pall G, Black JA, et al. The Toxoplasma Acto-MyoA motor complex is important but not essential for gliding motility and host cell invasion. PLOS One. 2014;9(3):e91819. Epub 2014/03/19. doi: 10.1371/journal.pone.0091819. PubMed PMID: 24632839; PubMed Central PMCID: PMC3954763.

27. Andenmatten N, Egarter S, Jackson AJ, Jullien N, Herman JP, Meissner M. Conditional genome engineering in Toxoplasma gondii uncovers alternative invasion mechanisms. Nat Methods. 2012;10(2):125-7. Epub 2012/12/25. doi: 10.1038/nmeth.2301. PubMed PMID: 23263690; PubMed Central PMCID: PMC3605914.

28. Meissner M, Schluter D, Soldati D. Role of Toxoplasma gondii myosin A in powering parasite gliding and host cell invasion. Science. 2002;298(5594):837–40. PubMed PMID: 12399593.

29. Bichet M, Touquet B, Gonzalez V, Florent I, Meissner M, Tardieux I. Genetic impairment of parasite myosin motors uncovers the contribution of host cell membrane dynamics to Toxoplasma invasion forces. BMC biology. 2016;14(1):97. doi: 10.1186/s12915-016-0316-8. PubMed PMID: 27829452; PubMed Central PMCID: PMCPMC5101828.

30. Frenal K, Marq JB, Jacot D, Polonais V, Soldati-Favre D. Plasticity between MyoC-and MyoA-glideosomes: an example of functional compensation in Toxoplasma gondii invasion. PLOS Pathog. 2014;10(10):e1004504. Epub 2014/11/14. doi: 10.1371/journal.ppat.1004504. PubMed PMID: 25393004; PubMed Central PMCID: PMC4231161.

31. Frenal K, Soldati-Favre D. Plasticity and Redundancy in Proteins Important for Toxoplasma Invasion. PLOS Pathog. 2015;11(8):e1005069. Epub 2015/08/14. doi: 10.1371/journal.ppat.1005069. PubMed PMID: 26270966; PubMed Central PMCID: PMC4536160.

32. Lamarque MH, Roques M, Kong-Hap M, Tonkin ML, Rugarabamu G, Marq JB, et al. Plasticity and redundancy among AMA-RON pairs ensure host cell entry of Toxoplasma parasites. Nat Commun. 2014;5:4098. Epub 2014/06/18. doi: 10.1038/ncomms5098. PubMed PMID: 24934579.

33. Bookwalter CS, Kelsen A, Leung JM, Ward GE, Trybus KM. A Toxoplasma gondii class XIV myosin, expressed in Sf9 with a parasite co-chaperone, requires two light chains for fast motility. J Biol Chem. 2014;289(44):30832–41. doi: 10.1074/jbc.M114.572453.

34. Harding CR, Meissner M. The inner membrane complex through development of Toxoplasma gondii and Plasmodium. Cell Microbiol. 2014;16(5):632-41. Epub 2014/03/13. doi: 10.1111/cmi.12285. PubMed PMID: 24612102.

35. Kono M, Prusty D, Parkinson J, Gilberger TW. The apicomplexan inner membrane complex. Front Biosci (Landmark Ed). 2013;18:982-92. Epub 2013/06/12. PubMed PMID: 23747861.

36. Gubbels MJ, Li C, Striepen B. High-Throughput Growth Assay for Toxoplasma gondii Using Yellow Fluorescent Protein. Antimicrob Agents Chemother. 2003;47(1):309–16. PubMed PMID: 12499207.

37. Mersch-Sundermann V, Knasmüller S, Wu XJ, Darroudi F, Kassie F. Use of a human-derived liver cell line for the detection of cytoprotective, antigenotoxic and cogenotoxic agents. Toxicology. 2004;198(1-3):329-40. Epub 2004/05/13. doi: 10.1016/j.tox.2004.02.009. PubMed PMID: 15138059.

38. Dobrowolski JM, Carruthers VB, Sibley LD. Participation of myosin in gliding motility and host cell invasion by Toxoplasma gondii. Mol Microbiol. 1997;26(1):163–73.

39. Hakansson S, Morisaki H, Heuser J, Sibley LD. Time-lapse video microscopy of gliding motility in Toxoplasma gondii reveals a novel, biphasic mechanism of cell locomotion. Mol Biol Cell. 1999;10(11):3539–47.

40. Farrell A, Coleman BI, Benenati B, Brown KM, Blader IJ, Marth GT, et al. Whole genome profiling of spontaneous and chemically induced mutations in Toxoplasma gondii. BMC Genomics. 2014;15:354. Epub 2014/06/03. doi: 10.1186/1471-2164-15-354. PubMed PMID: 24885922; PubMed Central PMCID: PMC4035079.

41. Work SS, Warshaw DM. Computer-assisted tracking of actin filament motility. Anal Biochem. 1992;202(2):275–85. PubMed PMID: 1519753.

42. Gras S, Jimenez-Ruiz E, Klinger CM, Schneider K, Klingl A, Lemgruber L, et al. An endocytic-secretory cycle participates in Toxoplasma gondii in motility. PLOS Biol. 2019;17(6):e3000060. Epub 2019/06/25. doi: 10.1371/journal.pbio.3000060. PubMed PMID: 31233488.

43. Meissner M, Ferguson DJ, Frischknecht F. Invasion factors of apicomplexan parasites: essential or redundant? Curr Opin Microbiol. 2013;16(4):438-44. Epub 2013/06/04. doi: 10.1016/j.mib.2013.05.002. PubMed PMID: 23727286.

44. Rompikuntal PK, Kent RS, Foe IT, Deng B, Bogyo M, Ward GE. Blocking Palmitoylation of Toxoplasma gondii Myosin Light Chain 1 Disrupts Glideosome Composition but Has Little Impact on Parasite Motility. mSphere. 2021;6(3). Epub 2021/05/21. doi: 10.1128/mSphere.00823-20. PubMed PMID: 34011689.

45. Tardieux I, Baum J. Reassessing the mechanics of parasite motility and host-cell invasion. J Cell Biol. 2016;214(5):507–15. doi: 10.1083/jcb.201605100. PubMed PMID: 27573462; PubMed Central PMCID: PMCPMC5004448.

46. Whitelaw JA, Latorre-Barragan F, Gras S, Pall GS, Leung JM, Heaslip A, et al. Surface attachment, promoted by the actomyosin system of Toxoplasma gondii is important for efficient gliding motility and invasion. BMC biology. 2017;15(1):1. doi: 10.1186/s12915-016-0343-5. PubMed PMID: 28100223; PubMed Central PMCID: PMCPMC5242020.

47. Pavlou G, Touquet B, Vigetti L, Renesto P, Bougdour A, Debarre D, et al. Coupling Polar Adhesion with Traction, Spring and Torque Forces Allows High Speed Helical Migration of the Protozoan Parasite Toxoplasma. ACS nano. 2020;14:7121-39. Epub 2020/05/21. doi: 10.1021/acsnano.0c01893. PubMed PMID: 32432851.

48. Frenal K, Polonais V, Marq JB, Stratmann R, Limenitakis J, Soldati-Favre D. Functional dissection of the apicomplexan glideosome molecular architecture. Cell Host Microbe. 2010;8(4):343–57. PubMed PMID: 20951968.

49. Bogyo M, Ward GE. Toxoplasma gondii Chemical Biology. In: Weiss LM, Kim K, editors. Toxoplasma gondii -The Model Apicomplexan: Perspectives and Methods. Second ed. London, UK: Academic Press; 2014. p. 707–30.

50. Ward GE, Carey KL, Westwood NJ. Using small molecules to study big questions in cellular microbiology. Cell Microbiol. 2002;4(8):471–82. doi: 10.1046/j.1462-5822.2002.00205.x. PubMed PMID: 12174082.

51. Bond LM, Tumbarello DA, Kendrick-Jones J, Buss F. Small-molecule inhibitors of myosin proteins. Future Med Chem. 2013;5(1):41-52. Epub 2012/12/22. doi: 10.4155/fmc.12.185. PubMed PMID: 23256812.

52. Ostap EM. 2,3-Butanedione monoxime (BDM) as a myosin inhibitor. J Muscle Res Cell Motil. 2002;23(4):305–8. PubMed PMID: 12630704.

53. Kortagere S, Mui E, McLeod R, Welsh WJ. Rapid discovery of inhibitors of Toxoplasma gondii using hybrid structure-based computational approach. J Comput Aided Mol Des. 2011;25(5):403–11. PubMed PMID: 21359560.

54. Moussaoui D, Robblee JP, Auguin D, Fisher F, Fagnant PM, Macfarlane JE, et al. Mechanism of small molecule inhibition of Plasmodium falciparum myosin A informs antimalarial drug design. bioRxiv. 2022:2022.09.09.507123. doi: 10.1101/2022.09.09.507123.

55. Trivedi DV, Karabina A, Bergnes G, Racca A, Wander H, Jung S, et al. A small-molecule myosin inhibitor as a targeted multi-stage antimalarial. bioRxiv. 2022:2022.09.09.507317. doi: 10.1101/2022.09.09.507317.

56. Istvan ES, Dharia NV, Bopp SE, Gluzman I, Winzeler EA, Goldberg DE. Validation of isoleucine utilization targets in Plasmodium falciparum. Proc Natl Acad Sci U S A. 2011;108(4):1627–32. PubMed PMID: 21205898.

57. Rosenberg A, Luth MR, Winzeler EA, Behnke M, Sibley LD. Evolution of resistance in vitro reveals mechanisms of artemisinin activity in Toxoplasma gondii. Proc Natl Acad Sci U S A. 2019. Epub 2019/12/07. doi: 10.1073/pnas.1914732116. PubMed PMID: 31806760.

58. Wu W, Herrera Z, Ebert D, Baska K, Cho SH, DeRisi JL, et al. A chemical rescue screen identifies a Plasmodium falciparum apicoplast inhibitor targeting MEP isoprenoid precursor biosynthesis. Antimicrob Agents Chemother. 2015;59(1):356-64. Epub 2014/11/05. doi: 10.1128/aac.03342-14. PubMed PMID: 25367906; PubMed Central PMCID: PMCPMC4291372.

59. Preller M, Manstein DJ. Myosin structure, allostery, and mechano-chemistry. Structure. 2013;21(11):1911-22. Epub 2013/11/12. doi: 10.1016/j.str.2013.09.015. PubMed PMID: 24210227.

60. Allingham JS, Smith R, Rayment I. The structural basis of blebbistatin inhibition and specificity for myosin II. Nat Struct Mol Biol. 2005;12(4):378–9. PubMed PMID: 15750603.

61. Duxbury MS, Ashley SW, Whang EE. Inhibition of pancreatic adenocarcinoma cellular invasiveness by blebbistatin: a novel myosin II inhibitor. Biochem Biophys Res Commun. 2004;313(4):992–7. PubMed PMID: 14706640.

62. Mikulich A, Kavaliauskiene S, Juzenas P. Blebbistatin, a myosin inhibitor, is phototoxic to human cancer cells under exposure to blue light. Biochim Biophys Acta. 2012;1820(7):870–7. doi: 10.1016/j.bbagen.2012.04.003. PubMed PMID: 22507270.

63. Barkhuff WD, Gilk SD, Whitmarsh R, Tilley LD, Hunter C, Ward GE. Targeted disruption of TgPhIL1 in Toxoplasma gondii results in altered parasite morphology and fitness. PLOS One. 2011;6(8):e23977. Epub 2011/09/09. doi: 10.1371/journal.pone.0023977. PubMed PMID: 21901148; PubMed Central PMCID: PMC3162014.

64. Green EM, Wakimoto H, Anderson RL, Evanchik MJ, Gorham JM, Harrison BC, et al. A small-molecule inhibitor of sarcomere contractility suppresses hypertrophic cardiomyopathy in mice. Science. 2016;351(6273):617–21. doi: 10.1126/science.aad3456. PubMed PMID: 26912705; PubMed Central PMCID: PMCPMC4784435.

65. Kortagere S, Welsh WJ, Morrisey JM, Daly T, Ejigiri I, Sinnis P, et al. Structure-based design of novel small-molecule inhibitors of Plasmodium falciparum. J Chem Inf Model. 2010;50(5):840–9. PubMed PMID: 20426475.

66. Bosch J, Turley S, Daly TM, Bogh SM, Villasmil ML, Roach C, et al. Structure of the MTIP-MyoA complex, a key component of the malaria parasite invasion motor. Proc Natl Acad Sci U S A. 2006;103(13):4852–7. PubMed PMID: 16547135.

67. Thomas JC, Green JL, Howson RI, Simpson P, Moss DK, Martin SR, et al. Interaction and dynamics of the Plasmodium falciparum MTIP-MyoA complex, a key component of the invasion motor in the malaria parasite. Mol Biosyst. 2010;6(3):494–8. PubMed PMID: 20174678.

68. Mueller C, Graindorge A, Soldati-Favre D. Functions of myosin motors tailored for parasitism. Curr Opin Microbiol. 2017;40:113–22. doi: 10.1016/j.mib.2017.11.003. PubMed PMID: 29161623.

69. Roos DS, Donald RG, Morrissette NS, Moulton AL. Molecular tools for genetic dissection of the protozoan parasite Toxoplasma gondii. Methods in Cell Biology. 451994. p. 27–63.

70. Robert-Paganin J, Robblee JP, Auguin D, Blake TCA, Bookwalter CS, Krementsova EB, et al. Plasmodium myosin A drives parasite invasion by an atypical force generating mechanism. Nat Commun. 2019;10(1):3286. Epub 2019/07/25. doi: 10.1038/s41467-019-11120-0. PubMed PMID: 31337750; PubMed Central PMCID: PMCPMC6650474.

71. Lionne C, Iorga B, Candau R, Travers F. Why choose myofibrils to study muscle myosin ATPase? J Muscle Res Cell Motil. 2003;24(2-3):139–48. Epub 2003/11/12. doi: 10.1023/a:1026045328949. PubMed PMID: 14609025.

72. Hwee DT, Cheng AJ, Hartman JJ, Hinken AC, Lee K, Durham N, et al. The Ca(2+) sensitizer CK-2066260 increases myofibrillar Ca(2+) sensitivity and submaximal force selectively in fast skeletal muscle. J Physiol. 2017;595(5):1657-70. Epub 2016/11/22. doi: 10.1113/jp273248. PubMed PMID: 27869319; PubMed Central PMCID: PMCPMC5330873.

73. Sirigu S, Hartman JJ, Planelles-Herrero VJ, Ropars V, Clancy S, Wang X, et al. Highly selective inhibition of myosin motors provides the basis of potential therapeutic application. Proc Natl Acad Sci U S A. 2016;113(47):E7448-e55. Epub 2016/11/07. doi: 10.1073/pnas.1609342113. PubMed PMID: 27815532; PubMed Central PMCID: PMCPMC5127359.

74. Harris DE, Warshaw DM. Smooth and skeletal muscle myosin both exhibit low duty cycles at zero load in vitro. J Biol Chem. 1993;268(20):14764-8. Epub 1993/07/15. PubMed PMID: 8325853.

75. Walcott S, Fagnant PM, Trybus KM, Warshaw DM. Smooth muscle heavy meromyosin phosphorylated on one of its two heads supports force and motion. J Biol Chem. 2009;284(27):18244–51. PubMed PMID: 19419961.

76. Carey KL, Westwood NJ, Mitchison TJ, Ward GE. A small-molecule approach to studying invasive mechanisms of Toxoplasma gondii. Proc Natl Acad Sci U S A. 2004;101(19):7433–8. doi: 10.1073/pnas.0307769101. PubMed PMID: 15123807.

77. Shen B, Brown KM, Lee TD, Sibley LD. Efficient gene disruption in diverse strains of Toxoplasma gondii using CRISPR/CAS9. MBio. 2014;5(3):e01114-14. Epub 2014/05/16. doi: 10.1128/mBio.01114-14. PubMed PMID: 24825012; PubMed Central PMCID: PMC4030483.

78. Piro F, Carruthers VB, Di Cristina M. PCR Screening of Toxoplasma gondii Single Clones Directly from 96-Well Plates Without DNA Purification. Methods Mol Biol. 2020;2071:117-23. Epub 2019/11/24. doi: 10.1007/978-1-4939-9857-9_6. PubMed PMID: 31758449.

79. Amos B, Aurrecoechea C, Barba M, Barreto A, Basenko EY, Bażant W, et al. VEuPathDB: the eukaryotic pathogen, vector and host bioinformatics resource center. Nucleic Acids Res. 2022;50(D1):D898–d911. Epub 2021/11/01. doi: 10.1093/nar/gkab929. PubMed PMID: 34718728; PubMed Central PMCID: PMCPMC8728164.

