## Supplementary material for "The class XIV myosin of *Toxoplasma gondii*, TgMyoA, is druggable in an animal model of infection": Suppl information_Compound synthesis and analysis

**Supplemental information: Compound synthesis and analysis**

General procedure for synthesis of semicarbazones (General Procedure **1**)

To a stirring suspension of semicarbazide hydrochloride (5.00g, 44.8 mmol, 1.00 eq.) in ethanol (50 mL), anhydrous sodium acetate (4.78g, 58.3 mmol, 1.30 eq.) was added. The resulting suspension was heated to reflux and stirred for 1 hour. The suspension was filtered whilst hot and the filtrate was added to a new flask. Ketone (40.4 mmol, 0.90 eq.) was added to the filtrate. The reaction was heated to reflux and stirred for 2 hours. The reaction was left to cool to room temperature and left to stand overnight. The resulting suspension was filtered and the solid was washed with cold ethanol (3 x 50 mL). The solid was collected and dried under reduced pressure in a vacuum oven at 60 °C to yield product with no further purification required.

General procedure for synthesis of 4-formyl-pyrazoles (General Procedure **2**)

A round bottomed flask charged with DMF (2.00 eq.) was cooled to 0 °C before the dropwise addition of phosphorus oxychloride (2.30 eq.). Once addition was complete, the solution was warmed to room temperature and stirred for 20 minutes. The solution was cooled to 0 °C and semicarbazone (1.00 eq.) was added in portions. The reaction was heated to 80 °C and stirred for 2 hours. The reaction was poured onto ice whilst hot. Once the ice had melted, the mixture was treated with 30% aqueous solution of sodium hydroxide until a pH of 8-9 was obtained. The resulting suspension was isolated by filtration and the filtrate was treated with conc. hydrochloric acid until a pH of 6-7 was obtained and the reaction left to stand overnight. The resulting suspension was filtered and the solid was washed with ice water (3 x 50 mL). The solid was collected and dried under reduced pressure in a vacuum oven to yield product with no further purification required.

General procedure for Szymoniak-Kulinkovich reaction [1] (General Procedure **3**)

To a flame dried flask, a solution of nitrile (1.00 eq.) in diethyl ether (5 mL/mmol, pre-dried over 4Å molecular sieves) was added. The solution was cooled to – 78 °C before the addition of titanium isopropoxide (1.10 eq.), followed by a 3M solution of ethyl magnesium bromide in diethyl ether (2.20 eq.). The resulting reaction was stirred at – 78 °C for 1 hour. The reaction was warmed to room temperature and stirred for a further hour. The reaction was cooled to 0 °C before the addition of boron trifluoride diethyl etherate (2.00 eq.). The reaction was warmed to room temperature and stirred for 1 hour. Once complete, the reaction was quenched by the addition of 1M HCl (3 mL/mmol) and diluted in diethyl ether (15 mL/mmol). 10% aqueous solution of sodium hydroxide was added (10 mL/mmol). The resulting suspension was filtered through celite. The two clear phases of the filtrate were separated and the organic layer was dried ($MgSO_4$) and concentrated *in vacuo* to yield crude product. Desired product was isolated by flash chromatography on silica gel using hexane/ethyl acetate (1:1) as eluent.

General procedures for reductive amination (General Procedures **4**)

General Procedure **4A**:

To a solution of amine (200 mg, 1.00 eq.) in THF (10 mL), aldehyde (1.30 eq.) was added. The reaction was stirred at room temperature for 24 hours. Sodium borohydride (1.30 eq.) was added and the reaction was stirred for 30 minutes. The reaction was quenched with water and extracted with ethyl acetate (3 x 15 mL). The combined organic extracts were washed with brine (3 x 15 mL), dried ($MgSO_4$)

and concentrated *in vacuo* to yield crude product. Desired product was isolated by flash chromatography on silica gel using hexane/ethyl acetate (1:1) as eluent.

General Procedure **4B**:

To a solution of amine (1.00 eq.) in THF (10 mL), aldehdyde (1.30 eq.) was added. The reaction was stirred at room temperature for 24 hours. The reaction was concentrated *in vacuo* and the residue was dissolved in methanol (10 mL). Sodium borohydride (1.30 eq.) was added and the reaction was stirred for 30 minutes. The reaction was quenched with water and extracted with ethyl acetate (3 x 10 mL). The combined organic extracts were washed with brine (3 x 15 mL), dried ($MgSO_4$) and concentrated *in vacuo.* Desired product was isolated by flash chromatography on silica gel using hexane/ethyl acetate (1:1) as eluent.

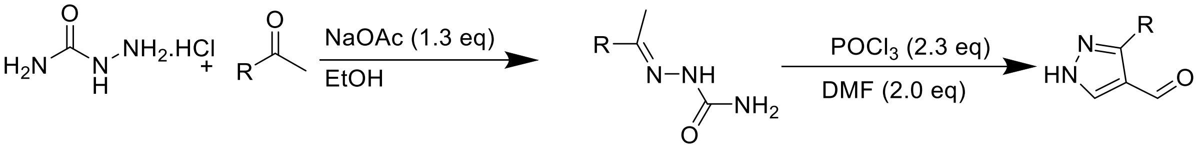


**Scheme S1**. Synthetic procedure used for the preparation of semicarbazones and aldehydes.

**<u>Analysis of Compounds</u>: see attached Appendix for NMR spectra**

**Analysis of Semicarbazones:**

2-(1-(thiophen-2-yl)ethylidene)hydrazine-1-carboxamide (**S1**) [(2)]

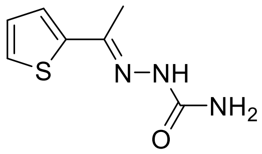

Compound synthesised using General Procedure 1 with 2-acetylthiophene (5.09 g, 40.4 mmol) to yield **S1** (5.17, 28.2 mmol, 63%) as a white crystalline solid.

**M.p** 190-191 °C  Lit: 187-190 °C [(3)]

**$^1$H NMR** (400 MHz, DMSO) δ 9.41 (s, 1H), 7.50 (dd, J = 5.1, 1.1 Hz, 1H), 7.37 (dd, J = 3.7, 1.2 Hz, 1H), 7.05 (dd, J = 5.1, 3.7 Hz), 6.29 (s, 2H), 2.21 (s, 3H)

Analytical data were in accordance with previous reports.[(2,3)]

2-(1-phenylethylidene)hydrazine-1-carboxamide (**S2**) [(2)]

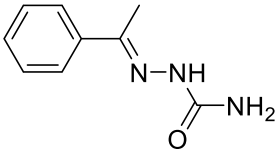

Compound synthesised using General Procedure 1 with acetophenone (4.85 g, 40.4 mmol) to yield **S2** (6.51 g, 36.8 mmol, 82%) as a white crystalline solid.

**M.p** 220-223 °C Lit: 224 °C [4]

**$^1$H NMR** (400 MHz, DMSO) δ 9.34 (s, 1H), 7.86-7.81 (m, 2H), 7.40-7.32 (m, 3H), 6.50 (s, 2H), 2.18 (s, 3H)

Analytical data were in accordance with previous reports. [2,4]

**Analysis of Aldehydes:**

3-(thiophen-2-yl)-1H-pyrazole-4-carbaldehyde (**S3**) [2]

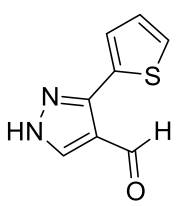

Compound synthesised using General Procedure 2 with semicarbazone **S1** (5.17 g, 28.2 mmol) to yield **S3** (1.60 g, 8.97 mmol, 32%) as a yellow solid.

**M.p** 77-79 °C Lit: 69-71 °C [2]

**$^1$H NMR** (400 MHz, $CDCl_3$) δ 10.11 (s, 1H), 8.19 (s, 1H), 7.80 (dd, J = 3.7, 1.2 Hz, 1H), 7.48 (dd, J = 5.1, 1.1 Hz, 1H), 7.19 (dd, J = 5.1, 3.7 Hz, 1H)

Analytical data were in accordance with previous reports. [2]

4-phenyl-1H-pyrazole-3-carbaldehyde (**S4**) [2]

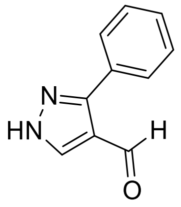

Compound synthesised using General Procedure 2 with semicarbazone **S2** (6.51 g, 36.8 mmol) to yield **S4** (3.54 g, 20.6 mmol, 56%) as a white solid.

**M.p :** 139-142 °C Lit: 145 °C [2]

**$^1$H NMR** (400 MHz, $CDCl_3$) δ 10.00 (s, 1H), 8.16 (s, 1H), 7.72-7.66 (m, 2H), 7.57-7.51 (m, 3H)

Analytical data were in accordance with previous reports. [2]

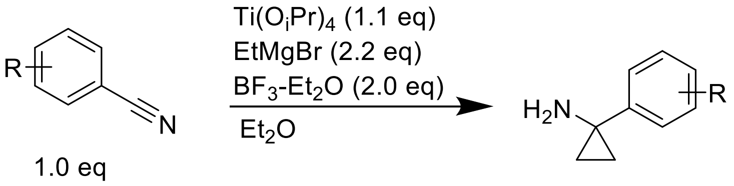


**Scheme S2**. Summary of procedure for the synthesis of the required amines. (1)

**Analysis of Amines:**

1-(4-methoxyphenyl)cyclopropan-1-amine (**S5**) (1)

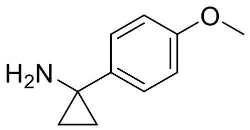


Compound synthesised using General Pocedure 3 with 4-methoxybenzonitrile (3.23 g, 24.2 mmol) to yield **S5** (2.21g, 13.6 mmol, 56%) as a yellow oil.

**$^1$H NMR** (400 MHz, $CDCl_3$) δ 7.29-7.24 (m, 2H), 6.90-6.85 (m, 2H), 3.82 (s, 3H), 1.90 (s, 2H), 1.05-1.01 (m, 2H), 0.95-0.92 (m, 2H)

Analytical data were in accordance with previous reports. (1)

1-phenylcyclopropan-1-amine (**S6**) (1)

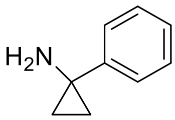


Compound synthesised using General Procedure 3 with benzonitrile (2.50 g, 24.3 mmol) to yield **S6** (1.36 g, 10.2 mmol ,42%) as a yellow oil.

**$^1$H NMR** (400 MHz, $CDCl_3$) δ 7.36-7.32 (m, 4H), 7.24-7.21 (m, 1H),1 91 (s, 2H), 1.11-1.09 (m, 2H), 1.02-1.00 (m, 2H)

Analytical data were in accordance with previous reports. (1)

1-(p-tolyl)cyclopropan-1-amine (**S7**) (1)

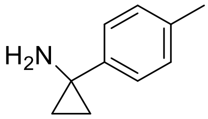


Compound synthesised using General Procedure 3 with p-tolylnitrile (1.00 g, 8.50 mmol) to yield **S7** (125 mg, 0.85 mmol, 10%) as a yellow oil.

**$^1$H NMR** (400 MHz, $CDCl_3$) δ 7.24-7.21 (m, 2H), 7.17-7.14 (m, 2H), 2.36 (s, 3H), 1.92 (s, 2H), 1.08-1.05 (m, 2H), 0.99-0.95 (m, 2H).

Analytical data were in accordance with previous reports. [1]

1-(4-chlorophenyl)cyclopropan-1-amine (**S8**) [1]

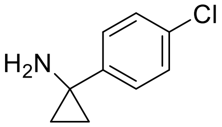


Compound synthesised using General Procedure 3 with 4-chlorobenzonitrile (1.00g, 7.27 mmol) to yield **S8** (451 mg, 2.69 mmol, 37%) as a yellow oil.

**$^1$H NMR** (400 MHz, $CDCl_3$) δ 7.30-7.27 (m, 2H), 7.26-7.23 (m, 2H), 1.89 (s, 2H), 1.11-1.08 (m, 2H), 0.99-0.96 (m, 2H).

Analytical data were in accordance with previous reports. [1]

1-(3,4-dichlorophenyl)cyclopropan-1-amine (**S9**)

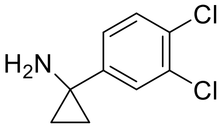


Compound synthesised using General Procedure 3 with 3,4,-dichlorobenzonitrile (3.00g, 17.4 mmol) to yield **S9** (1.97 g, 9.77 mmol, 56%) as a yellow oil.

**$^1$H NMR** (400 MHz, $CDCl_3$) δ 7.42 (d, J = 2.2 Hz, 1H), 7.37 (dd, J = 8.4, 1.2 Hz, 1H), 7.10 (dd, J = 8.4. 2.2 Hz, 1H), 1.88 (s, 2H), 1.14-1.10 (m, 2H), 1.00-0.97 (m, 2H)

**$^{13}$C NMR** (101 MHz, $CDCl_3$) δ 147.5 (ArC), 132.3 (ArC), 130.2 (ArCH), 129.6 (ArC), 127.5 (ArCH), 124.7 (ArCH), 36.1 (C), 18.6 ($CH_2$)

**HRMS** (ESI) m/z calculated for $C_9H_{10}Cl_2N$ $[M+H]^+$ : 202.0190; found: 202.0183

**Analysis of KNX-002 and its analogs:**

1-(4-methoxyphenyl)-N-((3-thiophen-2-yl)-1H-pyrazol-4-yl)methyl)cyclopropane-1-amine (**KNX-002**)

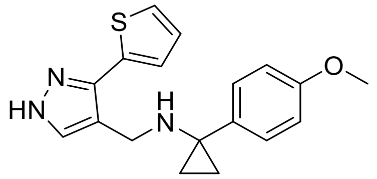

Compound synthesised using General Procedure 4a with **S5** (100 mg, 0.61 mmol), **S3** (141 mg, 0.79 mmol) and sodium borohydride (30 mg, 0.79 mmol) to yield **KNX-002** (15.0 mg, 0.06 mmol, 10%) as a white solid.

**MP.** 123-125 °C

**$^1$H NMR** (400 MHz, $CDCl_3$) δ 7.49 (s, 1H, ArH), 7.35-7.31 (m, 2H, ArH), 7.28-7.26 (m, 1H, ArH), 7.09 (dd, J = 3.6, 1.1, 1H, ArH), 7.01 (dd, J = 5.0, 3.5 Hz, 1H, ArH), 6.92-6.89 (m, 2H, ArH), 3.85 (s, 3H, $CH_3$), 3.76 (s, 2H, $CH_2$), 1.06-1.04 (m, 2H, $CH_2$), 0.92-0.89 (m, 2H, $CH_2$).

**$^{13}$C NMR** (101 MHz, $CDCl_3$) δ 158.6 (ArC), 142.7 (ArC), 134.9 (ArC), 134.5 (ArC), 132.0 (ArCH), 129.7 (ArCH), 127.5 (ArCH), 125.1 (ArCH), 125.0 (ArCH), 117.1 (ArC), 113.7 (ArCH), 55.3 ($CH_3$), 42.3 (C), 40.2 ($CH_2$), 14.7 ($CH_2$).

**IR** (FTIR) 3101, 2922, 1610, 1514, 1247, 1029, 810, 709, 555 $cm^{-1}$

**HRMS** (ESI) m/z calculated for $C_{18}H_{20}N_3OS$ $[M+H]^+$ : 326.1327; found: 326.1314.

N-((3-(thiophen-2-yl)-1H-pyrazol-4-yl)methyl)cyclopropanamine (**VEST1**)

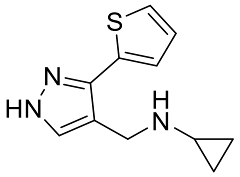

Compound was synthesised using General Procedure 4a with cyclopropylamine (100 mg, 1.75 mmol), **S3** (405 mg, 2.28 mmol) and sodium borohydride (86.0 mg, 2.28 mmol) to yield **VEST1** (78.0 mg, 0.36 mmol, 20%) as a colourless oil.

**$^1$H NMR** (400 MHz, $CDCl_3$) δ 7.59 (s, 1H, ArCH), 7.39 (dd, J = 3.6, 1.1 Hz, 1H, ArCH), 7.35 (dd, J = 5.1, 1.1 Hz, 1H, ArCH), 7.13 (dd, J = 5.1, 3.6 Hz, 1H, ArCH), 3.96 (s, 2H, $CH_2$), 2.25-2.21 (m, 1H, CH), 0.50-0.45 (m, 2H, $CH_2$), 0.45-0.42 (m 2H, $CH_2$)

**$^{13}$C NMR** (101 MHz, $CDCl_3$) δ 142.3 (ArC), 134.2 (ArC), 132.8 (ArCH), 127.6 (ArCH), 125.2 (ArCH), 125.1 (ArCH), 117.2 (ArC), 43.41 ($CH_2$), 30.3 (CH), 6.3 ($CH_2$)

**IR** (FTIR) 3113, 2897, 1068, 925, 813, 694, 617, 445 $cm^{-1}$

**HRMS (**ESI) m/z calculated for $C_{11}H_{14}N_3S$ $[M+H]^+$: 220.0908; found 220.0900

1-(4-chlorophenyl)-N-((3-(thiophen-2-yl)-1H-pyrazol-4-yl)methyl)cyclopropan-1-amine (**VEST2**)

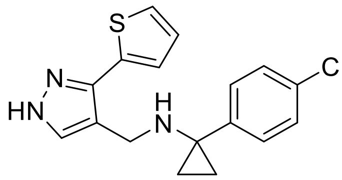


Compound synthesised using General Procedure 4b with **S8** (200 mg, 1.19 mmol), **S3** (276 mg ,1.55 mmol) and sodium borohydride (59 mg, 1.55 mmol) to yield **VEST2** (34mg, 0.10 mmol, 8%) as an off white solid.

**M.p** 126-129 °C

**$^{1}$H NMR** (400 MHz, $CDCl_3$) δ 7.51 (s, 1H, ArCH), 7.34-7.30 (m, 5H, ArCH), 7.14 (dd, J = 3.6, 1.1 Hz, 1H, ArCH), 7.05 (dd, J = 5.1, 3.6 Hz, 1H, ArCH), 3.75 (s, 2H, $CH_2$), 1.10-1.07 (m, 2H, $CH_2$), 0.95-0.93 (m, 2H, $CH_2$)

**$^{13}$C NMR** (101 MHz, $CDCl_3$) δ 142.3 (ArC), 141.6 (ArC), 134.8 (ArC), 132.2 (ArC), 132.1 (ArCH), 129.4 (ArCH), 128.5 (ArCH), 127.5 (ArCH) 127.4 (ArC), 125.2 (ArCH), 125.1 (ArCH), 42.1 (C), 40.3 ($CH_2$), 15.4 ($CH_2$)

**IR** (FTIR) 3265, 2850, 1259, 1099, 1010, 813, 702, 528 $cm^{-1}$

**HRMS (**ESI) m/z calculated for $C_{17}H_{17}ClN_3S$ $[M+H]^+$: 330.0832; found 330.0817

N-((3-(thiophen-2-yl)-1H-pyrazol-4-yl)methyl)-1-(p-tolyl)cyclopropan-1-amine (**VEST3**)

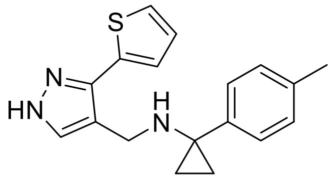


Compound synthesised using General Procedure 4b using **S7** (200 mg, 1.36 mmol), **S3** (315 mg ,1.77 mmol) and sodium borohydride (68 mg ,1.77 mmol) 1 to yield **VEST3** (203 mg, 0.66 mmol, 49%) as a colourless oil.

**$^{1}$H NMR** (400 MHz, $CDCl_3$) δ 7.51 (s, 1H, ArCH), 7.30-7.27 (m, 3H, ArCH), 7.19 (d, J = 7.7 Hz, 2H, ArCH), 7.13 (dd, J = 3.6, 1.1 Hz, 1H, ArCH), 7.02 (dd, J = 5.1, 3.5 Hz, 1H, ArCH), 3.77 (s, 2H, $CH_2$), 2.39 (s, 3H, $CH_3$), 1.07-1.05 (m, 2H, $CH_2$), 0.94-0.92 (m ,2H, $CH_2$)

**$^{13}$C NMR** (101 MHz, $CDCl_3$) δ 142.6 (ArCH), 139.9 (ArC), 136.3 (ArC), 134.2 (ArC) 132,4 (ArCH), 129.0 (ArCH), 128.1 (ArCH), 127.5 (ArCH), 125.1 (ArCH), 125.0 (ArCH), 117.1 (ArC), 42.4 (C), 40.3 ($CH_2$), 21.1 ($CH_3$), 15.1 ($CH_2$)

**IR** (FTIR) 3269, 3084, 2916, 2661, 1257, 1082, 923, 823,698, 555 $cm^{-1}$

**HRMS (**ESI) m/z calculated for $C_{18}H_{20}N_3S$ $[M+H]^+$: 310.1378; found 310.1364

1-(3,4-dichlorophenyl)-N-((3-(thiophen-2-yl)-1H-pyrazol-4-yl)methyl)cyclopropan-1-amine (**VEST4**)

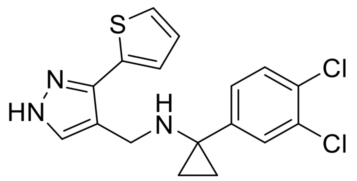

Compound synthesised using General Procedure 4b with **S9** (200 mg, 0.99 mmol), **S3** (230 mg, 1.29 mmol) and sodium borohydride (49 mg, 1.29 mmol) to yield **VEST4** (47 mg, 0.13 mmol, 13%) as a white solid.

**M.p** 54-57 °C

**$^1$H NMR** (400 MHz, $CDCl_3$) δ 7.49 (s, 1H, ArCH), 7.47 (d, J = 2.1 Hz, 1H, ArCH), 7.41 (d, J = 8.3 Hz, 1H, ArCH), 7.31 (dd, J = 5.1, 1.1 Hz, 1H, ArCH), 7.21-7.17 (m, 2H, ArCH), 7.06 (dd, J = 5.1, 3.6 Hz, 1H, ArCH), 3.74 (s, 2H, $CH_2$), 1.13-1.09 (m, 2H, $CH_2$), 0.97-0.93 (m, 2H, $CH_2$)

**$^{13}$C NMR** (101 MHz, $CDCl_3$) δ 143.6 (ArC), 142.6 (ArC), 134.0 (ArC), 132.4 (ArC), 132.0 (ArCH), 130.28 (ArC), 130.3 (ArCH), 129.9 (ArCH), 127.6 (ArCH), 127.1 (ArCH), 125.3 (ArCH), 125.2 (ArCH), 116.7 (ArC), 42.0 (C), 40.3 ($CH_2$), 15.6 ($CH_2$)

**IR** (FTIR) 3155, 2942, 1471, 1375, 1261, 1082, 1026, 798, 694,677, 609 $cm^{-1}$

**HRMS (**ESI) m/z calculated for $C_{17}H_{16}Cl_2N_3S$ $[M+H]^+$: 364.0442; found 364.0427

1-phenyl-N-((3-(thiophen-2-yl)-1H-pyrazol-4-yl)methyl)cyclopropan-1-amine (**VEST5**)

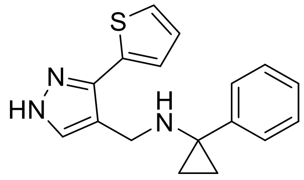

Compound synthesised using General Procedure 4b with **S6** (200 mg, 1.50 mmol), **S3** (347 mg, 1.95 mmol) and sodium borohydride (74 mg, 1.95 mmol) to yield **VEST5** (73 mg, 0.25 mmol, 17%) as an off white solid.

**M.p** 105-108 °C

**$^1$H NMR** (400 MHz, $CDCl_3$) δ 7.52 (s, 1H, ArCH), 7.41-7.36 (m, 4H, ArCH), 7.30-7.27 (m, 2H, ArCH), 7.13 (dd, J = 3.6, 1.1 Hz, 1H, ArCH), 7.03 (dd, J = 5.1, 3.5 Hz, 1H, ArCH), 3.79 (s, 2H, $CH_2$), 1.10-1.08 (m, 2H, $CH_2$), 0.98-0.96 (m ,2H, $CH_2$)

**$^{13}$C NMR** (101 MHz, $CDCl_3$) δ 143.0 (ArC), 142.2 (ArC), 134.7 (ArC), 131.6 (ArCH), 128.4 (ArCH), 128.0 (ArCH), 127.5 (ArCH), 126.6 (ArCH), 125.1 (ArCH), 125.0 (ArCH), 117.0 (ArC), 42.7 (C), 40.4 ($CH_2$), 15.2 ($CH_2$)

**IR** (FTIR) 3383, 3163, 2943, 1076, 1051, 813, 690 $cm^{-1}$

**HRMS (**ESI) m/z calculated for $C_{17}H_{18}N_3S$ $[M+H]^+$: 296.1221; found 296.1212

1-(4-methoxyphenyl)-N-((3-phenyl-1H-pyrazol-4-yl)methyl)cyclopropan-1-amine (**VEST6**)

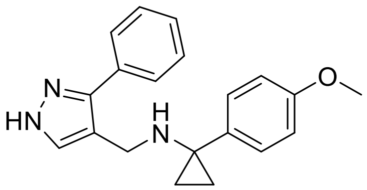

Compound synthesised using General Procedure 4b with **S5** (200 mg, 1.22 mmol), **S4** (274 mg, 1.59 mmol) to yield **VEST6** (82.0 mg, 0.26 mmol, 21%) as a white solid.

**M.p** 115-119 °C

**$^1$H NMR** (400 MHz, $CDCl_3$) δ 7.54 (dd, J = 7.7, 1.9 Hz, 2H, ArCH), 7.52 (s, 1H, ArCH), 7.40-7.36 (m, 3H, ArCH), 7.29 (d, J = 8.2, 2H, ArCH), 6.90-6.87 (m, 2H, ArCH), 3.84 (s, 3H, $CH_3$), 3.70 (s, 2H, $CH_2$), 1.02-1.00 (m ,2H, $CH_2$), 0.90-0.88 (m, 2H, $CH_2$)

**$^{13}$C NMR** (101 MHz, $CDCl_3$) δ 158.3 (ArC), 145.4 (ArC), 135.1 (ArC), 132.0 (ArC) 129.4 (ArCH), 128.7 (ArCH), 128.0 (ArCH), 127.4 (ArCH), 117.1 (ArC), 113.6 (ArCH), 55.3 ($CH_3$), 42.2 (C), 40.1 ($CH_2$), 14.8 ($CH_2$)

**IR** (FTIR) 2889, 2353, 1610, 1514, 1417, 1249, 1033, 831, 804, 698, 557 $cm^{-1}$

**HRMS** (ESI) m/z calculated for $C_{20}H_{22}N_3O$ $[M+H]^+$: 320.1763; found 320.1751

N-benzyl-1-(4-methoxyphenyl)cyclopropan-1-amine (**VEST7**)

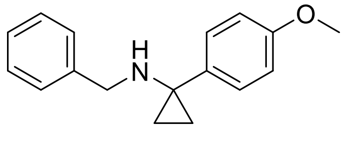

Compound synthesised using General Procedure 4b using **S5** (200 mg, 1.22 mmol), benzaldehyde (169 mg, 1.59 mmol) and sodium borohydride (60.0 mg, 1.59 mmol) to yield **VEST7** (120 mg, 0.51 mmol, 42%) as a white solid.

**MP.** 43-45 °C

**$^1$H NMR** (400 MHz, $CDCl_3$) δ 7.35-7.32 (m, 2H, ArCH), 7.31-7.26 (m, 4H, ArCH), 7.25-7.22 (m, 1H, ArCH), 6.93-6.90 (m, 2H, ArCH), 3.85 (s, 3H, $CH_3$), 3.69 (s, 2H, $CH_2$), 1.03-1.00 (m, 2H, $CH_2$), 0.92-0.90 (m, 2H, $CH_2$)

**$^{13}$C NMR** (101 MHz, $CDCl_3$) δ 158.2 (ArC), 140.8 (ArC), 135.5 (ArC), 129.1 (ArCH), 128.3 (ArCH), 128.2 (ArCH), 126.8 (ArCH), 113.7 (ArCH) 55.3 ($CH_3$), 50.6 ($CH_2$), 42.2 (C), 15.3 ($CH_2$)

**IR** (FTIR) 3008, 2358, 1606, 1504, 1234, 1176, 1026, 837, 740, 655, 559 $cm^{-1}$

**HRMS** (ESI) m/z calculated for $C_{17}H_{20}NO$ $[M+H]^+$: 254.1545; found 245.1535

1-(4-methoxyphenyl)-N-(thiazol-5-ylmethyl)cyclopropan-1-amine (**VEST8**)

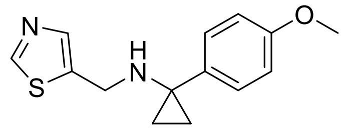

Compound synthesised using General Procedure 4b using **S5** (200 mg, 1.22 mmol), thiazole-4-carbaldehyde (180 mg, 1.59 mmol) and sodium borohydride (60.0 mg, 1.59 mmol) to yield **VEST8** (17 mg, 0.06 mmol, 5%) as a tan solid.

**M.p** 60-63 **°C**

**$^{1}$H NMR** (400 MHz, $CDCl_3$) δ 8.71 (s, 1H, ArH) 7.65 (s, 1H, ArH), 7.33-7.29 (m, 2H, ArH), 6.93-6.89 (m, 2H, ArH), 3.92 (s, 2H, $CH_2$), 3.84 (s, 3H, $CH_3$), 1.01-0.99 (m, 2H, $CH_2$), 0.91-0.89 (m, 2H, $CH_2$)

**$^{13}$C NMR** (101 MHz, $CDCl_3$) δ 158.6 (ArC), 152.6 (ArCH), 140.4 (ArCH), 139.2 (ArC), 134.7 (ArC), 129.2 (ArCH), 113.8 (ArCH), 55.3 ($CH_3$), 42.7 ($CH_2$), 42.3 (C), 15.0 ($CH_2$)

**IR** (FTIR) 3255, 2831, 1608, 1514, 1438, 1236, 1026, 8031, 613, 543 $cm^{-1}$

**HRMS** (ESI) m/z calculated for $C_{14}H_{17}N_2OS$ $[M+H]^+$: 261.1062; found 261.1056

N-(furan-3-ylmethyl)-1-(4-methoxyphenyl)cyclopropan-1-amine (**VEST9**)

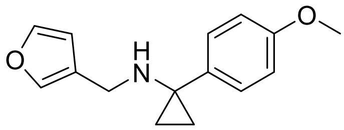

Compound synthesised using General Procedure 4b using **S5** (200 mg, 1.22 mmol), furan-3-carbaldehyde (153 mg, 1.59 mmol) and sodium borohydride (60 mg, 1.59 mmol) to yield **VEST9** (46 mg, 0.19 mmol, 16%) as a colourless oil.

**$^{1}$H NMR** (400 MHz, $CDCl_3$) δ 7.36 (dd, J = 1.7 Hz, 1H, ArCH), 7.33-7.29 (m, 3H, ArCH), 6.92-6.89 (m, 2H, ArCH), 6.33-6.31 (m ,1H, ArCH), 3.84 (s, 3H, $CH_3$), 3.55 (s, 2H, $CH_2$), 1.02-1.00 (m, 2H, $CH_2$), 0.90-0.88 (m, 2H, $CH_2$)

**$^{13}$C NMR** (101 MHz, $CDCl_3$) δ 158.2 (ArC), 142.9 (ArCH), 139.6 (ArCH), 135.1 (ArC), 129.1 (ArCH), 124.4 (ArC), 113.7 (ArCH), 110.6 (ArCH), 55.3 ($CH_3$), 42.1 (C), 41.1 ($CH_2$), 15.1 ($CH_2$)

**IR** (FTIR) 3298, 3130, 3003, 2846, 1604, 1510, 1234, 1016, 871, 763, 543 $cm^{-1}$

**HRMS** (ESI) m/z calculated for $C_{15}H_{18}NO_2$ $[M+H]^+$: 244.1338; found 244.1332

N-((1H-pyrrol-3-yl)methyl)-1-(4-methoxyphenyl)cyclopropan-1-amine (**VEST10**)

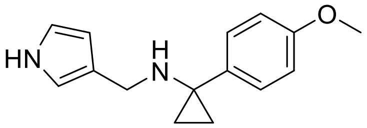

Compound synthesised using General Procedure 4b using **S5** (200 mg, 1.22 mmol), 1*H*-pyrrole-3-carbaldehyde (151 mg, 1.59 mmol) and sodium borohydride (60 mg, 1.59 mmol) to yield **VEST10** (26 mg, 0.11 mmol, 9%) as a tan solid.

**M.p** 93-96 °C

**$^{1}$H NMR** (400 MHz, $CDCl_3$) δ 7.34-7.31 (m, 2H, ArCH), 6.92-6.88 (m, 2H, ArCH), 6.73-6.71 (m, 1H, ArCH), 6.68-6.66 (m, 1H, ArCH), 6.14-6.12 (m, 1H, ArCH), 3.84 (s, 3H, $CH_3$), 3.61 (s, 2H, $CH_2$), 1.04-1.02 (m, 2H, $CH_2$), 0.90-0.88 (m, 2H, $CH_2$)

**$^{13}$C NMR** (101 MHz, $CDCl_3$) δ 158.4 (ArC), 135.7 (ArC), 129.0 (ArCH), 123.1 (ArC), 117.8 (ArCH), 115.5 (ArCH), 113.6 (ArCH), 108.4 (ArCH), 55.4 ($CH_3$), 43.1 ($CH_2$), 42.2 (C), 15.2 ($CH_2$)

**IR** (FTIR) 3151, 2885, 1608, 1514, 1417, 1236, 1070, 1022, 833, 771, 707, 603, 561 $cm^{-1}$

**HRMS** (ESI) m/z calculated for $C_{15}H_{19}N_2O$ $[M+H]^+$: 243.1498; found 243.1490

N-(4-methoxybenzyl)-1-(3-(thiophen-2-yl)-1H-pyrazol-4-yl)methanamine (**VEST11**)

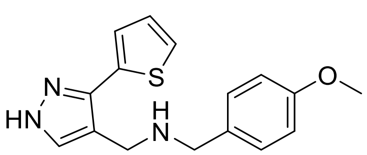


Compound was synthesised using General Procedure 4a with 4-methoxybenzylamine (200 mg, 1.46 mmol), **3** (339 mg, 1.90 mmol) and sodium borohydride (72 mg, 1.90 mmol) to yield **VEST11** (28 mg, 0.09 mmol, 6%) as a white solid. For characterisation purposes the HCl salt was prepared.

**M.p** 136-139 °C

**$^{1}$H NMR** (400 MHz, MeOD-d4) δ 8.11 (s, 1H, ArH), 7.60 (dd, J = 5.2, 1.1 Hz, 1H, ArH), 7.44-7.40 (m, 2H, ArH), 7.26 (dd, J = 3.6, 1.1 Hz, 1H, ArH), 7.16 (dd, J = 5.1, 3.6 Hz, 1H, ArH), 7.02-6.99 (m, 2 H, ArH), 4.35 (s, 2H, $CH_2$), 4.22 (s, 2H, $CH_2$), 3.84 (s, 3H, $CH_3$)

**$^{13}$C NMR** (101 MHz, MeOD-d4) δ 160.9 (ArC), 142.4 (ArC), 134.7 (ArCH), 131.5 (ArCH), 130.6 (ArC), 127.7 (ArC), 122.4 (ArC), 114.2 (ArCH), 108.7 (ArC), 54.5 ($CH_3$), 50.0 ($CH_2$), 39.7 ($CH_2$)

**IR** (FTIR) 3304, 2391, 2274, 1610, 1514, 1251, 1163, 1026, 935, 842, 715, 609, 534 $cm^{-1}$

**HRMS** (ESI) m/z calculated for $C_{16}H_{17}N_3OSNa$ $[M+Na]^+$: 322.0990; found 322.0979

N-benzyl-1-(3-(thiophen-2-yl)-1H-pyrazol-4-yl)methanamine (**VEST12**)

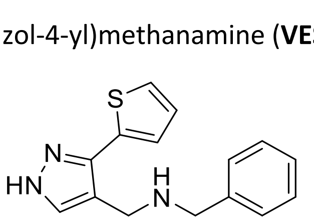


Compound was synthesised using General Procedure 4a using benzylamine (200 mg, 1.86 mmol), **S3** (431 mg, 2.42 mmol) and sodium borohydride (92.0 mg, 2.42 mmol) to yield **VEST12** (152 mg, 0.56 mmol, 30%) as a white solid. For characterisation purposes the HCl salt was prepared.

**M.p** 134-136 °C

**$^{1}$H NMR** (400 MHz, , MeOD-d4) δ 7.98 (s, 1H, ArCH), 7.55 (dd, J = 5.1, 1.1 Hz, 1H, ArCH), 7.49 (s, 5H, ArCH) 7.23 (dd, J = 3.6, 1.1 Hz, 1H, ArCH), 7.15 (dd, J = 5.1, 3.6 Hz, 1H, ArCH), 4.37 (s, 2H, ArCH), 4.27 (s, 2H, ArCH)

**$^{13}$C NMR** (101 MHz, MeOD-d4) δ 142.5 (ArC), 134.5 (ArCH), 130.8 (ArC), 129.8 (ArCH), 129.4 (ArCH), 129.0 (ArCH), 127.6 (ArCH), 126.5 (ArCH), 126.3 (ArCH), 108.2 (ArC), 50.4 ($CH_2$), 40.2 ($CH_2$)

**IR** (FTIR) 3143, 2931, 2812, 1452, 1041, 945, 839, 742, 700, 605 $cm^{-1}$

**HRMS** (ESI) m/z calculated for $C_{15}H_{16}N_3S$ $[M+H]^+$: 270.1065; found 270.1056

2-phenyl-N-((3-(thiophen-2-yl)-1H-pyrazol-4-yl)methyl)propan-2-amine (**VEST13**)

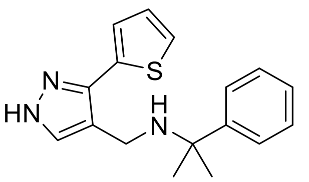


Compound was synthesised using General Procedure 4b using cumylamine (200 mg, 1.48 mmol), **S3** (342 mg, 1.92 mmol) and sodium borohydride (73.0 mg, 1.92 mmol) to yield **VEST13** (82.0 mg, 0.27 mmol, 18%) as an off white solid.

**M.p** 115-117 °C

**$^{1}$H NMR** (400 MHz, $CDCl_3$) δ 7.58 (s, 1H, ArCH), 7.52-7.50 (m ,2H, ArCH), 7.39-7.35 (m ,2H, ArCH), 7.31 (dd, J = 5.1, 1.1 Hz, 1H, ArCH), 7.28-7.25 (m ,1H, ArCH), 7.24 (dd, J = 3.6, 1.1 Hz, 1H, ArCH), 7.07 (dd, J = 5.1, 3.5 Hz, 1H, ArCH), 3.60 (s, 2H, $CH_2$), 1.57 (s, 6H, 2 x $CH_3$)

**$^{13}$C NMR** (101 MHz, $CDCl_3$) δ 147.5 (ArC), 142.6 (ArC), 134.9 (ArC), 131.3 (ArCH), 128.3 (ArCH), 127.5 (ArCH), 126.4 (ArCH), 125.8 (ArCH), 125.2 (ArCH), 125.0 (ArCH), 117.8 (ArC), 56.9 (C), 37.5 ($CH_2$), 29.4 ($CH_3$)

**IR** (FTIR) 3379, 3155, 2942, 1076, 846, 736, 690, 565 $cm^{-1}$

**HRMS (**ESI) m/z calculated for $C_{17}H_{20}N_3S$ $[M+H]^+$: 298.1378; found 298.1367

(*R*)-1-(4-methoxyphenyl)-N-((3-(thiophen-2-yl)-1H-pyrazol-4-yl)methyl)ethan-1-amine (**VEST14**)

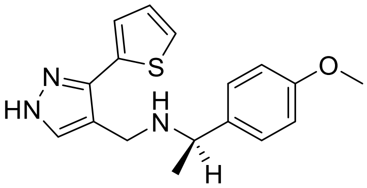


Compound synthesised using General Procedure 4a using (*R*)-1-(4-methoxyphenyl)-ethylamine (200 mg, 1.32 mmol), **S3** (340 mg, 1.72 mmol) and sodium borohydride (65.0 mg, 1.72 mmol) to yield **VEST14** (147 mg, 0.47 mmol, 62%) as an white solid.

**M.p** 131-134 °C

**$^{1}$H NMR** (400 MHz, $CDCl_3$) δ 7.54 (s, 1H, ArCH), 7.31 (dd, J = 5.1, 1.1 Hz, 1H, ArCH), 7.29-7.27 (m, 3H, ArCH), 7.08 (dd, J = 5.1, 3.6 Hz, 1H, ArCH), 6.91-6.89 (m, 2H, ArCH), 3.84-3.81 (m ,4H, $CH_{3,}$ CH), 3.72 (s, 2H, $CH_2$), 1.38 (d, J = 6.6 Hz, 3H, $CH_3$)

**$^{13}$C NMR** (101 MHz, $CDCl_3$) δ 158.6 (ArC), 142.8 (ArC), 137.3 (ArC), 134.6 (ArC), 132.4 (ArCH), 127.8 (ArCH), 127.6 (ArCH), 125.2 (ArCH), 125.1 (ArCH), 117.1 (ArC), 113.8 (ArCH), 57.3 ($CH_3$), 55.3 (CH), 41.6 ($CH_2$), 24.2 ($CH_3$)

**IR** (FTIR) 3161, 2954, 1510, 1247, 1035, 827, 756, 549 $cm^{-1}$

**HRMS** (ESI) m/z calculated for $C_{17}H_{20}N_3OS$ [M+H] 314.1327; found 314.1319

**[α]$_D$** -53.4 (c 0.01 in $CHCl_3$)

(*S*)-1-(4-methoxyphenyl)-N-((3-(thiophen-2-yl)-1H-pyrazol-4-yl)methyl)ethan-1-amine (**VEST15**)

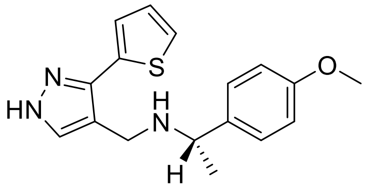

Compound synthesised using General Procedure 4a using (S)-1-(4-methoxyphenyl)-ethylamine (200 mg, 1.32 mmol), **S3** (340 mg, 1.72 mmol) and sodium borohydride (65 mg, 1.72 mmol) to yield **VEST15** (93mg, 0.30 mmol, 23%) as an off white solid.

**M.p** 131-132 °C

**$^{1}$H NMR** (400 MHz, $CDCl_3$) δ 7.54 (s, 1H, ArCH), 7.32 (dd, J = 5.1, 1.1 Hz, 1H, ArCH), 7.29-7.27 (m, 3H, ArCH), 7.08 (dd, J = 5.1, 3.6 Hz, 1H, ArCH), 6.92-6.89 (m, 2H, ArCH), 3.85-3.81 (m, 4H, $CH_{3,}$ CH), 3.72 (s, 2H, $CH_2$), 1.38 (d, J = 6.6 Hz, 3H, $CH_3$)

**$^{13}$C NMR** (101 MHz, $CDCl_3$) δ 158.6 (ArC), 141.2 (ArC), 137.3 (ArC), 134.6 (ArC), 132.4 (ArCH), 127.8 (ArCH), 127.6 (ArCH), 125.2 (ArCH), 125.1 (ArCH), 117.21 (ArC), 113.8 (ArCH) 57.3 ($CH_3$), 55.3 (CH), 41.6 ($CH_2$), 24.2 ($CH_3$)

**IR** (FTIR) 3149, 2953, 1510, 1247, 1033, 827, 756, 709, 549 $cm^{-1}$

**HRMS (**ESI) m/z calculated for $C_{17}H_{20}N_3OS$ $[M+H]^+$: 314.1327; found 314.1315

**[α]$_D$:** +53.7 (c 0.01 in $CHCl_3$)

References:


[1] P. Bertus, J. Szymoniak, *J. Org. Chem.,* 2003, **68**, 7133-7136

[2] A. V. Lebdev, A. B. Lebdeva, V. D, Sheludyakov, E. A. Kovaleva, O. L. Usinova, I. B. Kozhevnikov, *Russ. J. Gen. Chem.,* 2005, **75**, 782-789

[3] S. Shrivastava, N. Fahmi, R. V. Singh, *J. Sulphur Chem.,* 2010, **31**, 515-524

[4] L. Bhal, R. V. Singh, J. Tandon, *Acta Chim. Hung.,* 1984, **115**, 251-260

### Appendix

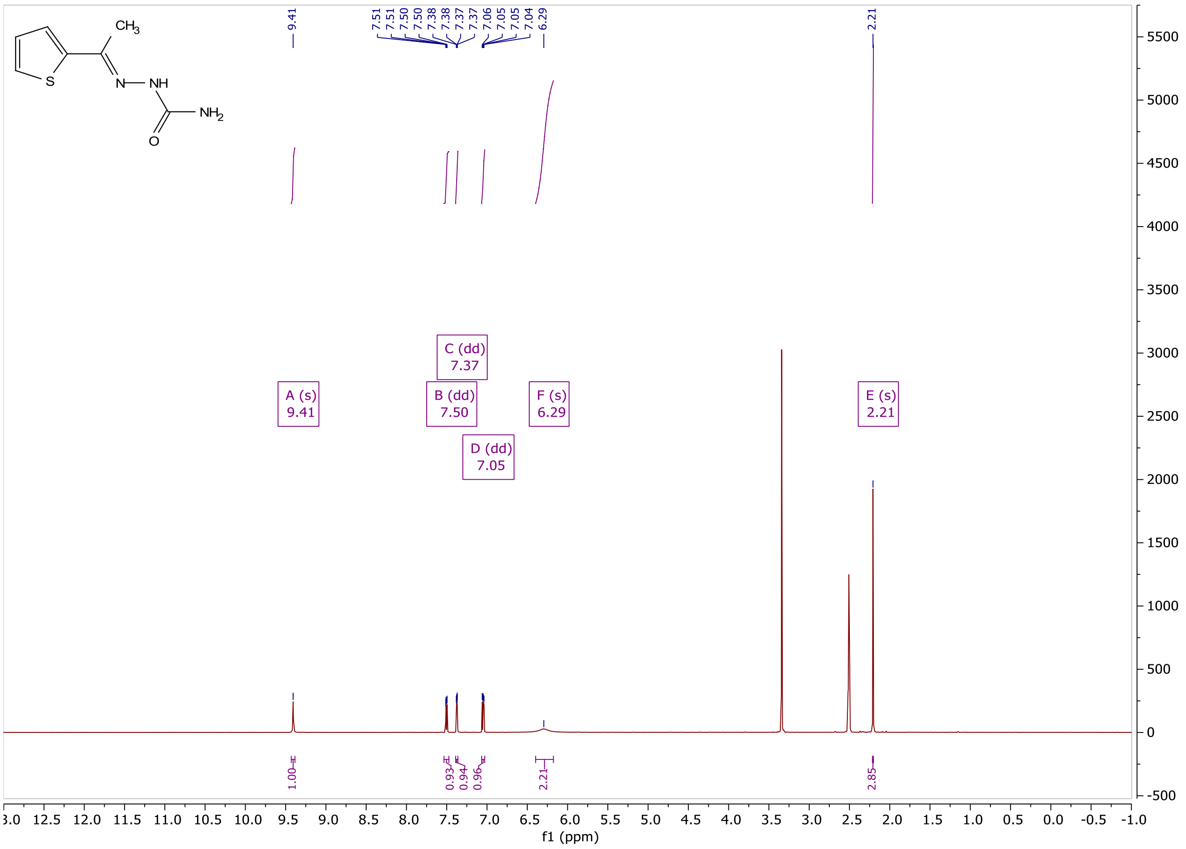


Figure **A1**: $^{1}$H NMR of **S1**

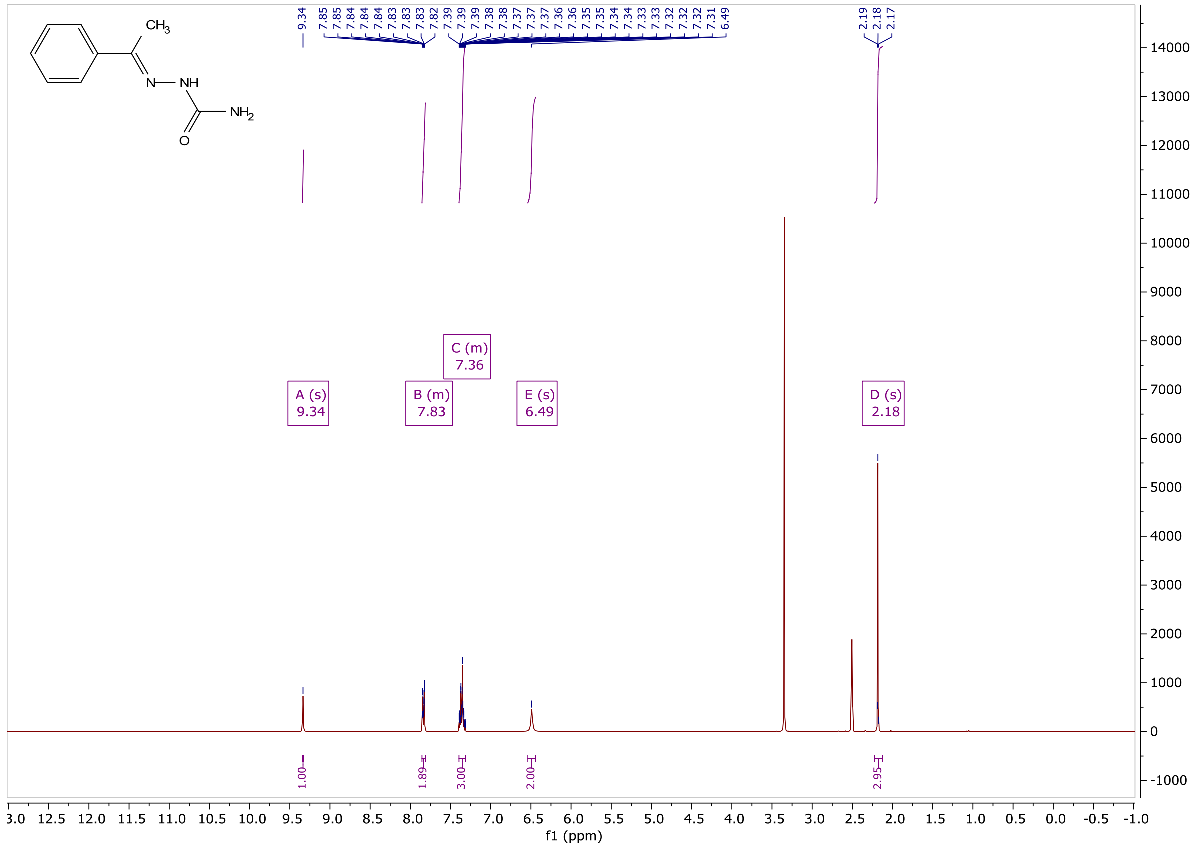

Figure **A2**: $^{1}$H NMR of **S2**

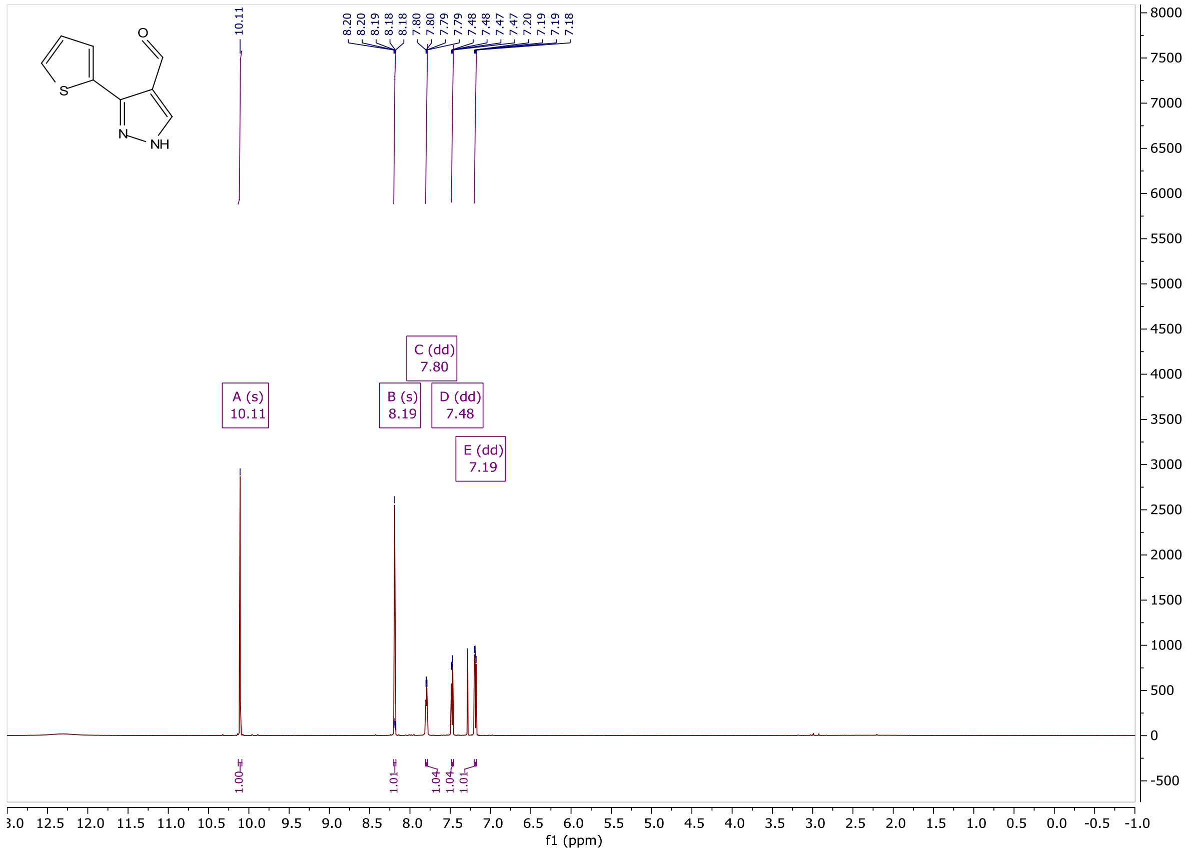

Figure **A3**: $^{1}$H NMR of **S3**

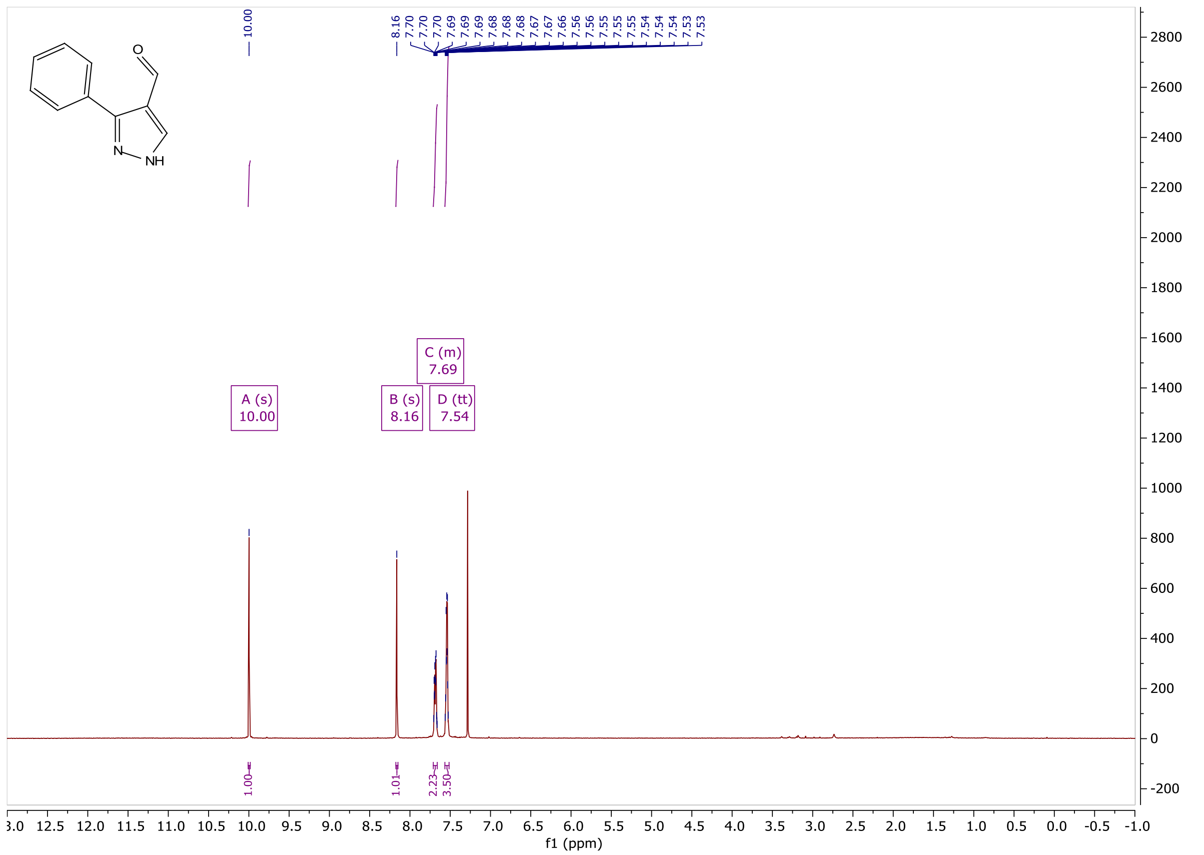


Figure **A4**: $^{1}$H NMR of **S4**

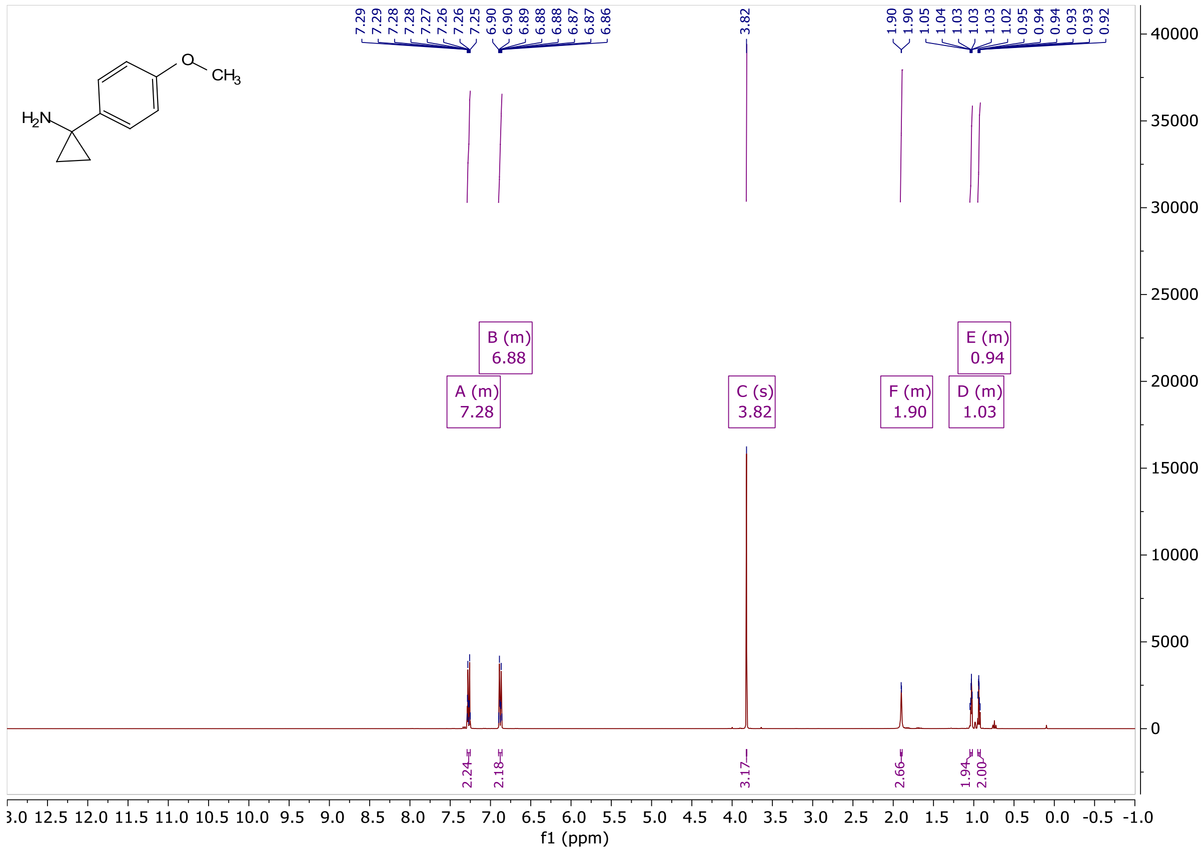


Figure **A5**: $^{1}$H NMR of **S5**

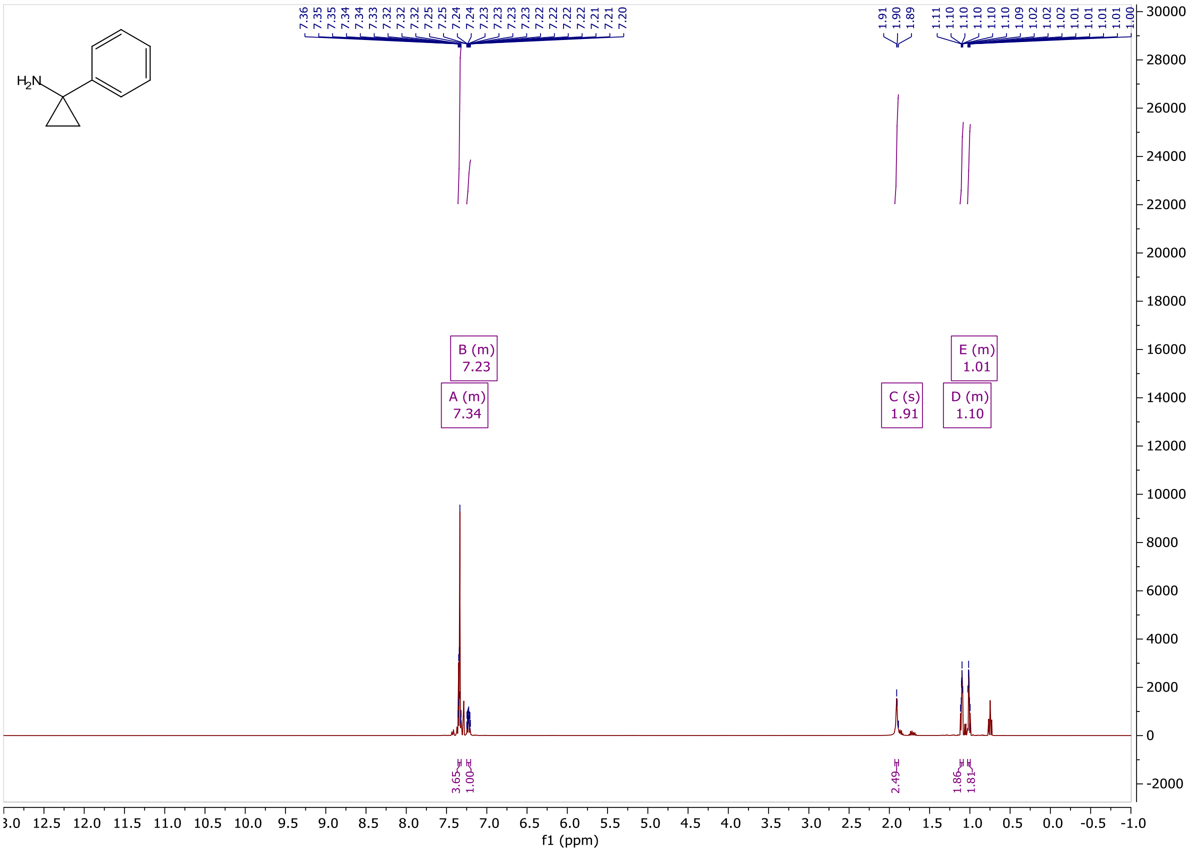

Figure **A6**: $^{1}$H NMR of **S6**

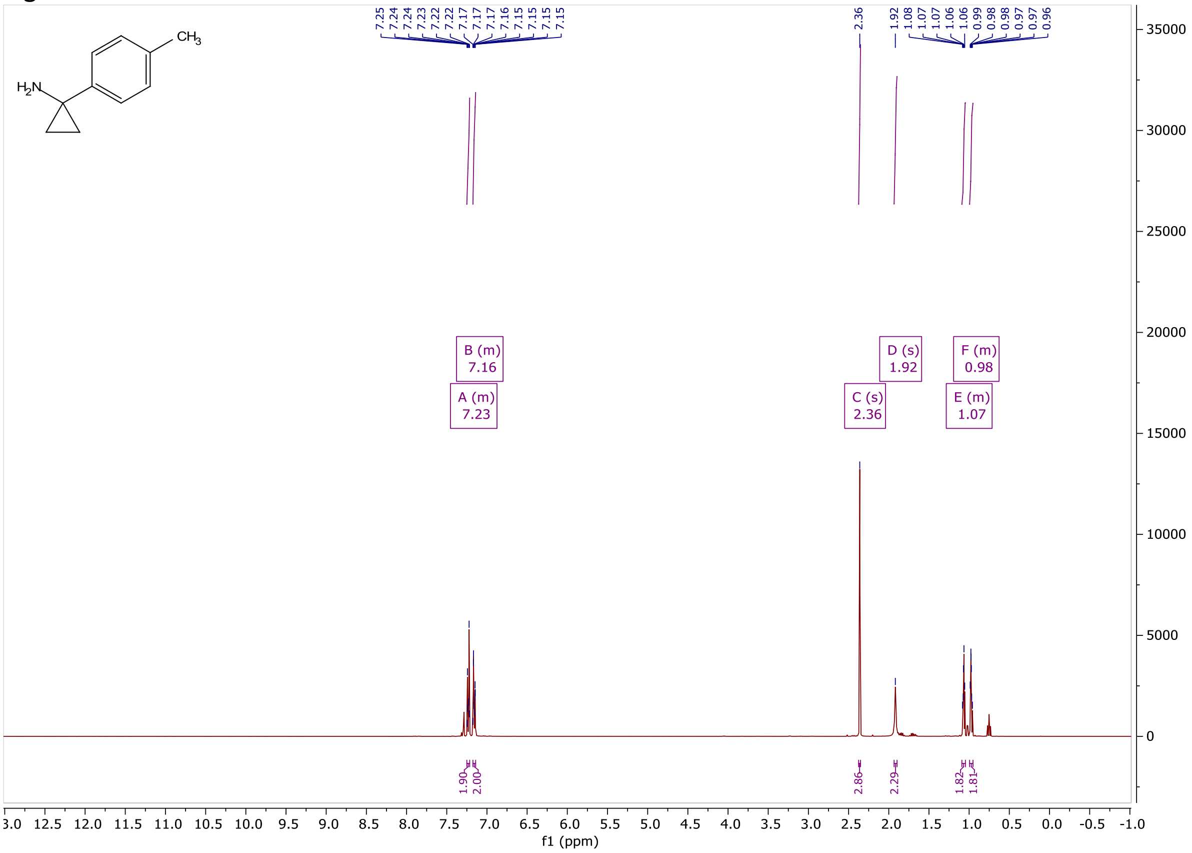

Figure **A7**: $^{1}$H NMR of **S7**

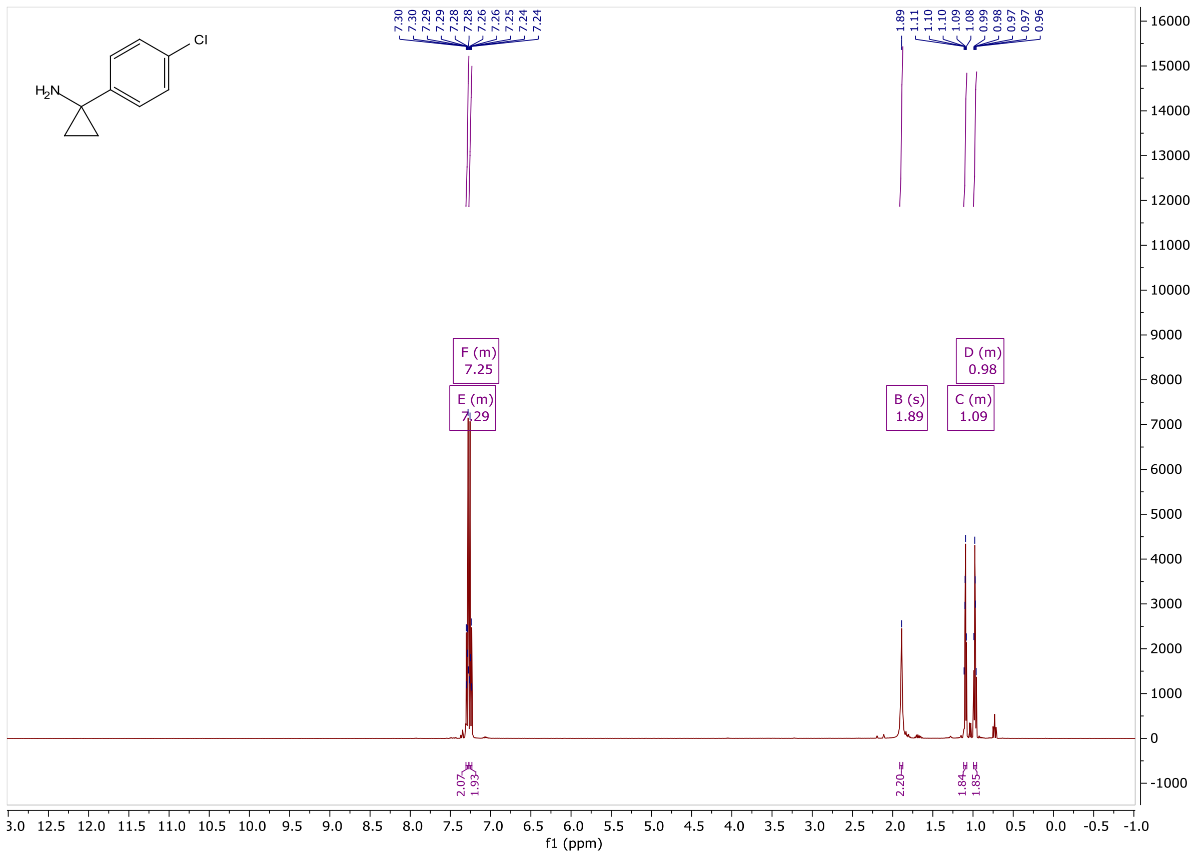

Figure **A8**: $^{1}$H NMR of **S8**

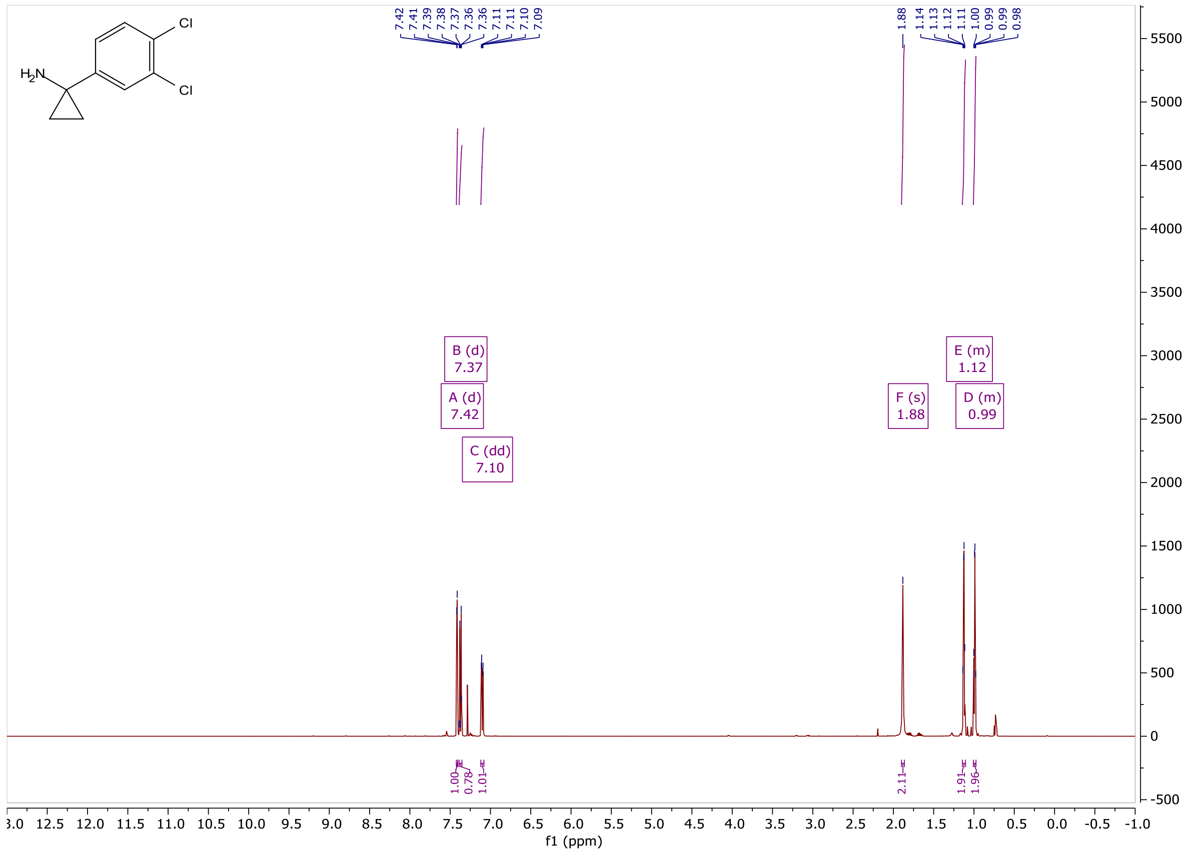

Figure **A9**: $^{1}$H NMR of **S9**

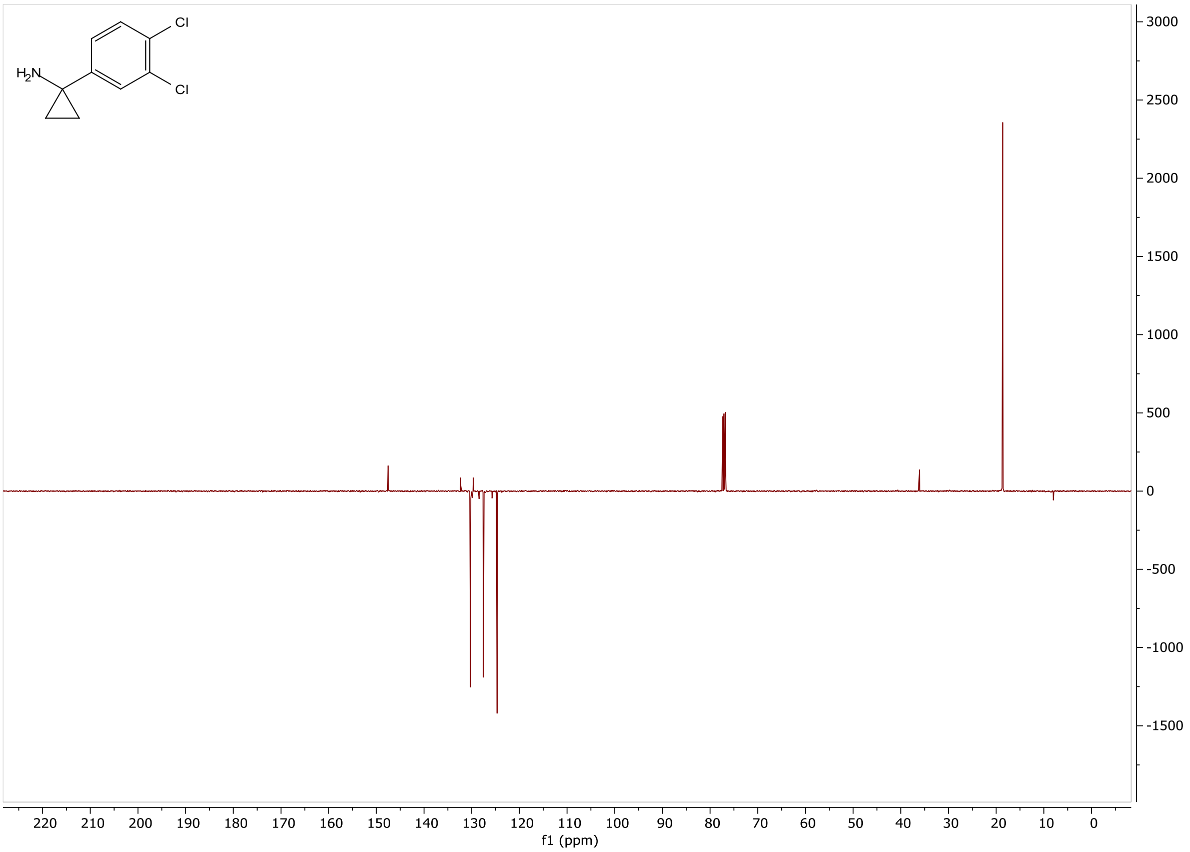

Figure **A10**: $^{13}$C NMR of **S9**

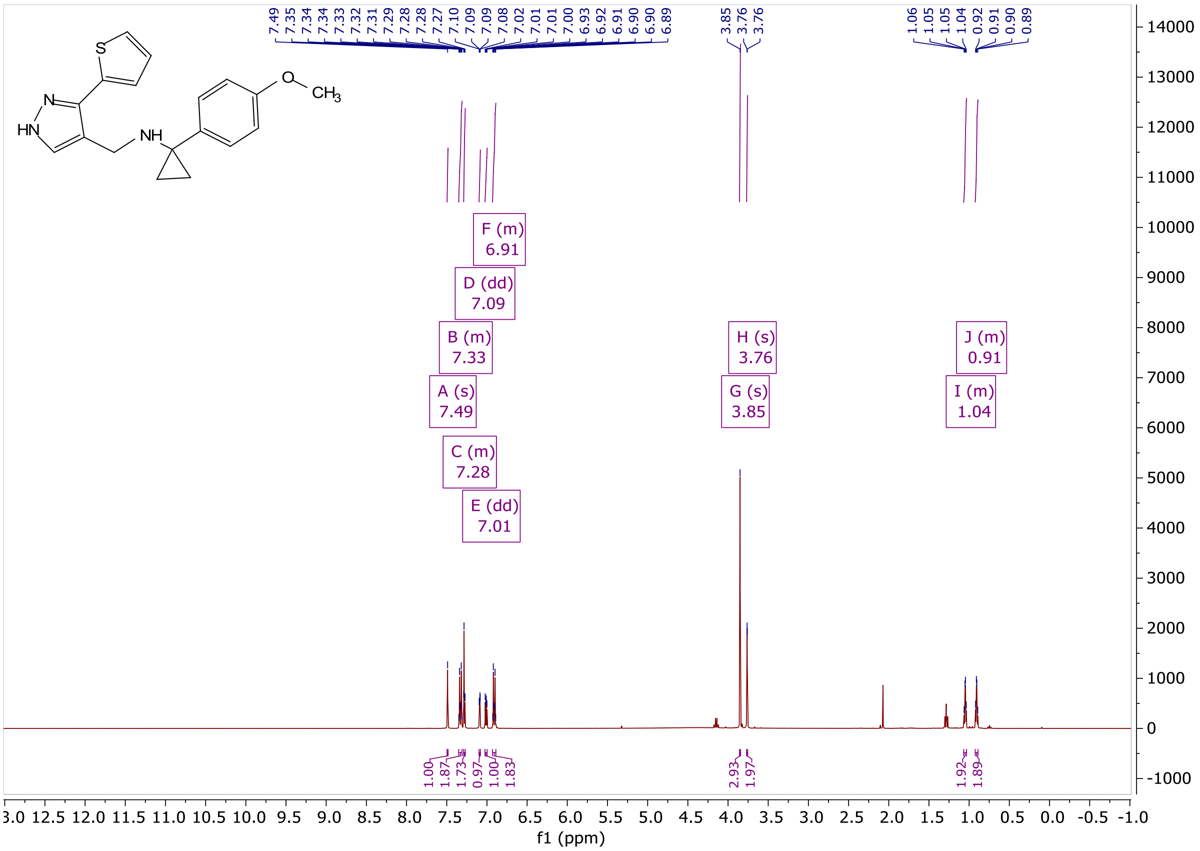

Figure **A11**: $^{1}$H NMR of **KNX-002**

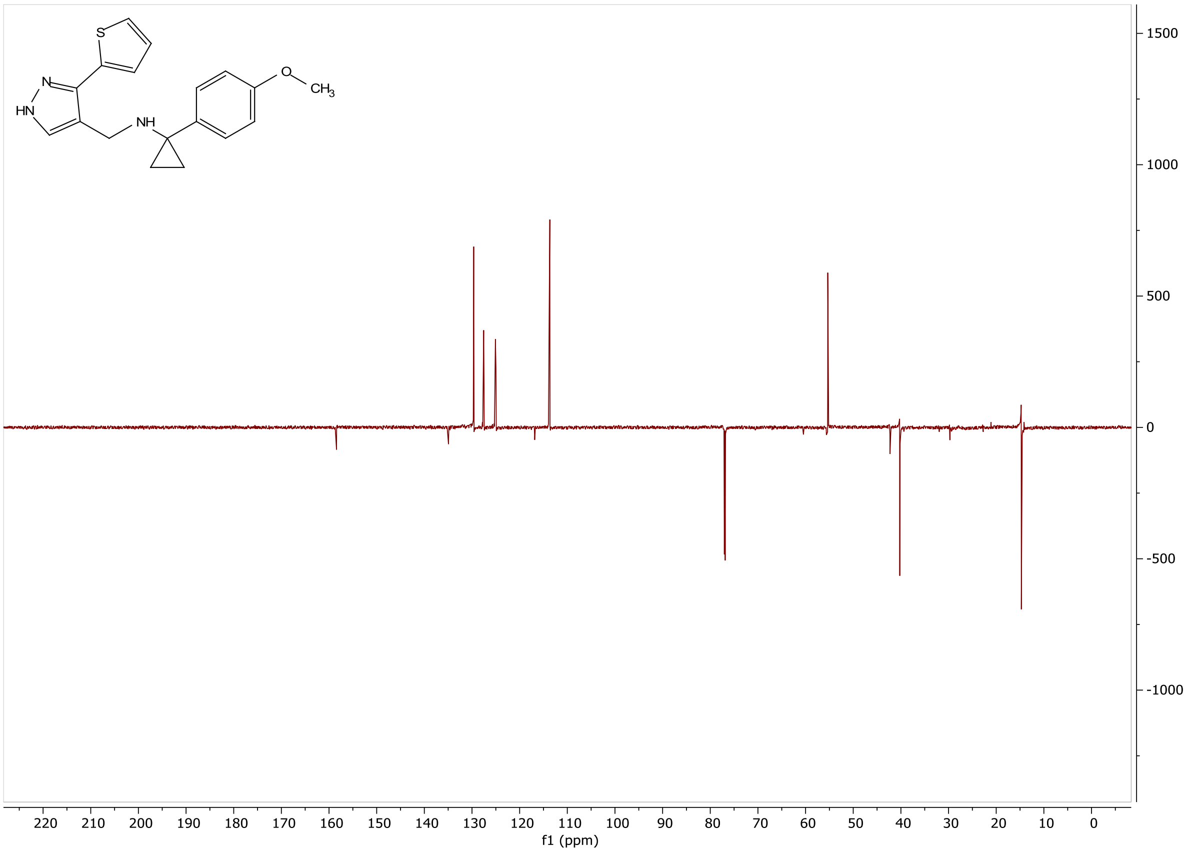


Figure **A12**: $^{13}$C NMR of **KNX-002**

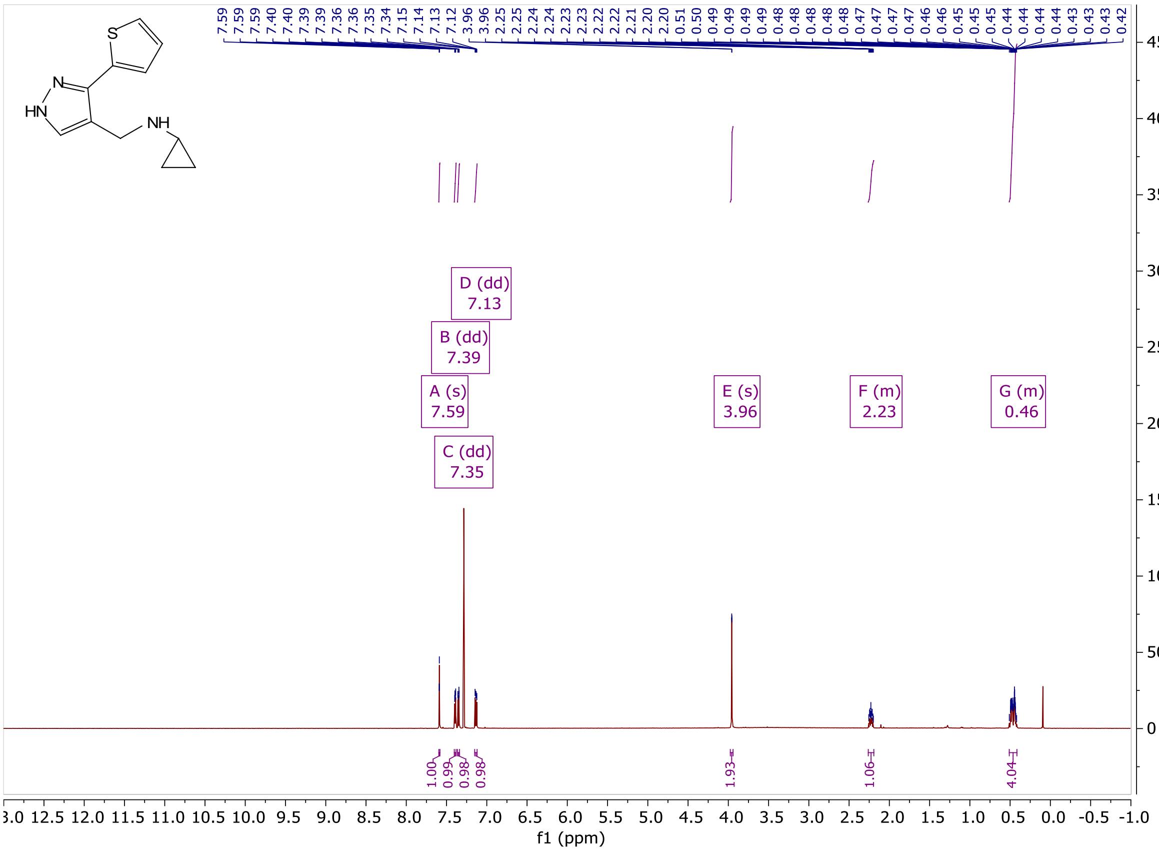


Figure **A13**: $^{1}$H NMR of **VEST1**

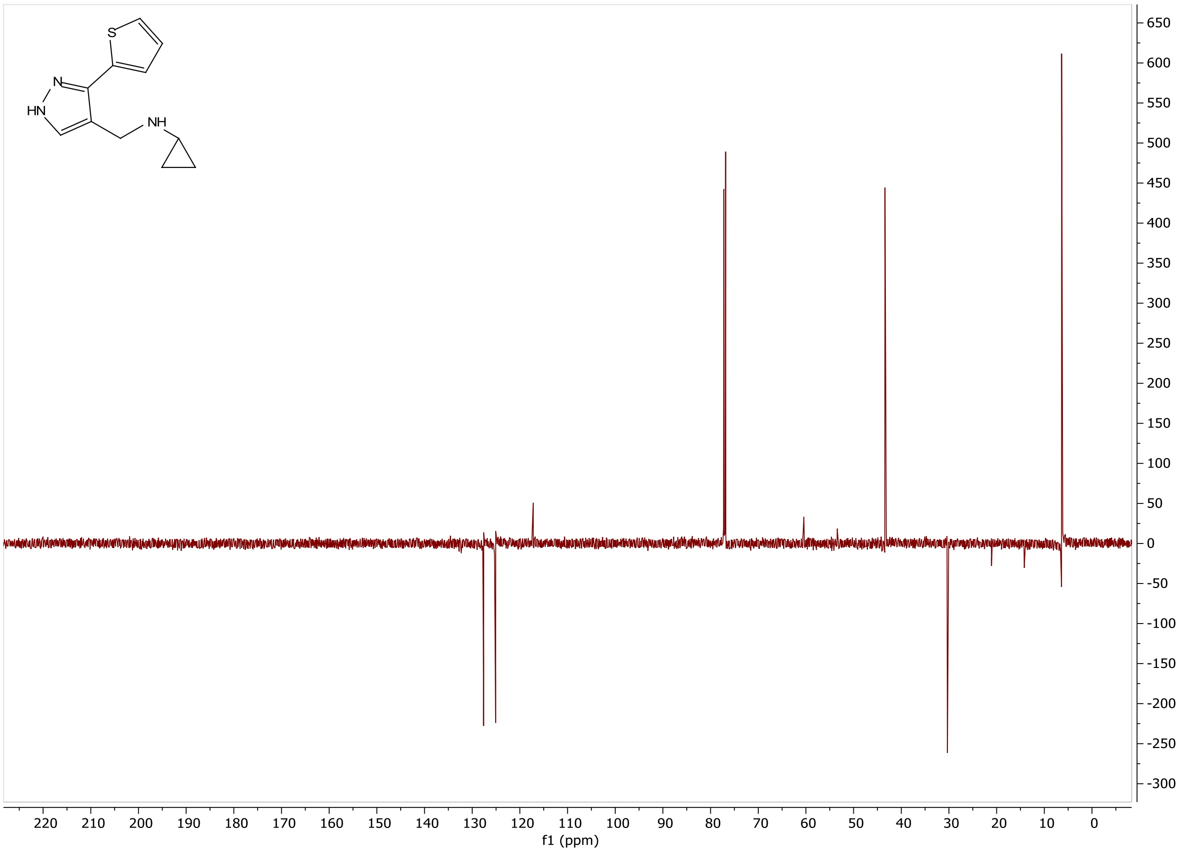


Figure **A14**: $^{13}$C NMR of **VEST1**

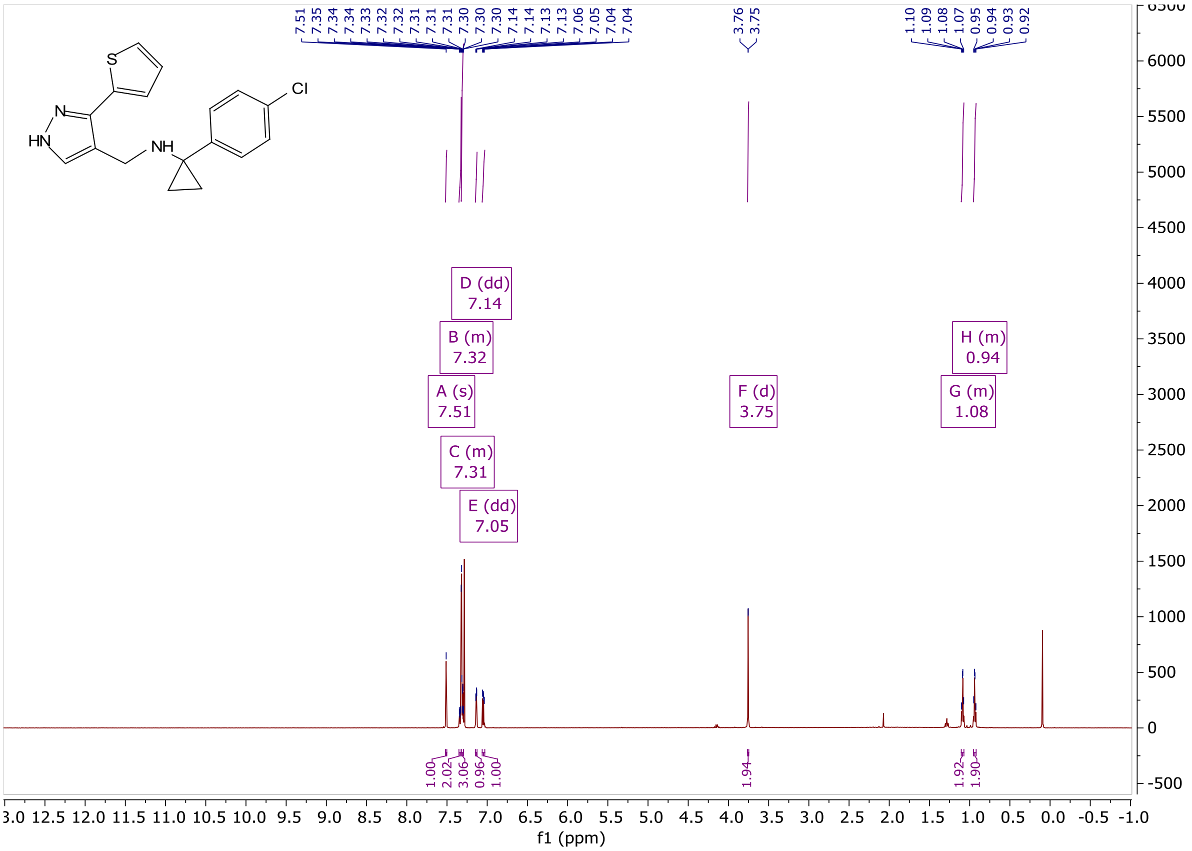


Figure **A15**: $^{1}$H NMR of **VEST2**

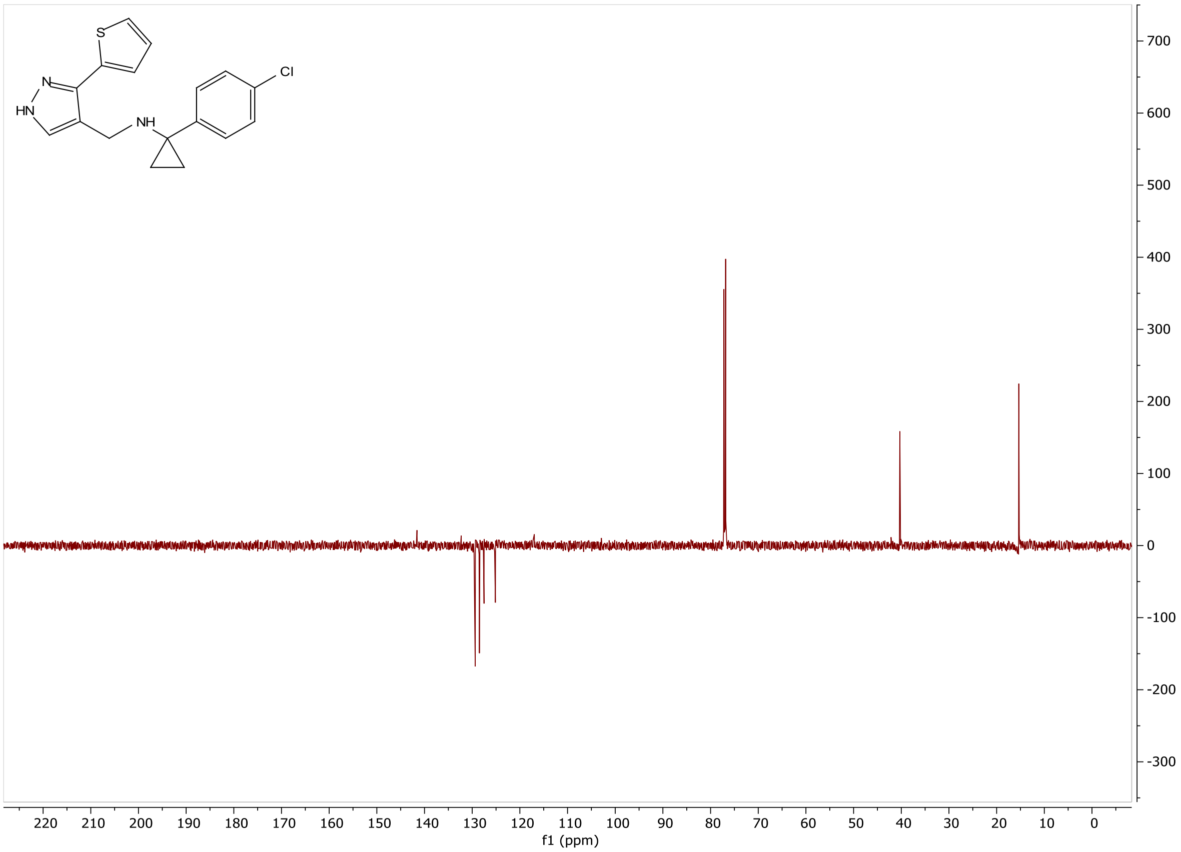

Figure **A16**: $^{13}$C NMR of **VEST2**

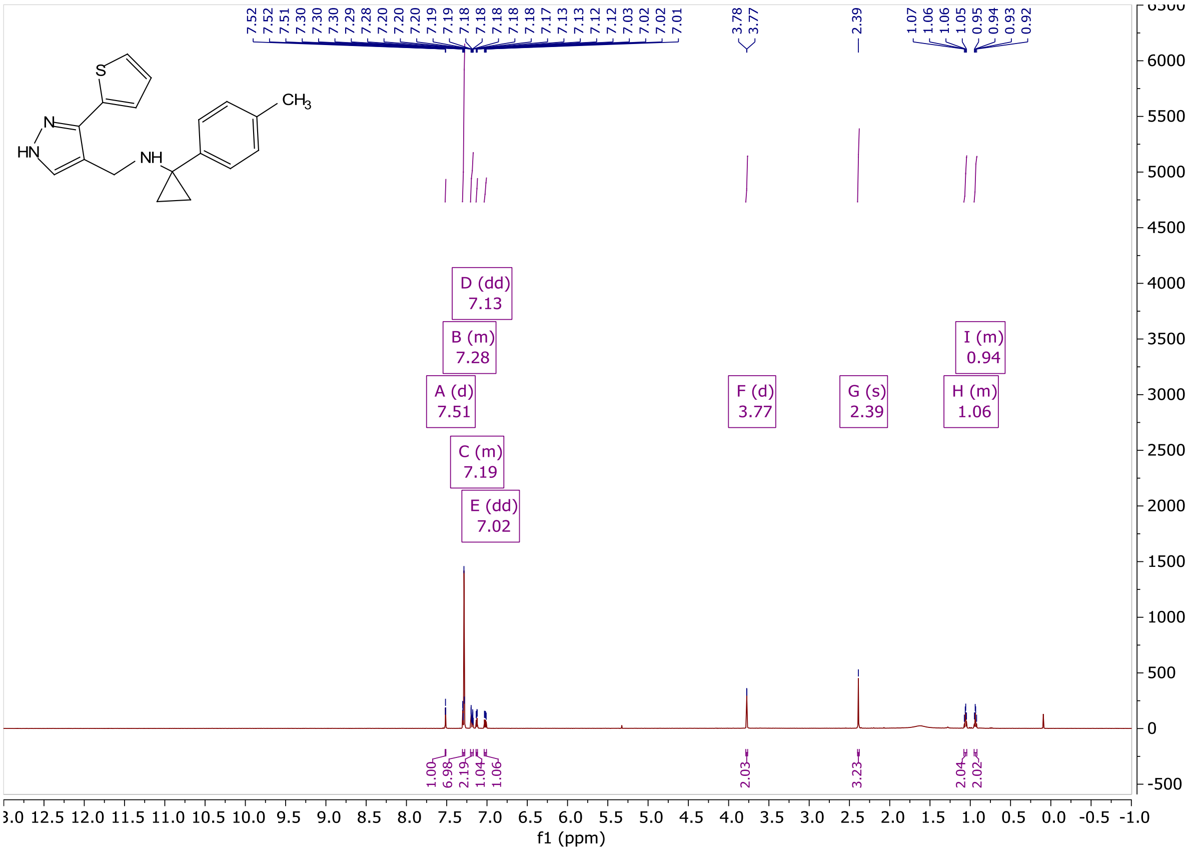

Figure **A17**: $^{1}$H NMR of **VEST3**

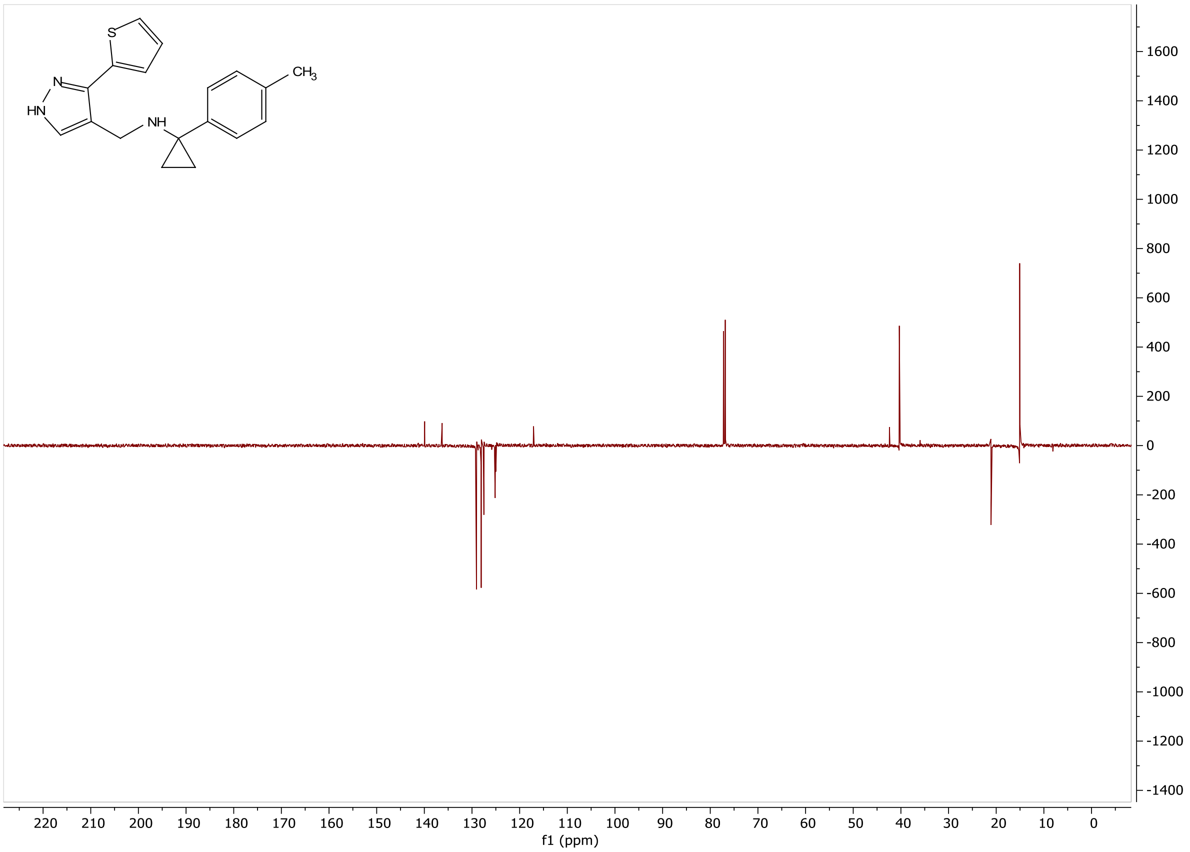

Figure **A18**: $^{13}$C NMR of **VEST3**

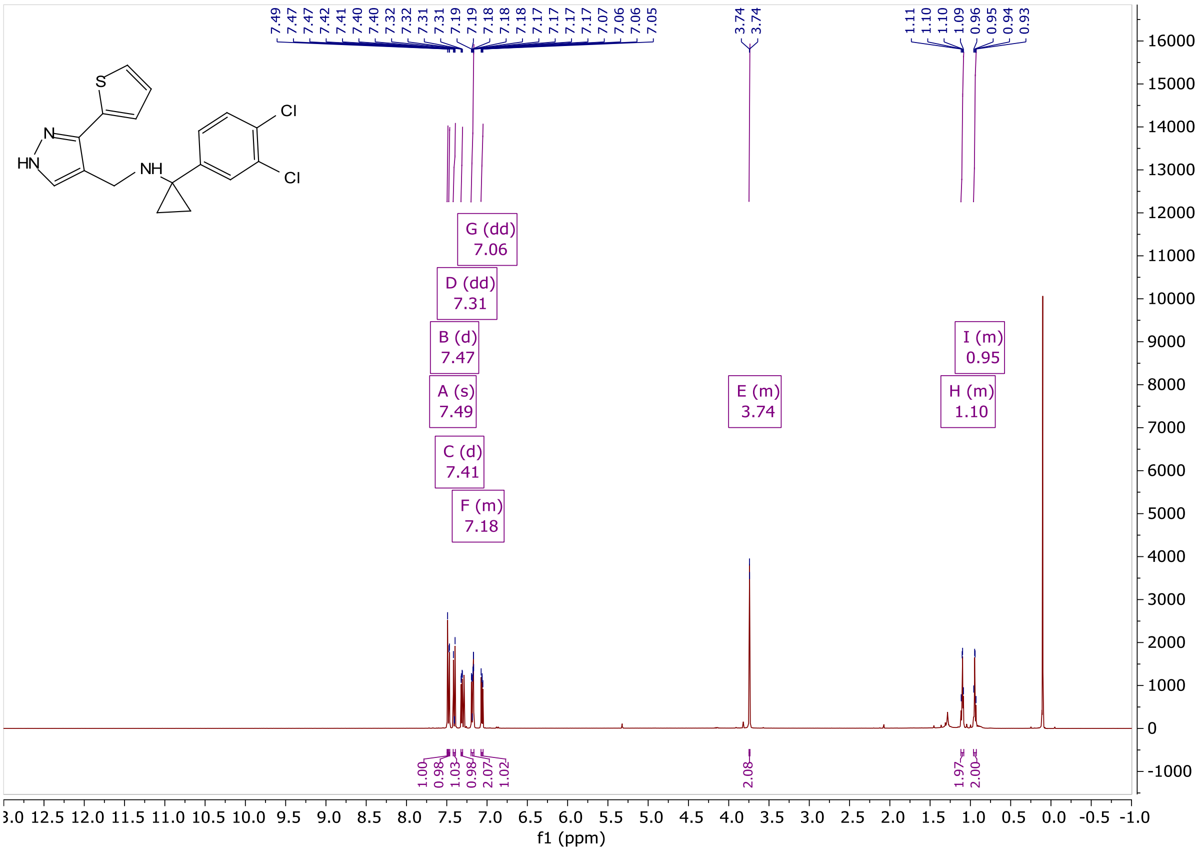


Figure **A19**: $^{1}$H NMR of **VEST4**

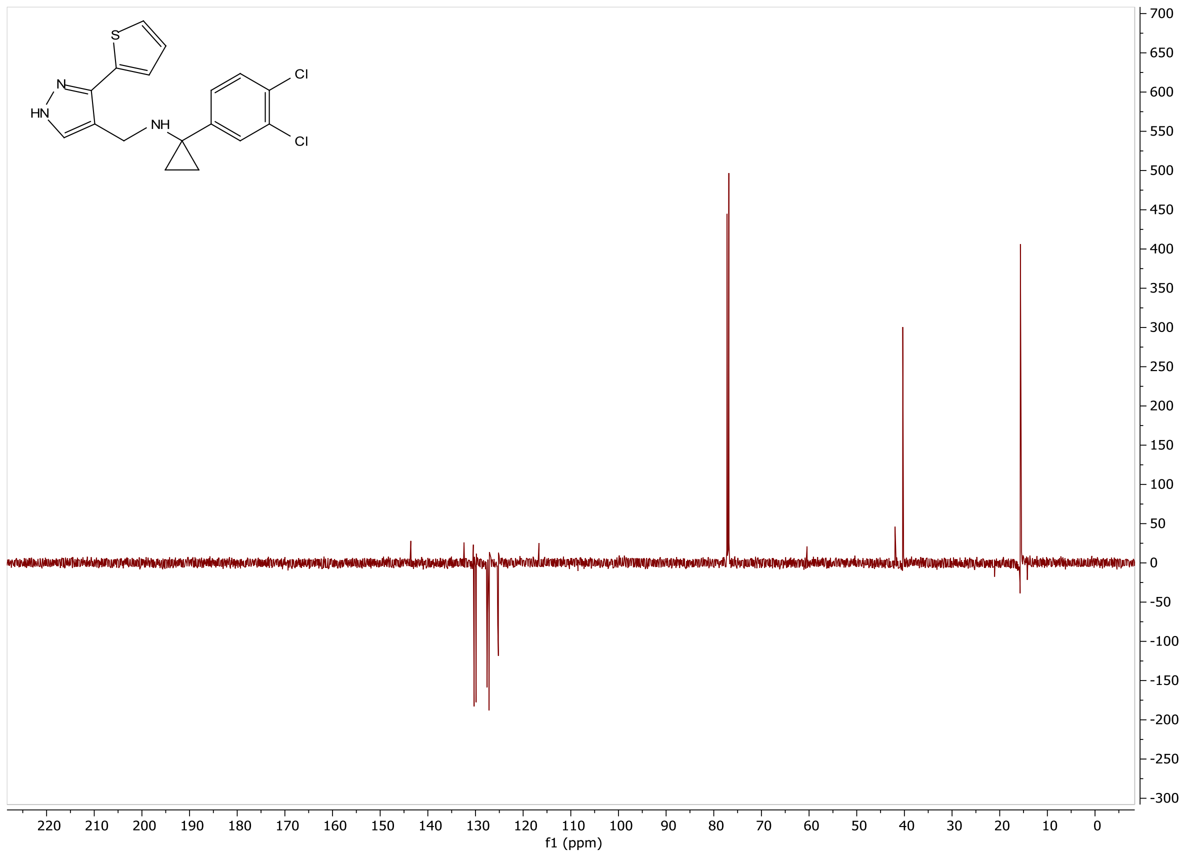

Figure **A20** [13]C NMR of **VEST4**

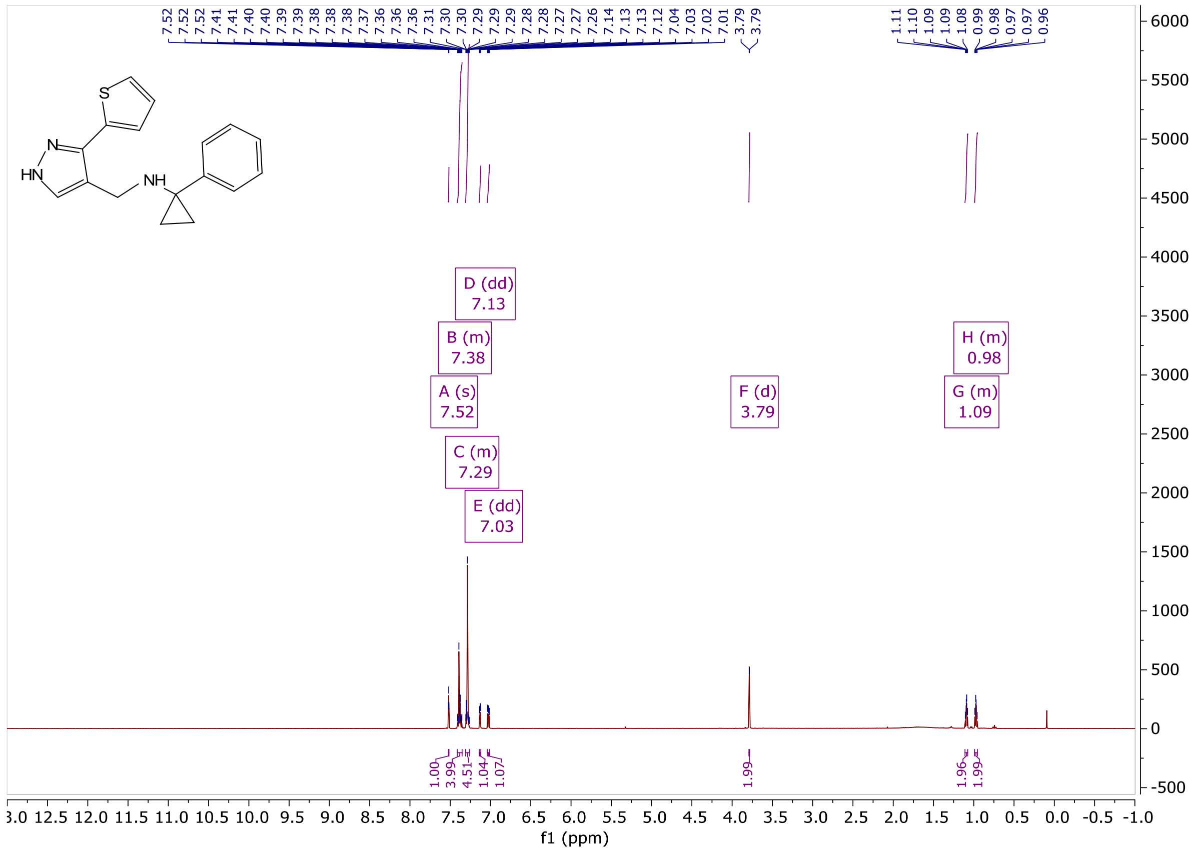

Figure **A21**: [1]H NMR of **VEST5**

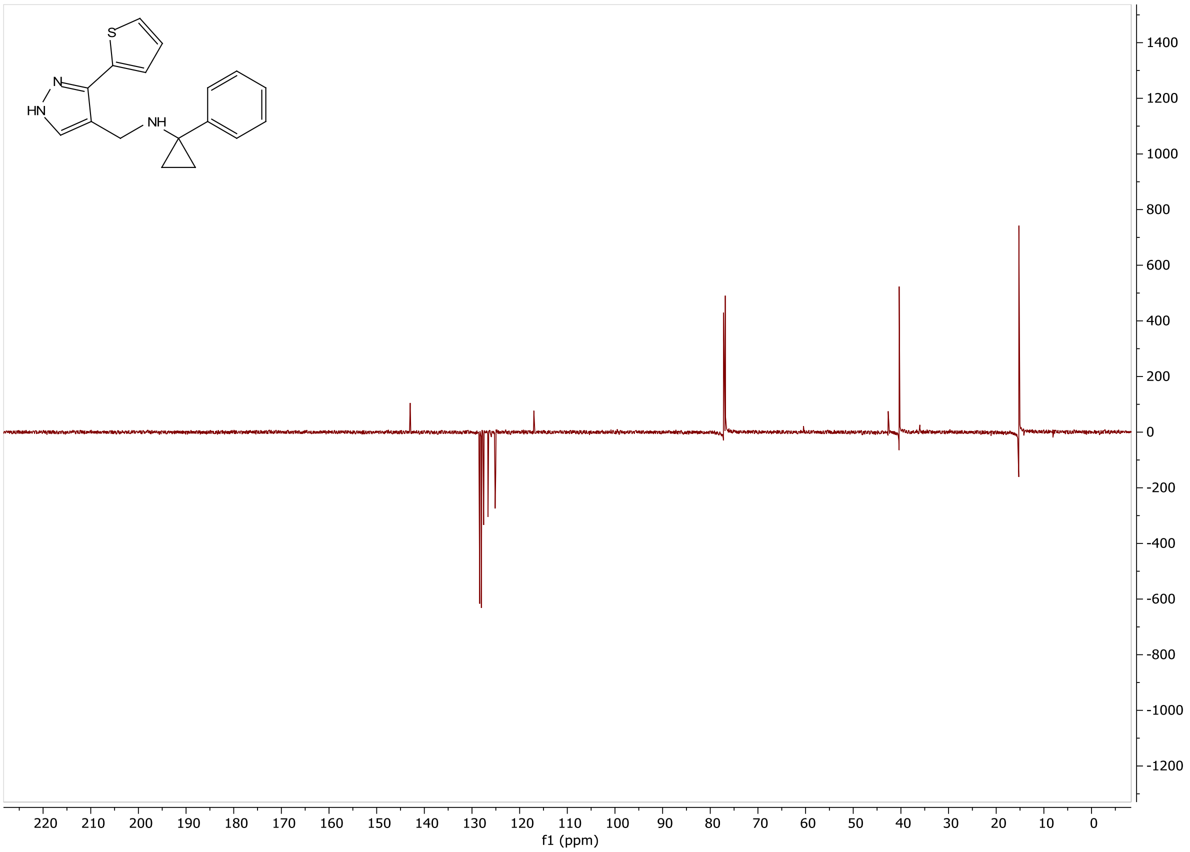


Figure **A22**: $^{13}$C NMR of **VEST5**

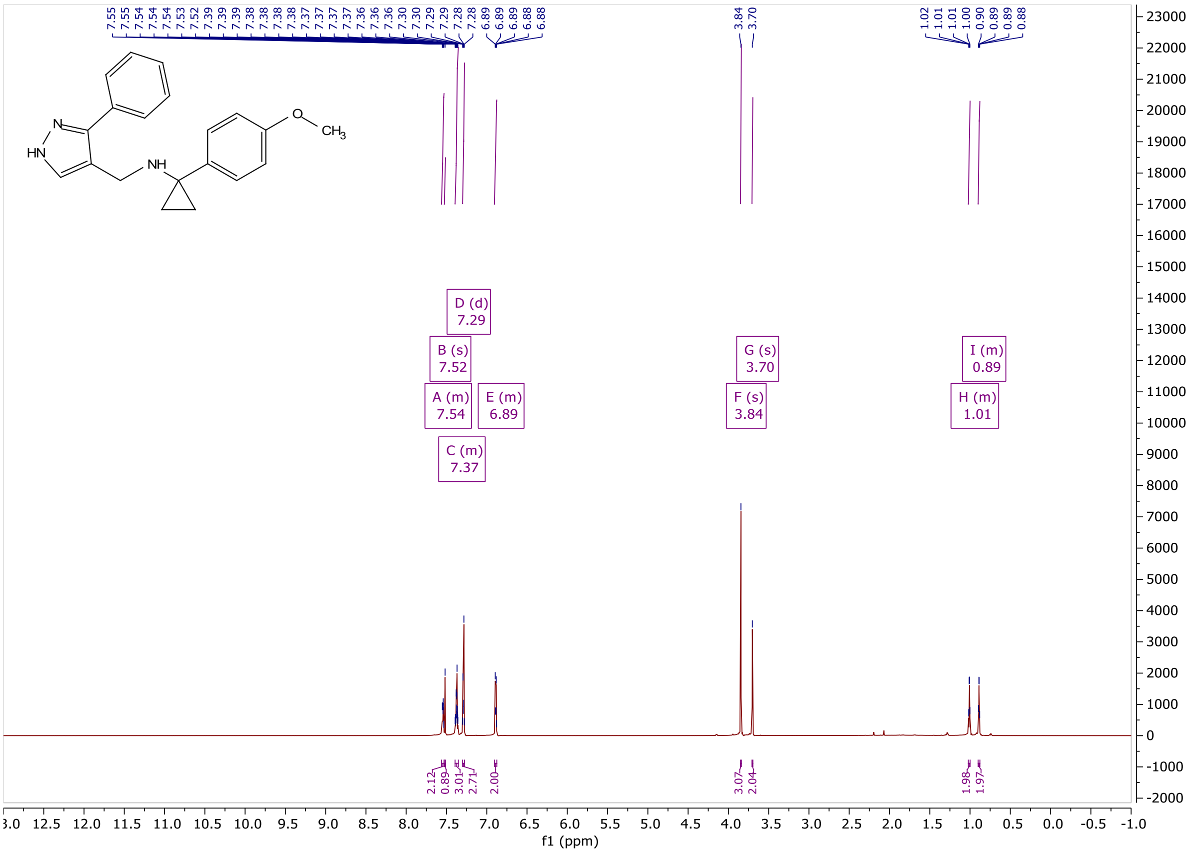


Figure **A23**: $^{1}$H NMR of **VEST6**

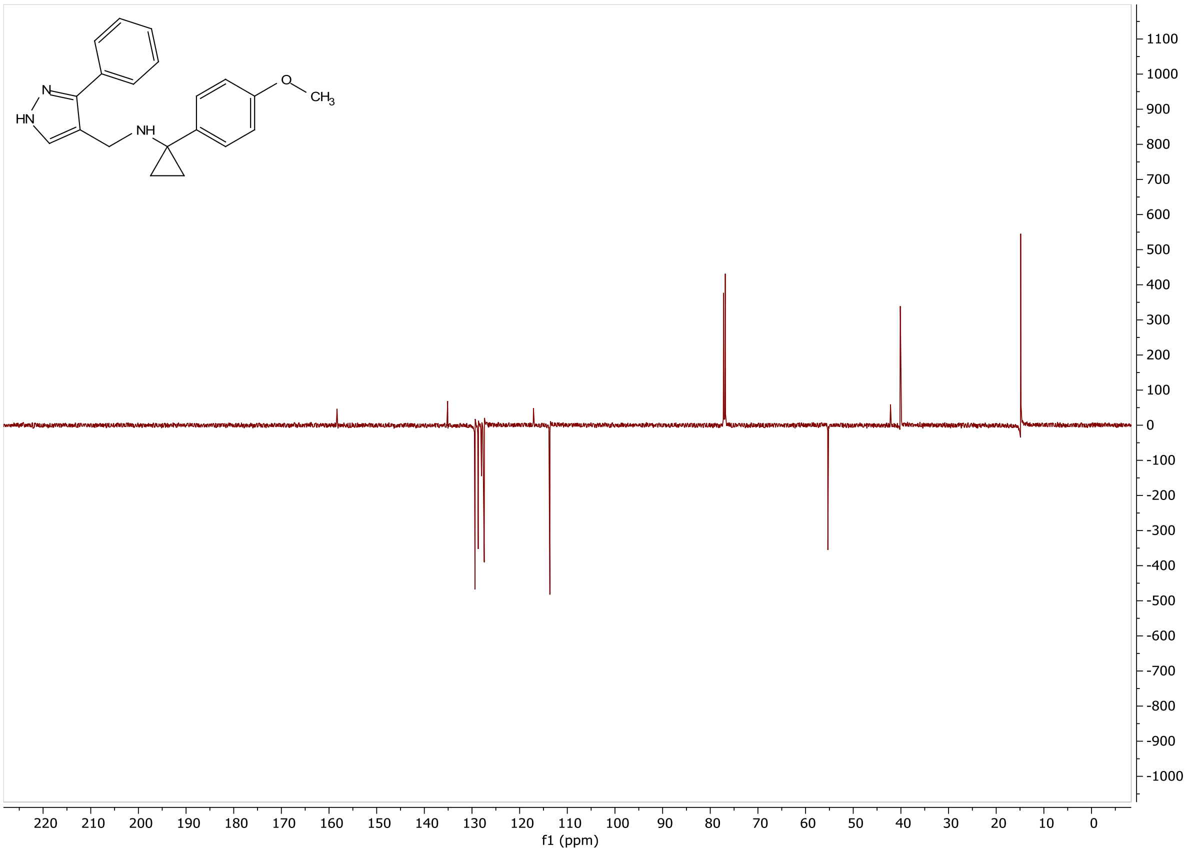


Figure **A24**: $^{13}$C NMR of **VEST6**

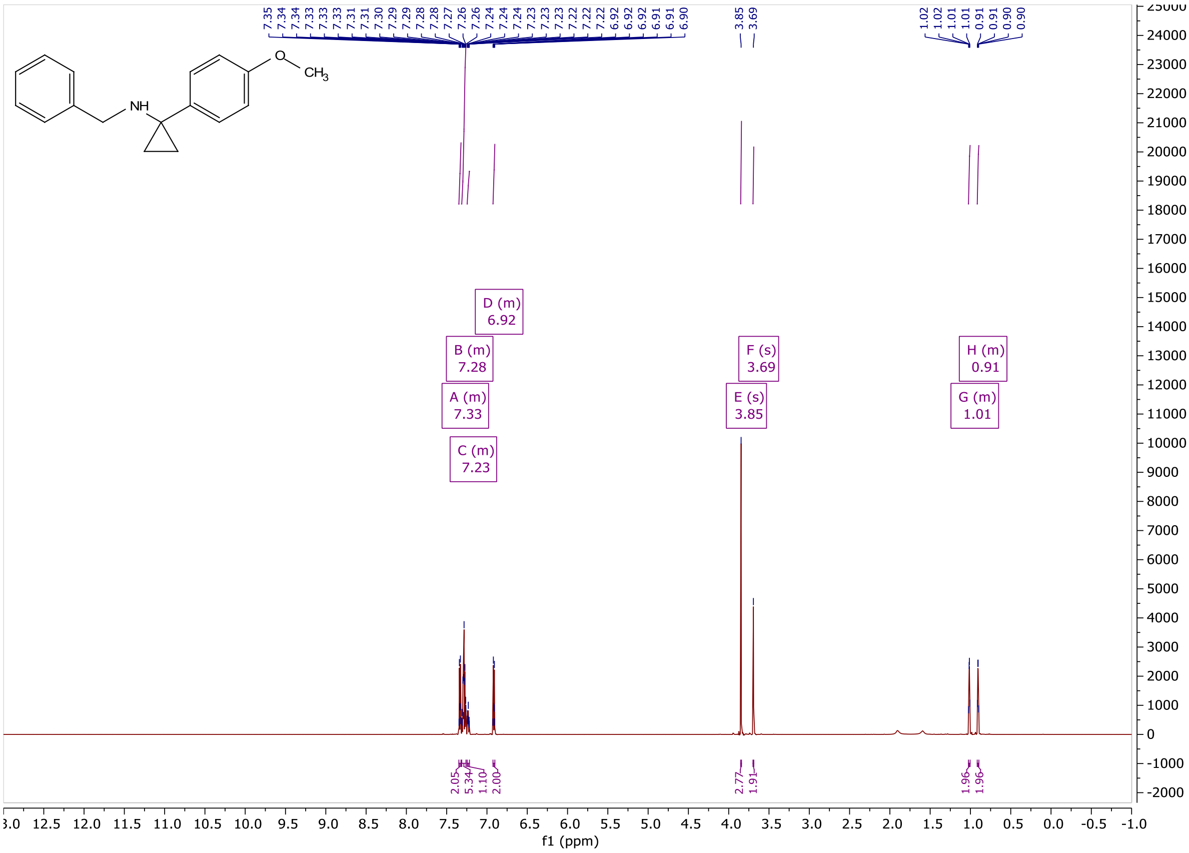


Figure **A25**: $^{1}$H NMR of **VEST7**

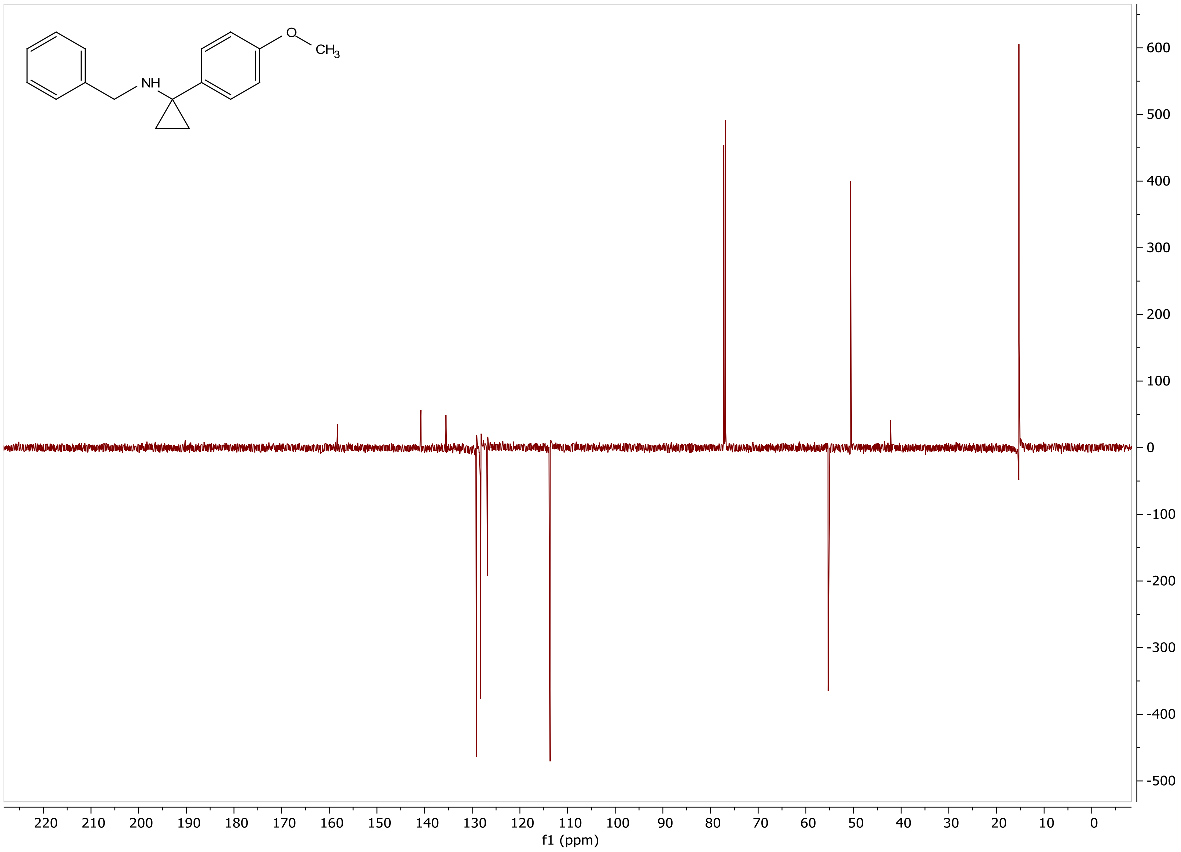


Figure **A26**: $^{13}$C NMR of **VEST7**

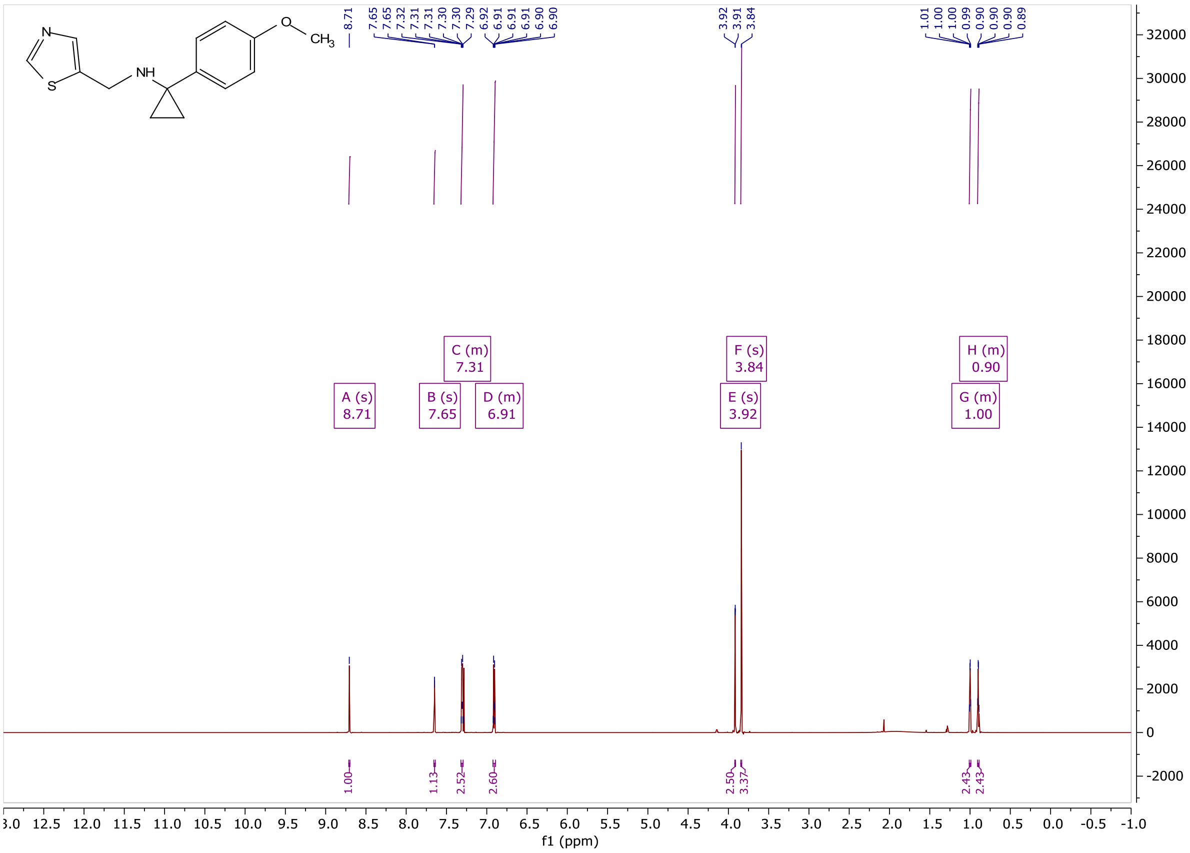


Figure **A27**: $^{1}$H NMR of **VEST8**

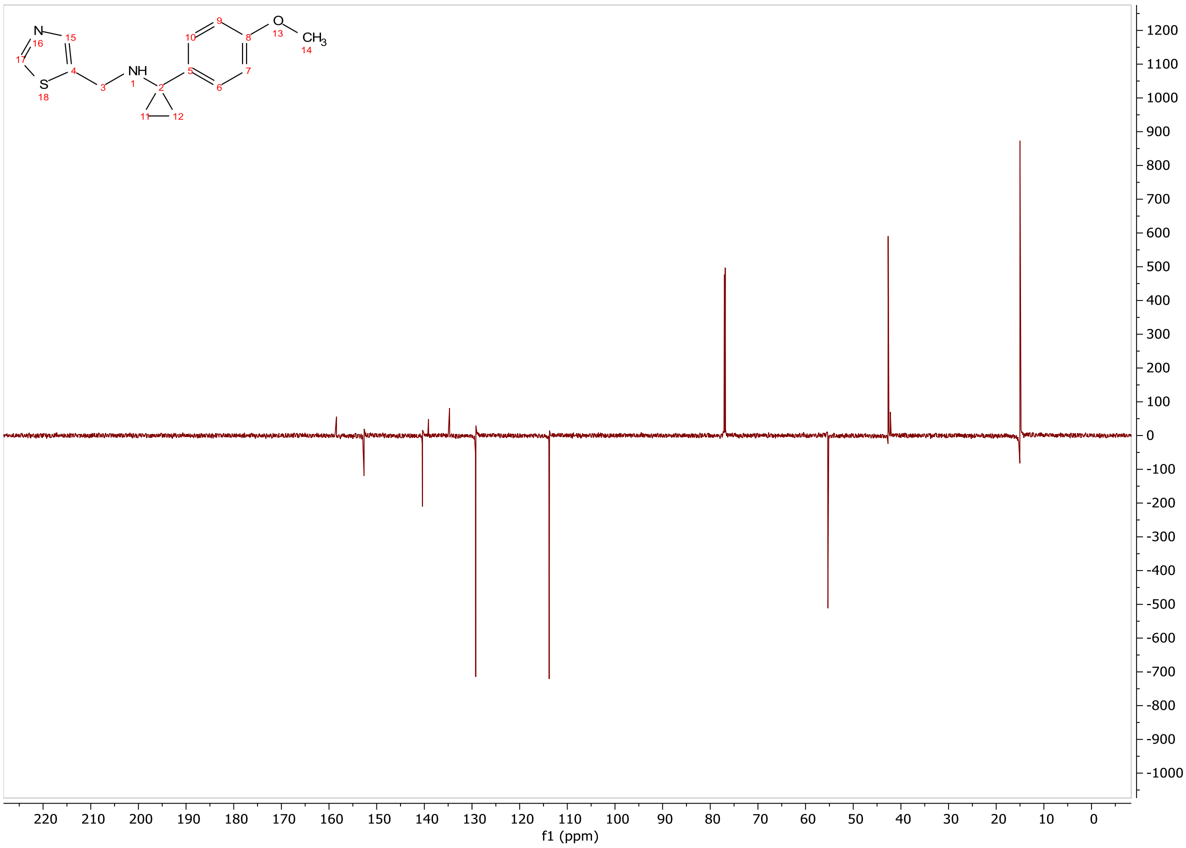

Figure **A28**: $^{13}$C NMR of **VEST8**

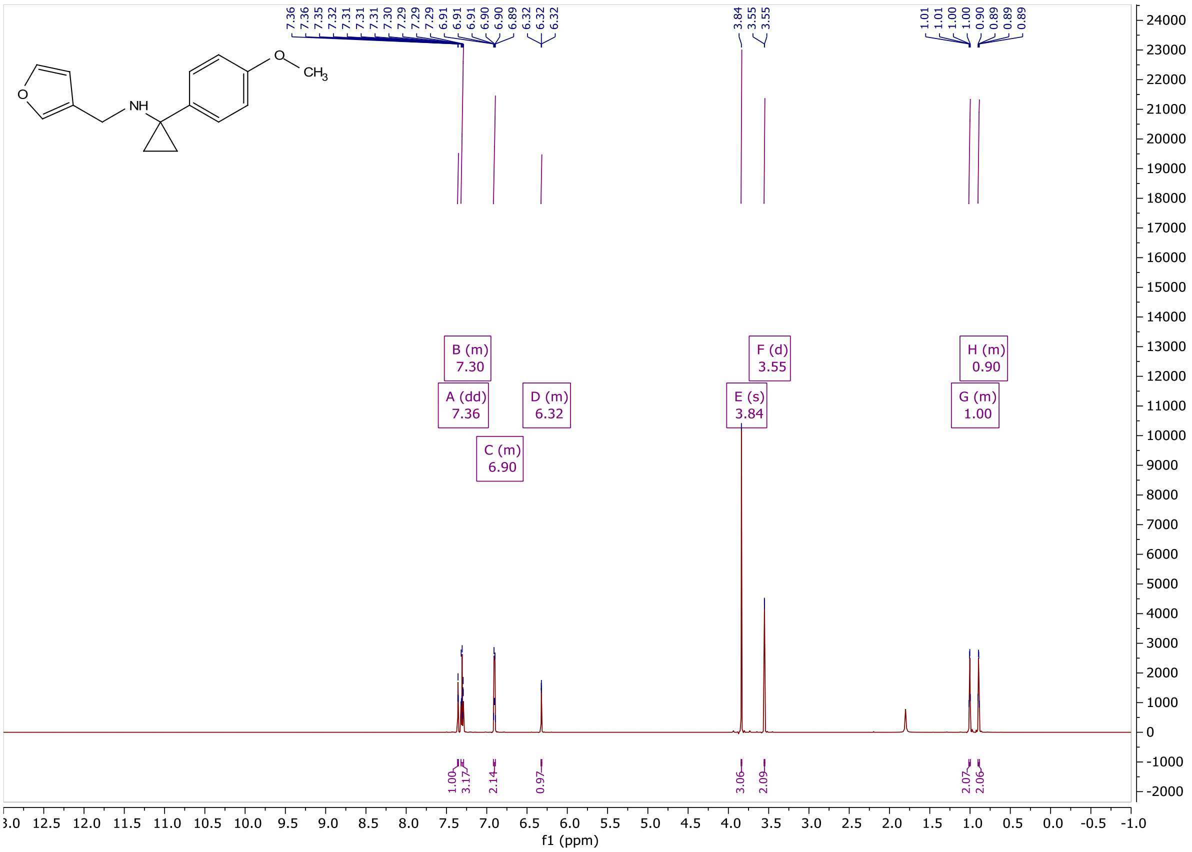

Figure **A29**: $^{1}$H NMR of **VEST9**

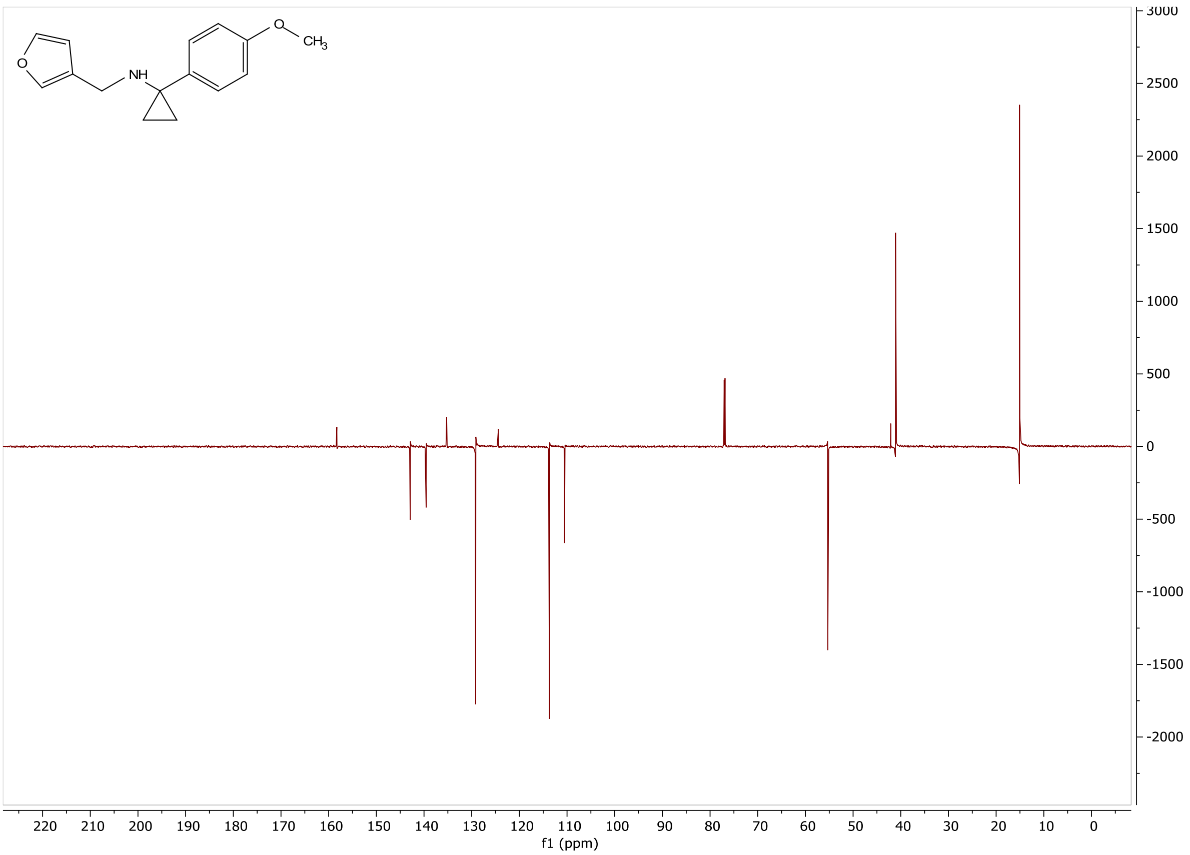

Figure **A30**: $^{13}$C NMR of **VEST9**

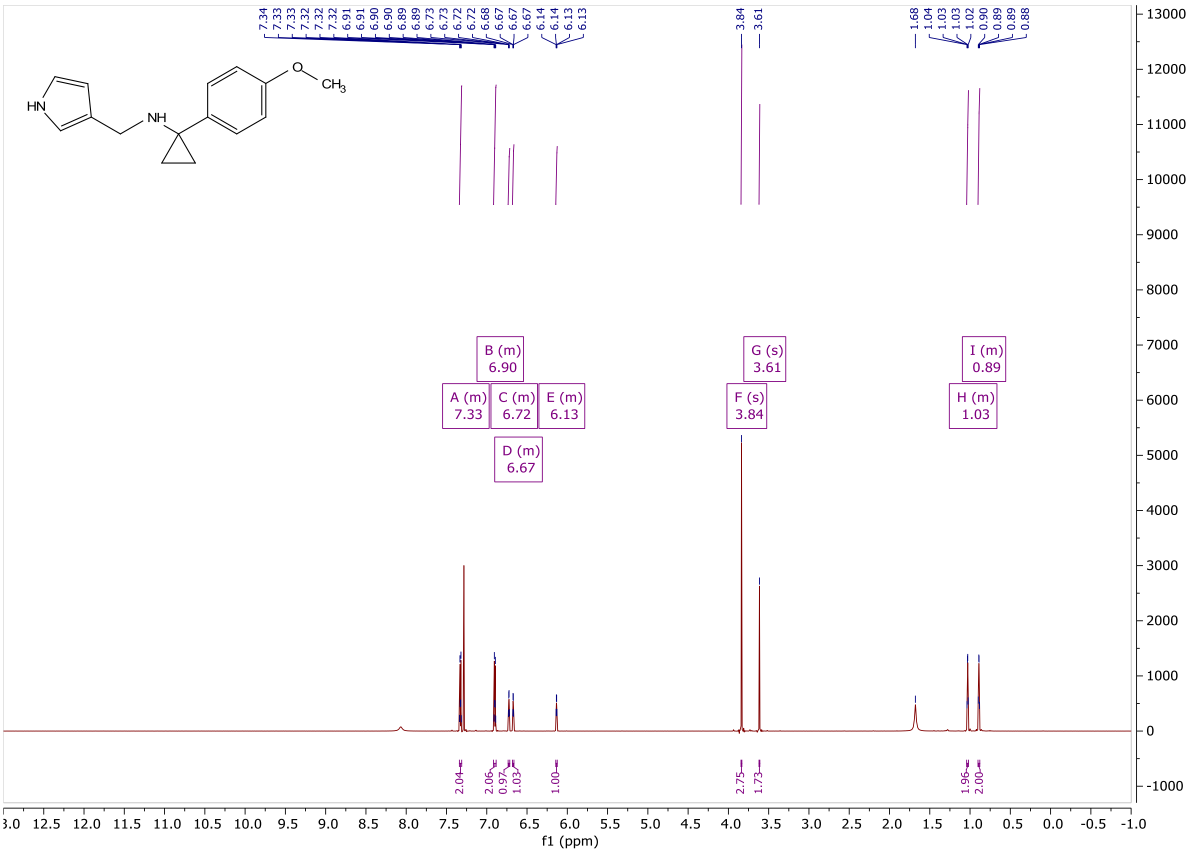

Figure **A31**: $^{1}$H NMR of **VEST10**

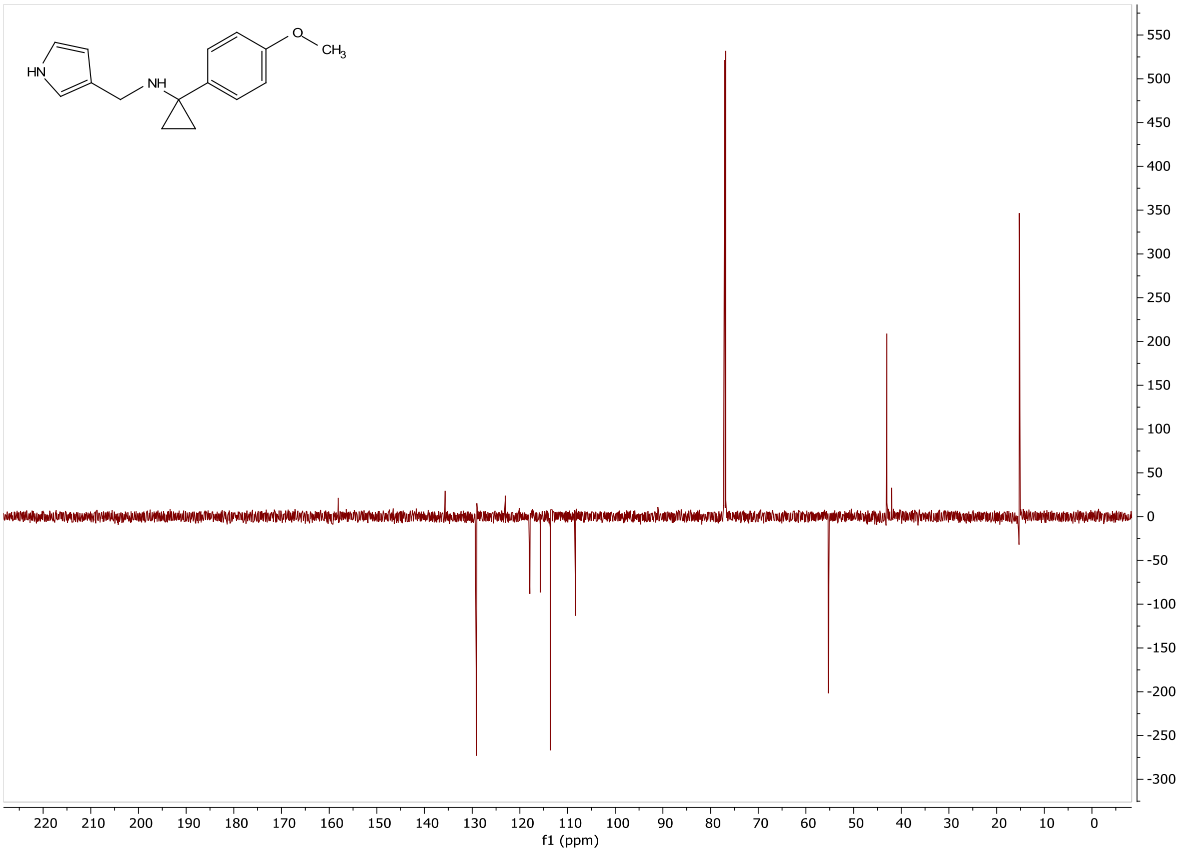


Figure **A32**: $^{13}$C NMR of **VEST10**

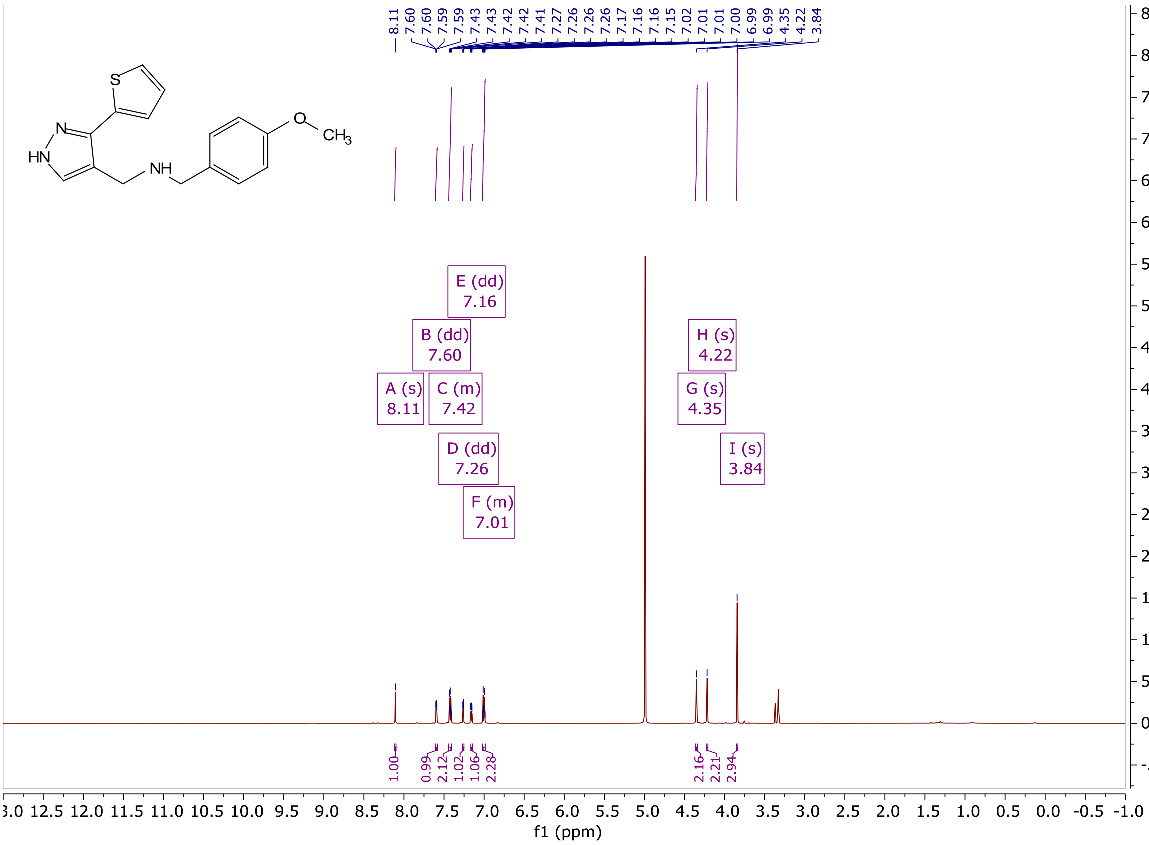


Figure **A33**: $^{1}$H NMR of **VEST11**

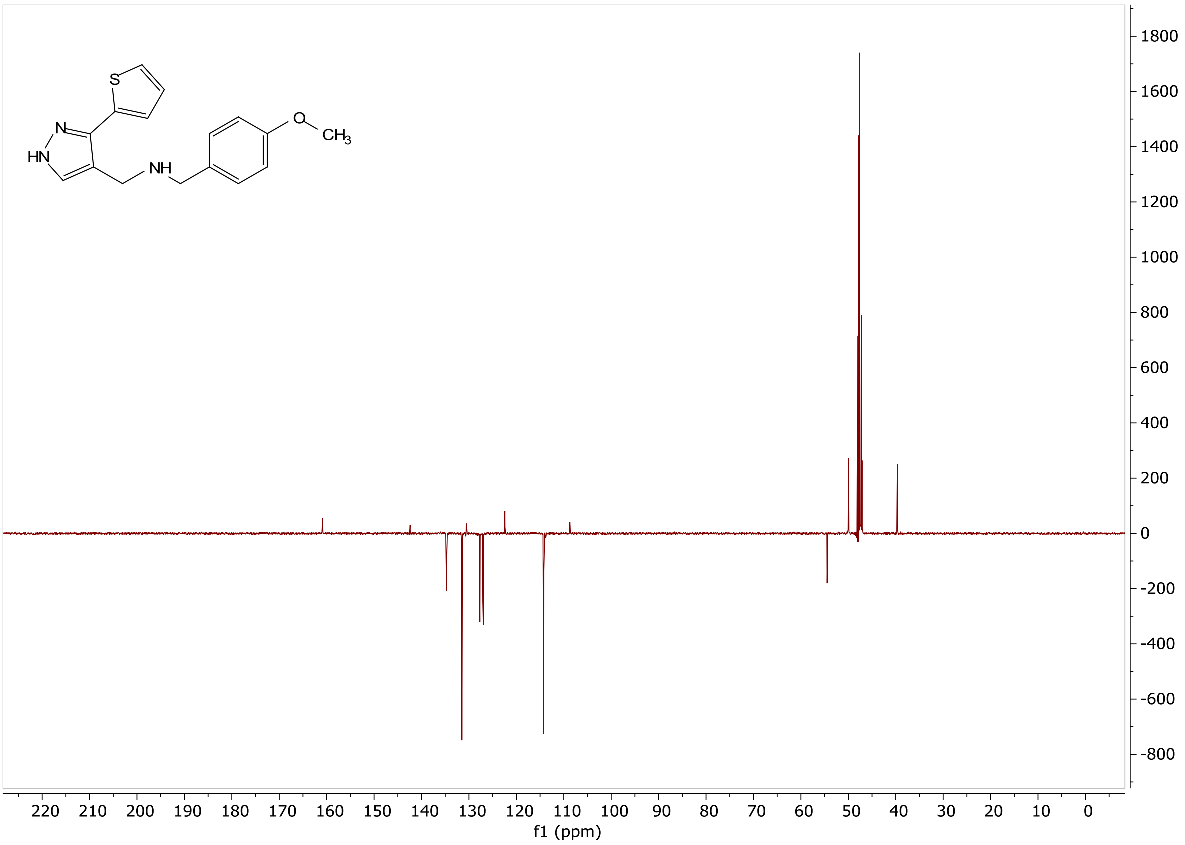


Figure **A34**: $^{13}$C NMR of **VEST11**

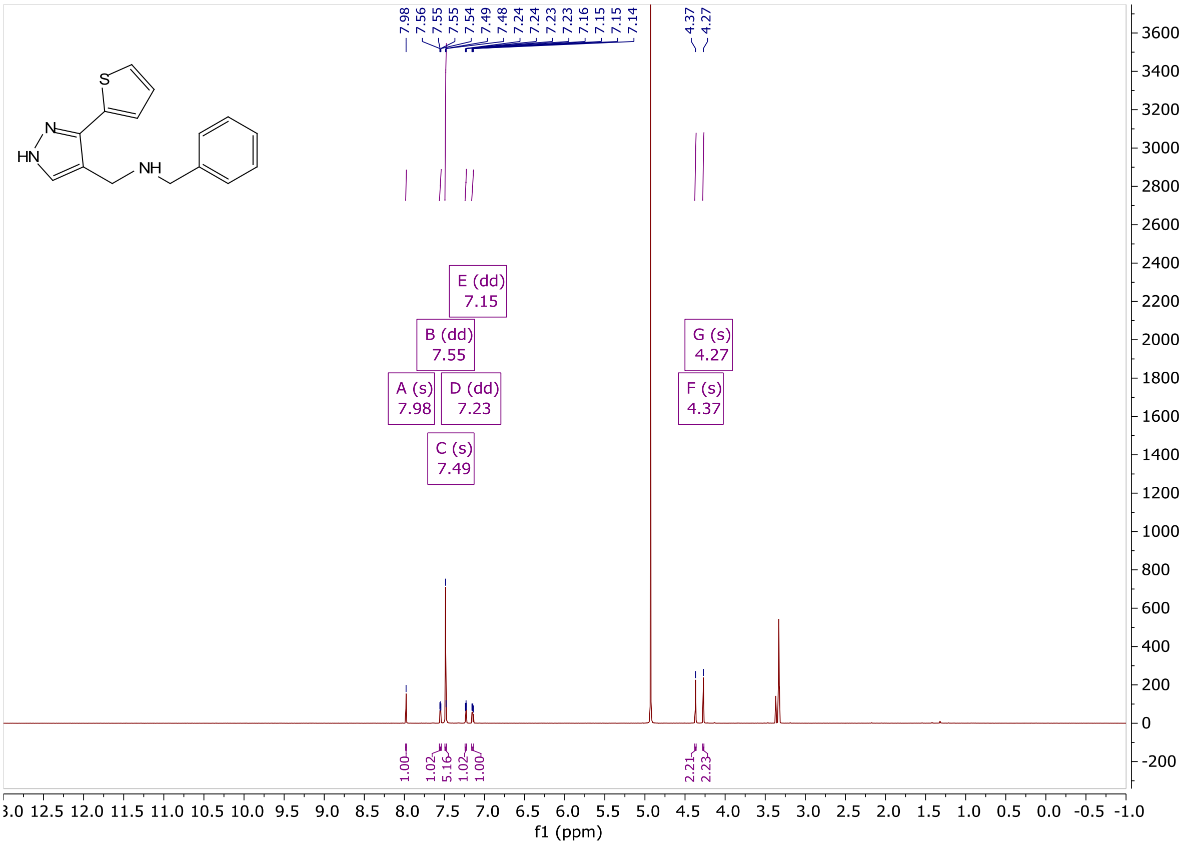


Figure **A35**: $^{1}$H NMR of **VEST12**

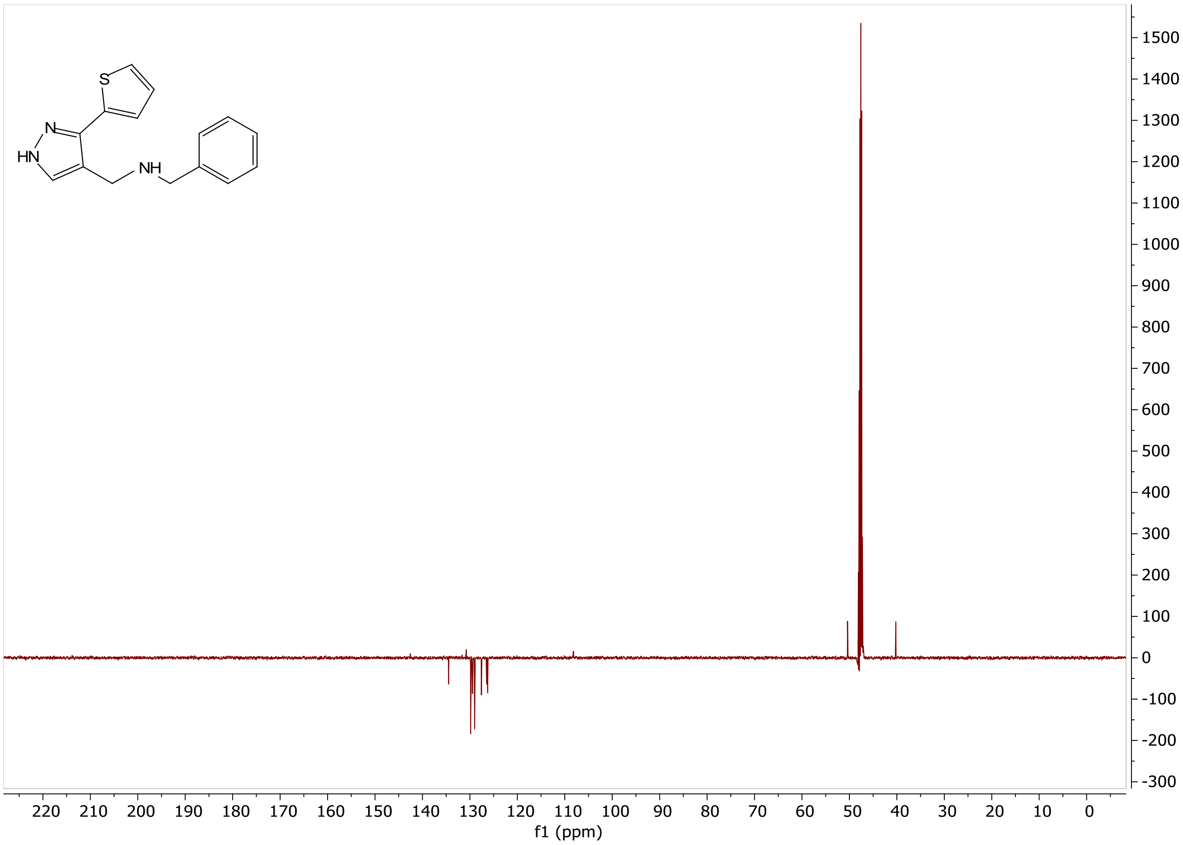


Figure **A36**: $^{13}C$ NMR of **VEST12**

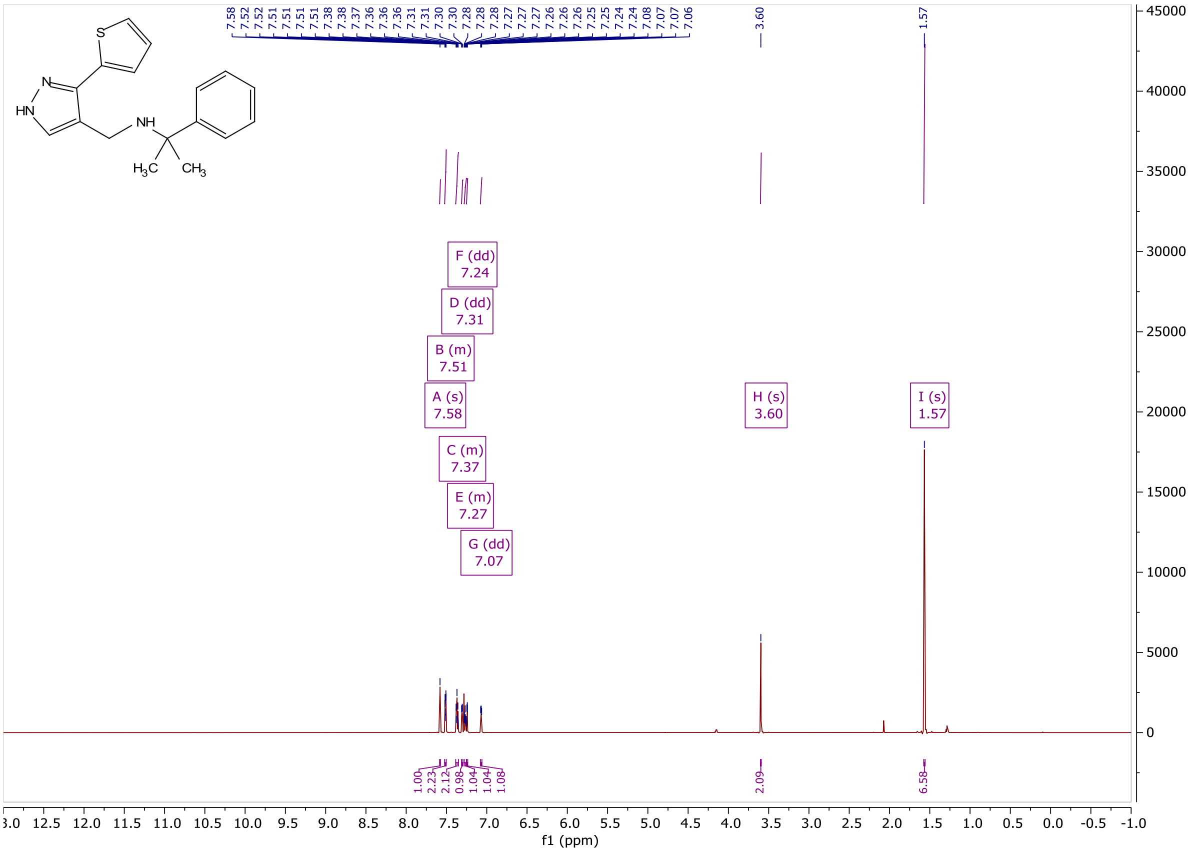


Figure **A37**: $^{1}H$ NMR of **VEST13**

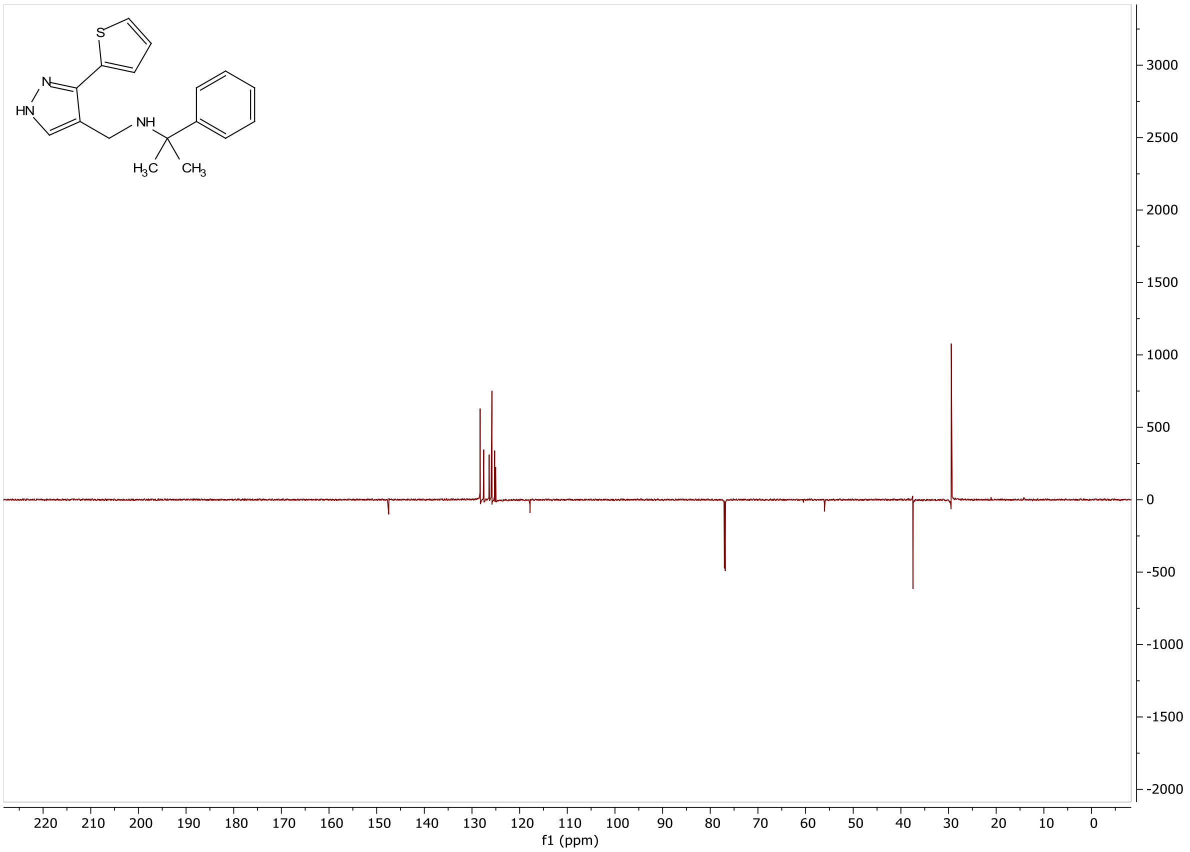


Figure **A38**: $^{13}C$ NMR of **VEST13**

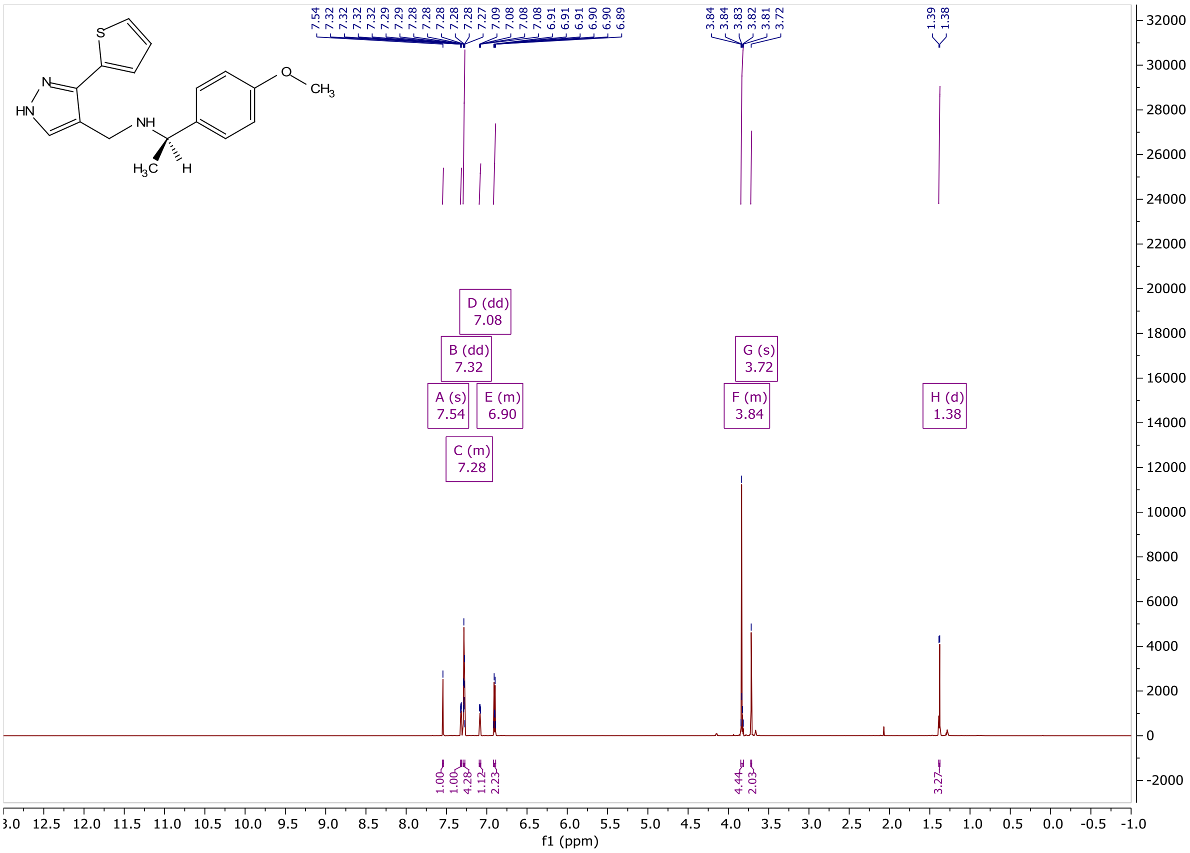


Figure **A39**: $^{1}H$ NMR of **VEST14**

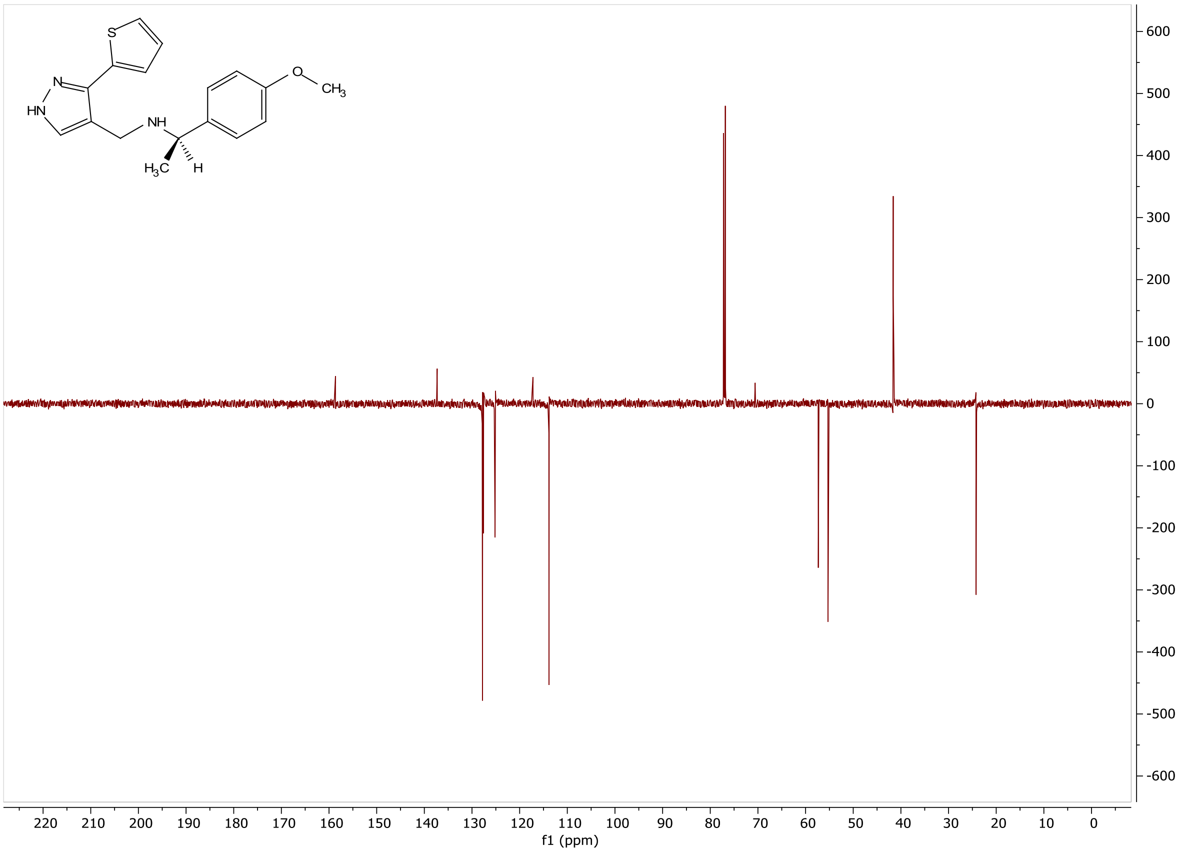


Figure **A40**: $^{13}$C NMR of **VEST14**

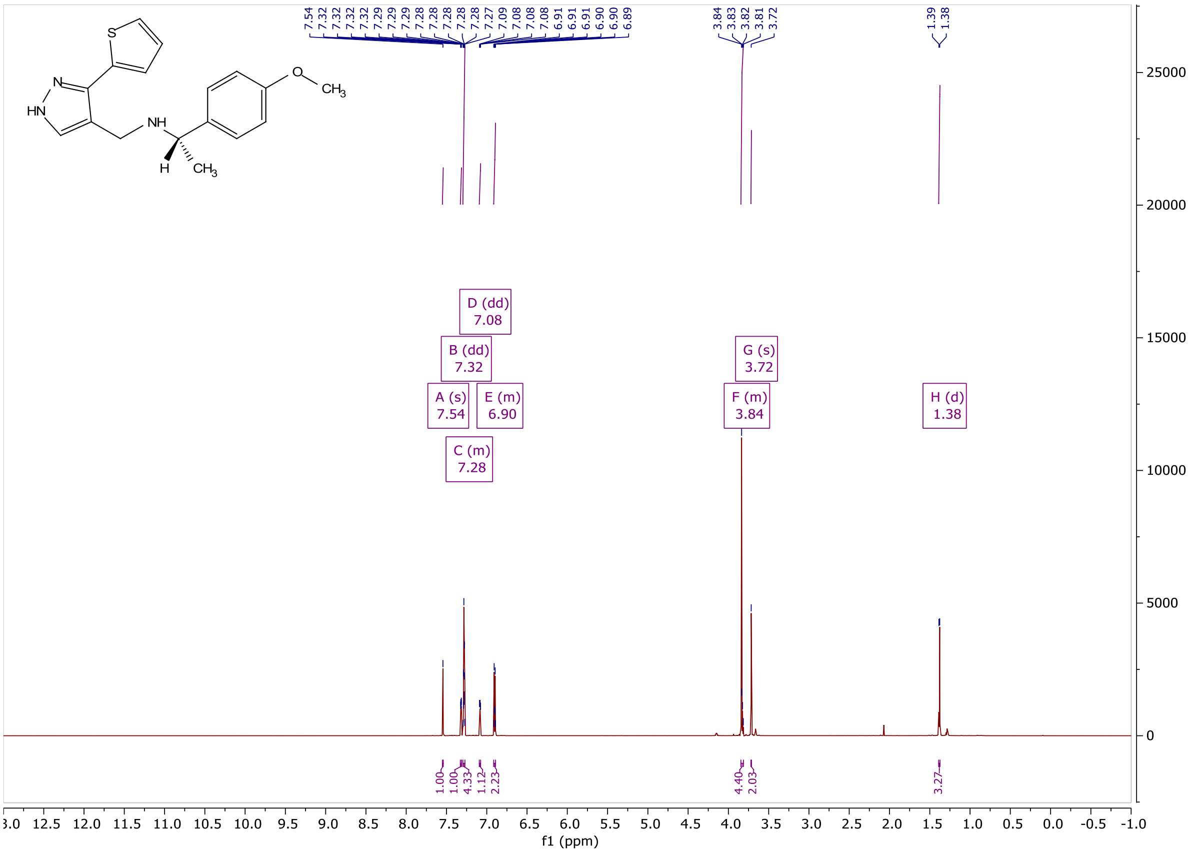


Figure **A41**: $^{1}$H NMR of **VEST15**

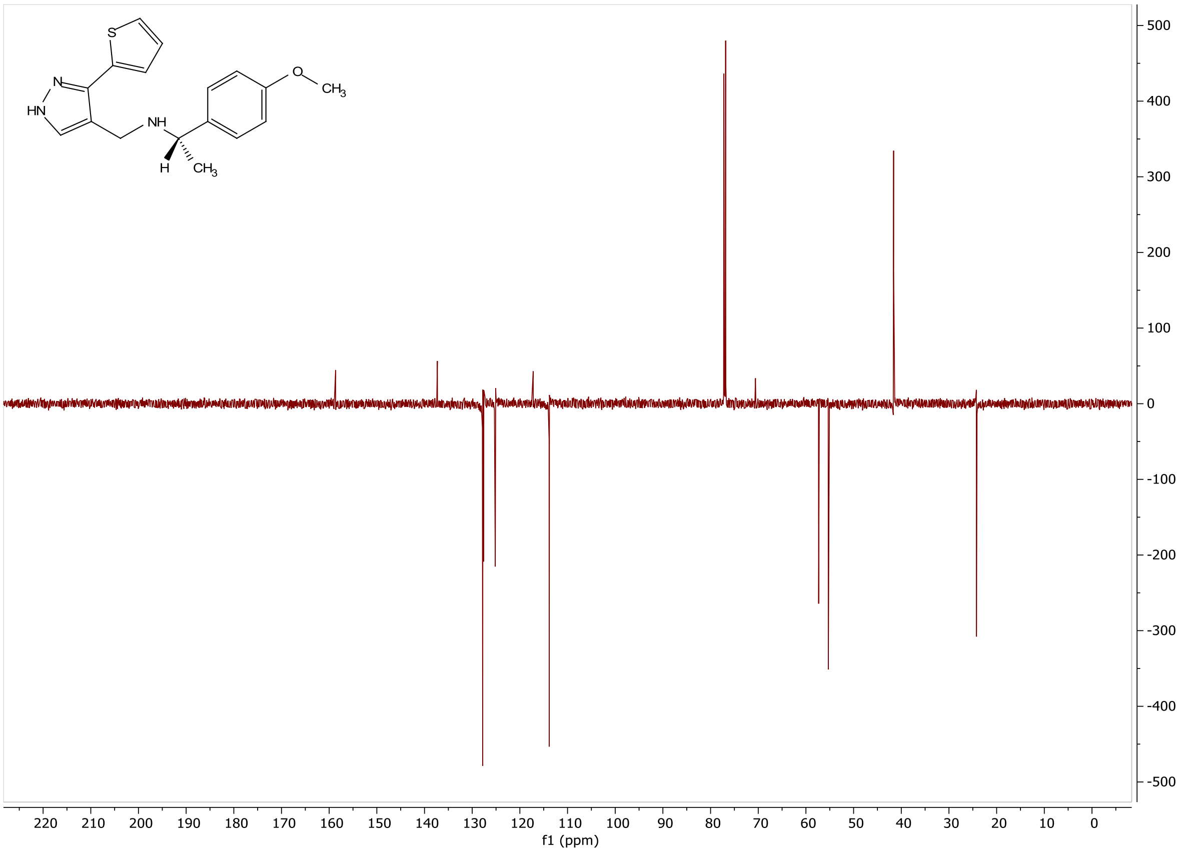


Figure **A42**: $^{13}$C NMR of **VEST15**
