## Supplementary material for "The class XIV myosin of *Toxoplasma gondii*, TgMyoA, is druggable in an animal model of infection": Suppl. Figures 1-14

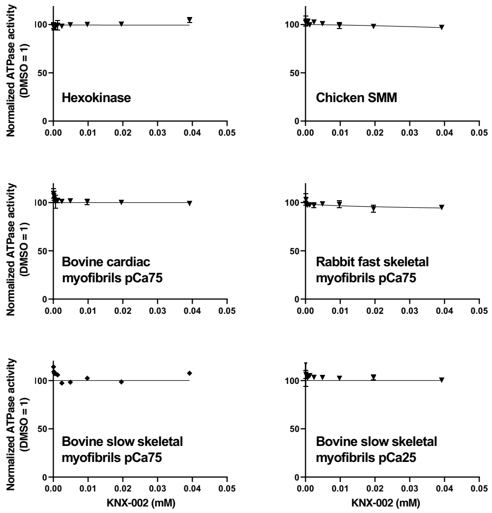


<u>**Suppl. Figure 1**</u>**: KNX-002 has little to no effect on the activity of various vertebrate myosins.** Dose-response curves showing the effect of KNX-002 on the actin-activated ATPase activity of various vertebrate myosins. SMM = smooth muscle myosin; pCa25 and pCa75 refer to assays done at 25% and 75% calcium activation, respectively.

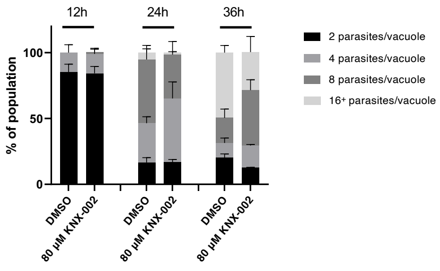


**<u>Suppl. Figure 2</u>: KNX-002 has no effect on parasite intracellular replication.** Newly infected HFF cells were incubated at 37ºC in culture medium containing 80 µM KNX-002 or an equivalent volume of DMSO (vehicle) for the indicated times. The cells were then fixed and processed for immunofluorescence using an antibody against the inner membrane complex marker IMC1, enabling the number of parasites in each of 100-120 vacuoles to be counted. Shading of the bars indicates number of parasites per vacuole (see key); the means ± SEM at each time point of three biological replicates are shown. There were no significant differences between the number of DMSO- and compound-treated parasites per vacuole at each of the time points, based on unpaired Student's t-tests using a 2-step Benjamini, Krieger and Yekutieli correction for multiple comparisons.

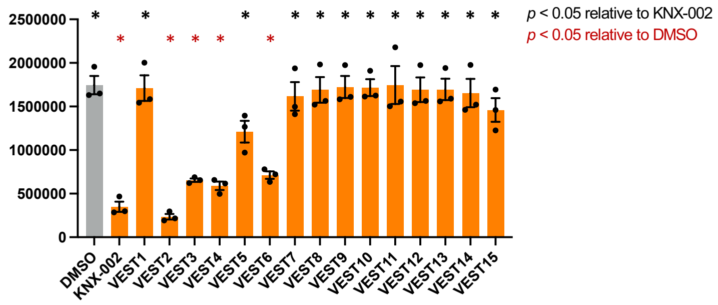


**<u>Suppl. Figure 3</u>: SAR analysis of KNX-002 analogs: TgMyoA ATPase activity.** Actin-activated ATPase activity of wild-type TgMyoA in the presence of DMSO (vehicle), KNX-002, or each of the 15 KNX-002 analogs shown in Figure 3. All compounds were used at 20 μM. ATPase activity is expressed as relative luminescence units (RLU). Each data point represents a single biological replicate composed of two technical replicates. Bars show the mean of the biological replicates ± SEM. RLU values were compared pairwise to KNX-002 (black asterisks) or to DMSO (red asterisks) by one-way ANOVA with Dunnett's test for multiple comparisons; asterisks indicate $p < 0.05$.

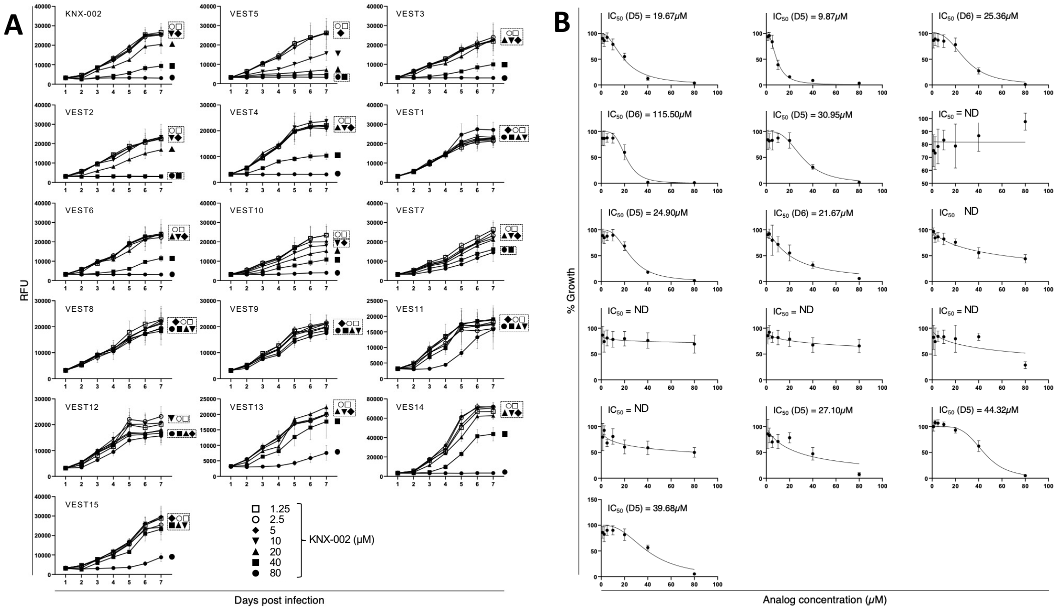


**<u>Suppl. Figure 4</u>: SAR analysis of KNX-002 analogs: parasite growth. (A)** tdTomato-expressing wild-type parasites were preincubated for 5 minutes with various concentrations (see key, bottom right) of KNX-002 or one of the 15 analogs shown in Figure 3 and then added to HFF cells on a 384-well plate. Fluorescence was measured daily over the next 7 days to quantify parasite growth. RFU = relative fluorescence units. **(B)** The data from either day 5 (D5) or day 6 (D6) of the growth assay shown in panel (A) were used to calculate the $IC_{50}$ for parasite growth; ND = insufficient inhibition from 0-80μM to determine an $IC_{50}$.

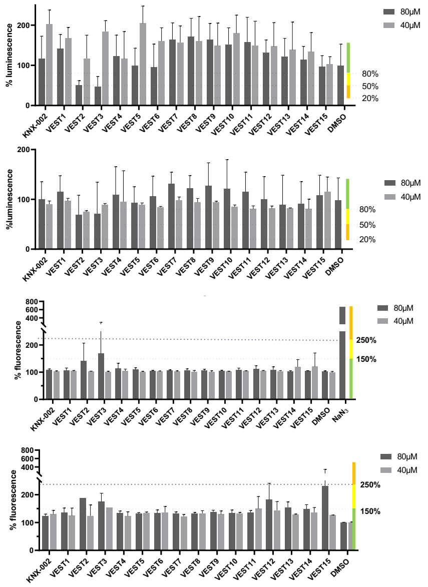


**<u>Suppl. Figure 5</u>: SAR analysis of KNX-002 analogs: host cell toxicity.** Subconfluent HFF and HepG2 cells were treated for 72 hr with DMSO (vehicle), KNX-002, or each of the 15 KNX-002 analogs shown in Figure 3, at a final concentration of either 40 µM (light gray bars) or 80 µM (dark gray bars). The treated cells were then subjected to both CellTox Green and CellTiter-Glo viability and toxicity assays, as indicated. Colored bars on the right indicate thresholds for negligible (green), mild (yellow) and moderate (orange) loss of viability (top 2 panels; CellTiter-Glo) and toxicity (bottom 2 panels; CellTox Green). These data are summarized in Figure 3.

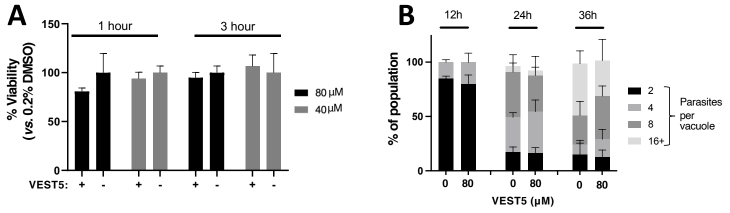


**<u>Suppl. Figure 6</u>: VEST5 is non-toxic to parasites. (A)** <u>Extracellular parasite toxicity</u>: Wild-type parasites were incubated at 37ºC in culture medium containing DMSO (-) or VEST5 (+) at either 40 µM (light grey) or 80 µM (dark grey) for the indicated times. Parasite viability was then determined using the Cell-TiterGlo® assay. Bars show the mean of two biological replicates ± SEM. **(B)** <u>Intracellular parasite toxicity</u>: Newly infected HFF cells were incubated at 37ºC in culture medium containing 0 (DMSO only) or 80 µM VEST5 for the indicated times. The cells were then fixed and processed for immunofluorescence using an antibody against the inner membrane complex marker IMC1, enabling the number of parasites in each of 50 vacuoles to be counted. Shading of the bars indicates number of parasites per vacuole (see key); the means ± SEM at each time point of two biological replicates are shown.

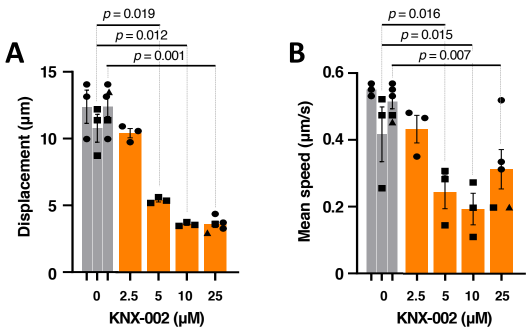


**Suppl. Figure 7: Dose-dependent effects of KNX-002 on parasite motility parameters. (A)** First-to-last point trajectory displacement and **(B)** Mean speed along the trajectory during a 60 sec 3D motility assay in the presence of the indicated concentrations of KNX-002. Note that the number of trajectories included in the displacement and speed measurements decreases with increasing concentration of compound due to the reduction in percent moving (See Figure 4C). Each data point represents a single biological replicate composed of three technical replicates. Sets of DMSO- and compound-treated parasite data captured on the same days, indicated by the similar symbol shapes, were compared by Student's one-tailed paired t-tests. Bars show the mean of the biological replicates ± SEM. Only the statistically significant differences ($p < 0.05$) are indicated.

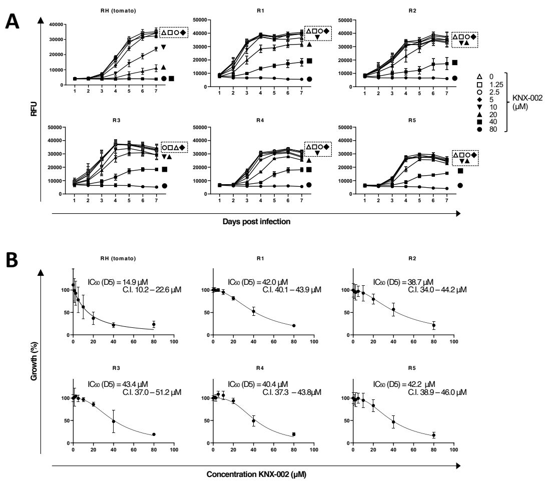


<u>Suppl. Figure 8</u>. **Growth assays of mutant parasites showing reduced sensitivity to KNX-002. (A)** tdTomato-expressing were mutagenized, selected in 40 µM KNX-002, and cloned as described in Methods. Each of the 26 clones recovered from two independent rounds of mutagenesis and selection was preincubated with various concentrations of KNX-002 for 5 minutes and then added to HFF cells on a 384-well plate. Fluorescence was measured daily over the next 7 days to quantify parasite growth. RFU = relative fluorescence units. **(B)** $IC_{50}$ curves corresponding to the growth assay data shown in panel (A), for the five clones that showed an $IC_{50}$ shift of > 2.5-fold compared to the un-mutagenized RH (tomato) parasites. The data shown are from three independent biological replicates, each consisting of 2-3 technical replicates at all time points. Vertical bars indicate SEM in panel (A) and 95% C.I. in panel (B). Clone R3 contains a T130A mutation in the gene encoding TgMyoA; none of the other four clones contain mutations in the genes encoding TgMyoA, TgMLC1, TgELC1 or TgELC2.

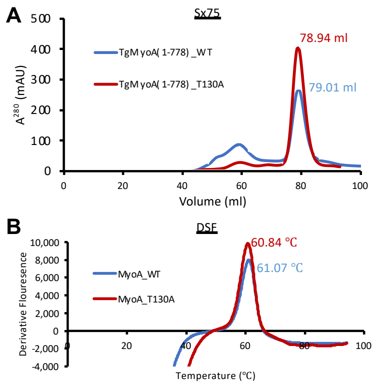


**Suppl. Figure 9. The T130A mutation does not affect the overall structure or stability of TgMyoA.** **(A)** SEC UV traces of TgMyoA motor domain, wild-type (blue) and T130A mutant (red), showing identical elution profiles. **(B)** Derivative fluorescence data from DSF experiment showing minimal changes in thermostability. Values are average of triplicate data.

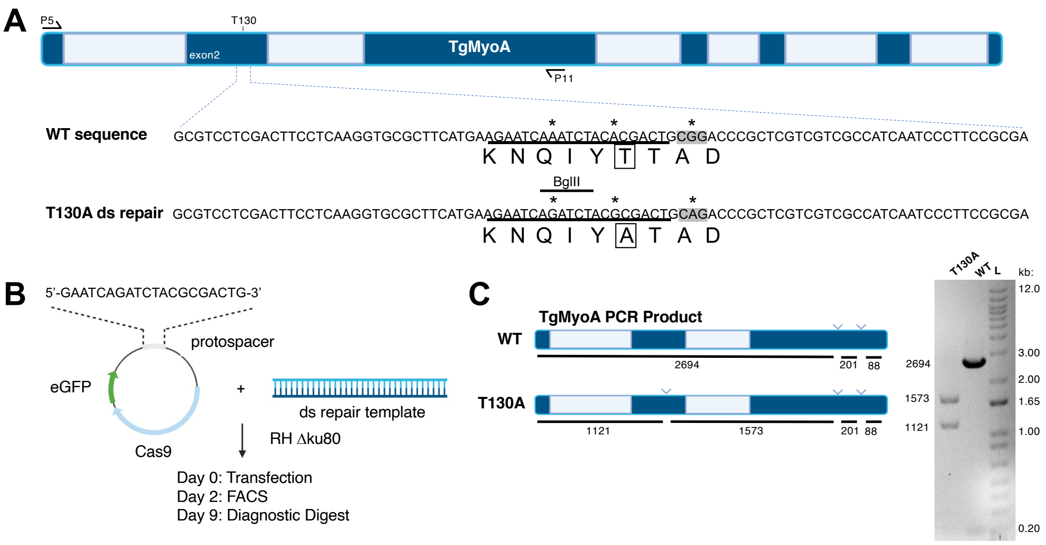


**<u>Suppl. Figure 10</u>: Generation of the TgMyoA T130A mutant parasite using CRISPR/Cas9-directed mutagenesis. (A)** Schematic of the *TgMyoA* gene (exons in dark blue, introns light blue) and the amino acids surrounding T130 (boxed) in exon 2. Mutations introduced by the double stranded (ds) repair template are denoted with asterisks (*): the leftmost mutation introduces a diagnostic BglII site, the middle mutation generates the T130A substitution (boxed), and the rightmost mutation ablates the PAM sequence. The guide sequence (protospacer) is underlined, and the PAM sequence is highlighted in gray. The location of primers P5 and P11 used to amplify a fragment from gDNA for diagnostic digestion (see Panel C) are shown. **(B)** Timeline showing transfection of RHΔku80Δhxgprt parasites with the Cas9 GFP guide vector and ds repair template on day 0, sorting of eGFP+ parasites on day 2, and diagnostic digest of gDNA from individual eGFP+ clones on day 9. **(C)** Left schematic shows predicted fragment sizes after BglII digestion of the PCR fragment generated by P5/P11, for parental (WT) and T130A parasites. Right panel shows the BglII restriction digest of a T130A positive clone (leftmost lane) and WT clone (middle lane); the 201 and 88 bp fragments are not resolved on this gel. Right lane = ladder, selected band sizes indicated in kb. Clones that showed the expected digestion pattern for the T130A substitution were verified by sequencing using primers P5-P13 (Table 1).

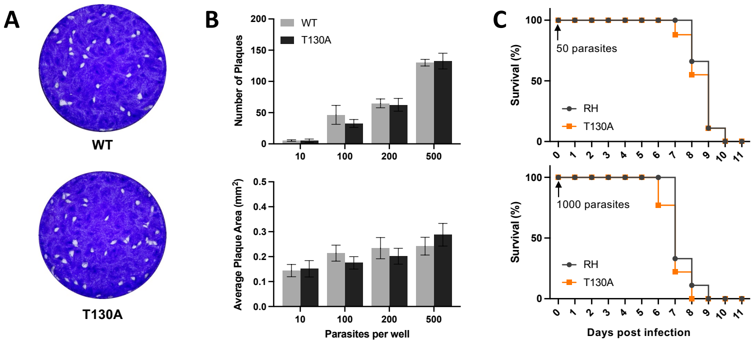


**<u>Suppl. Figure 11</u>: The TgMyoA T130A mutation has little to no impact on parasite growth *in vitro* or virulence *in vivo*. (A)** Representative images of a plaque assay using wildtype (top image; WT) and T130A mutant (bottom image) parasites, 7 days after inoculating confluent HFF monolayers with 100 parasites/well. The entire well (22.6mm diameter) from the 12-well plate is shown for each parasite line. **(B)** Total number of plaques per well and average plaque area in $mm^2$ of HFF monolayers inoculated with 10, 100, 200, and 500 parasites. The data represent the mean ± SEM from 4 independent biological replicates. There was no significant difference in plaque number or size between WT and T130A parasites at any of the inoculum sizes assayed. **(C)** Nine mice were infected on day 0 with 50 (top panel) or 1000 (bottom panel) wild-type RH (black circles) or T130A mutant parasites (orange squares). There was no significant difference in survival over the next 11 days between between mice infected with the same number of wildtype or T130A parasites (log-rank Mantel-Cox test).

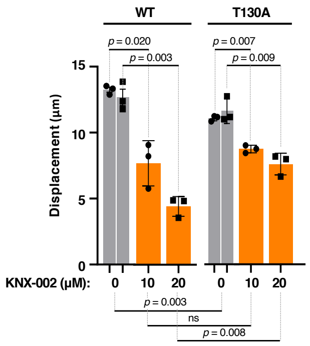


**<u>Suppl. Figure 12</u>: Parasite displacement in the 3D motility assay is less impacted by KNX-002 treatment in parasites expressing the TgMyoA T130A mutation, compared to wildtype.**
Displacement of wildtype (WT) and T130A mutant parasites during 80 sec of motility in Matrigel, in the presence of 0 (DMSO vehicle only), 10 or 20 µM KNX-002. The data were derived from the same set of experiments shown in Figure 7B-D. Each data point represents a single biological replicate composed of three technical replicates; three of the biological replicates were collected on the same three days (circles), and three of the biological replicates were collected on a different three days (squares). Bars show the mean of the biological replicates ± SEM. Sets of biological replicates using the same parasite line and collected on the same days were compared by Student's one-tailed paired t-tests (significance indicated above the graphs). Sets of biological replicates comparing different parasite lines were analyzed by Student's two-tailed unpaired t-tests (significance indicated below the graphs). ns = not significant.

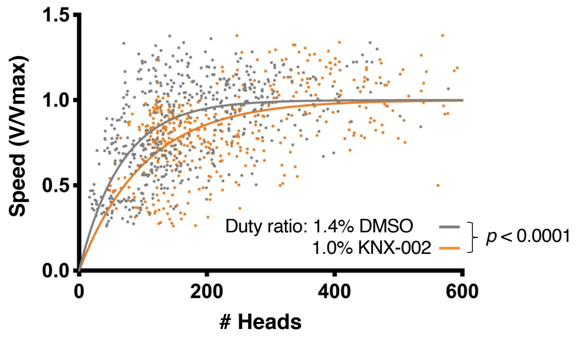


**<u>Suppl. Figure 13</u>: KNX-002 lowers the duty ratio of the TgMyoA motor.** Actin filament velocity (V/Vmax) was plotted as a function of the number of myosin heads capable of interacting with the actin filament. The duty ratios calculated from these data (see Methods) were significantly different for TgMyoA treated with DMSO (grey datapoints) *vs.* 10 μM KNX-002 (orange; Kolmogorov-Smirnov test, $p < 0.0001$).

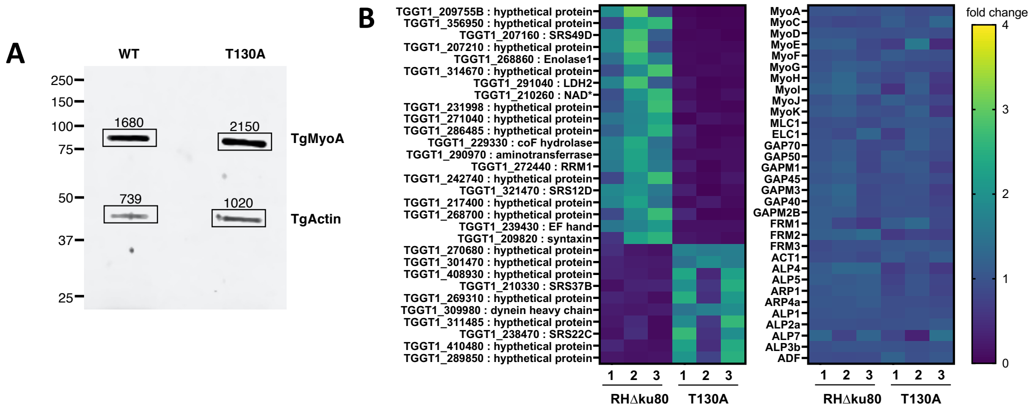


<u>Suppl. Figure 14</u>: **The T130A mutation neither upregulates TgMyoA nor changes the expression of other known motility-related genes. (A)** The expression level of TgMyoA in wild-type (WT) and T130A mutant parasites was compared via quantitative Western blotting. The blot was also probed for TgACT1 as a loading control; when normalized to the corresponding actin signals, the TgMyoA band intensities were within 7% of each other. **(B)** Left panel: Most differentially expressed genes between WT (RHΔku80) and T130A parasites (20 upregulated and 10 downregulated), after filtering for padj < 0.05 and FPKM > 0.1, and eliminating any genes that did not show the same pattern in all three replicates. Right panel: Analysis of differential expression of selected genes with the potential to compensate for loss of TgMyoA function, including other myosin motors, known MyoA interacting proteins, glideosome components, actin and actin regulatory factors.
